# A multi-looping chromatin signature predicts dysregulated gene expression in neurons with familial Alzheimer’s disease mutations

**DOI:** 10.1101/2024.02.27.582395

**Authors:** Harshini Chandrashekar, Zoltan Simandi, Heesun Choi, Han-Seul Ryu, Abraham J. Waldman, Alexandria Nikish, Srikar S. Muppidi, Wanfeng Gong, Dominik Paquet, Jennifer E. Phillips-Cremins

**Affiliations:** Department of Bioengineering, University of Pennsylvania, Philadelphia, PA; Epigenetics Institute, Perelman School of Medicine, University of Pennsylvania; Department of Genetics, Perelman School of Medicine, University of Pennsylvania; Institute for Stroke and Dementia Research (ISD), University Hospital, LMU Munich, 81377, Munich, Germany; Munich Cluster for Systems Neurology (SyNergy), Munich, Germany

## Abstract

Mammalian genomes fold into tens of thousands of long-range loops, but their functional role and physiologic relevance remain poorly understood. Here, using human post-mitotic neurons with rare familial Alzheimer’s disease (FAD) mutations, we identify hundreds of reproducibly dysregulated genes and thousands of miswired loops prior to amyloid accumulation and tau phosphorylation. Single loops do not predict expression changes; however, the severity and direction of change in mRNA levels and single-cell burst frequency strongly correlate with the number of FAD-gained or -lost promoter-enhancer loops. Classic architectural proteins CTCF and cohesin do not change occupancy in FAD-mutant neurons. Instead, we unexpectedly find TAATTA motifs amenable to binding by DLX homeodomain transcription factors and changing noncoding RNAPolII signal at FAD-dynamic promoter-enhancer loops. *DLX1/5/6* mRNA levels are strongly upregulated in FAD-mutant neurons coincident with a shift in excitatory-to-inhibitory gene expression and miswiring of multi-loops connecting enhancers to neural subtype genes. *DLX1* overexpression is sufficient for loop miswiring in wildtype neurons, including lost and gained loops at enhancers with tandem TAATTA arrays and singular TAATTA motifs, respectively. Our data uncover a genome structure-function relationship between multi-loop miswiring and dysregulated excitatory and inhibitory transcriptional programs during lineage commitment of human neurons homozygously-engineered with rare FAD mutations.

## Main

Alzheimer’s disease (AD) is a complex neurodegenerative disorder which presents clinically as the progressive deterioration of cognitive function and memory^1, 2^. Classic AD neuropathological hallmarks include the accumulation of β-amyloid (Aβ)-containing extracellular plaques and intracellular neurofibrillary tangles of the hyperphosphorylated microtubule-associated protein tau^3–5^. AD onset and progression also involves synaptic defects^6^, neuronal subtype-specific degeneration^7–10^, astrocyte/microglia activation^11^, and cell type-specific gene expression changes^12,13^. Molecular and cellular defects manifest decades before clinical symptoms^14^, but a definitive diagnosis is typically made post-mortem. Progress in ascertaining the temporal sequence of molecular, cellular, and tissue-scale pathological events has been slowed by the lack of tissue availability across early developmental stages prior to cognitive impairment.

Genetic factors play a critical and well-established role in AD, yet if and how genetic mutations modify epigenetic modifications and transcription to give rise to disease phenotypes remains poorly understood. The majority (90-95%) of cases are considered late-stage and sporadic (SAD) with an age of onset >65 years. Genome-wide association studies have uncovered more than 70 SAD-associated common single nucleotide variants^15–27^, and widespread transcriptional changes have been reported in post-mortem tissue from SAD patients^12, 13, 28^. Approximately 1-5% of cases are early-onset, familial (FAD), and driven by autosomal dominant, highly penetrant mutations in amyloid precursor protein (*APP*) and presenilin (*PSEN1/2*) genes^29–32^. In the case of FAD, chromatin and gene expression studies in post-mortem brain tissue are rare^33^. Alterations to chromatin accessibility, histone modifications, and gene expression have been reported genome-wide in transgenic mouse models^33^ and human induced pluripotent stem cell (iPSC)-derived neurons^34, 35^ with penetrant FAD mutations, but studies remain limited. Elucidating the mechanisms underlying the earliest stages of transcriptional dysregulation in human neurons with FAD mutations would shed light on the preclinical phase preceding early onset neurodegeneration.

Here, we sought to examine changes that occur in higher-order chromatin folding, epigenetic modifications to the linear genome, and gene expression due to rare FAD mutations during the earliest stages of human neural lineage commitment. To focus on neuron-restricted, cell autonomous molecular events, we used isogenic human induced pluripotent stem cell (iPSC) lines homozygously-engineered with mutations in the *APP* and *PSEN1* genes and differentiated into 2D monolayer post-mitotic neurons or 3D cortical organoids^36, 37^. We elected to focus on autosomal dominant APP(Swe) and PSEN1 (M146V) mutations because they are highly penetrant, of high effect size, and exhibit clear phenotypic changes early in brain development even in their heterozygous form^38, 39^. Employing homozygously engineered FAD-mutant neurons *in vitro* afforded us the opportunity to observe clear changes to the genome’s structure-function relationship without confounders of different genetic backgrounds, diluted signal-to-noise *in vivo*, or cellular subpopulation variation typical of post-mortem brain tissue. By integrating single-cell and ensemble genomics, computational biology, and transgene overexpression experiments, we uncover a link between multi-loop miswiring and dysregulated excitatory and inhibitory transcriptional programs during lineage commitment of human FAD-mutant neurons prior to formation of the classic neurodegeneration hallmarks Aβ plaques and hyperphosphorylated tau tangles. Our observations open up future studies dissecting the interplay among rare FAD mutations, amyloid peptides, and disruptions to the genome’s structure-function relationship.

## Results

### Isogenic human iPSCs engineered with rare FAD mutations exhibit amyloid accumulation and phosphorylated tau in long-term brain organoid but not neuron monolayer cultures

We previously reported the derivation and characterization of isogenic iPSCs engineered with homozygous APP KM670/671NL (“Swedish”, APP(Swe)) and PSEN1 (M146V) mutations using CRISPR/Cas9 (**Fig. 1a**, **Fig. S1a**)^34, 36^. These variants represent well-characterized, highly penetrant, rare mutations causing severe forms of FAD^38, 39^. Both APP(Swe) and PSEN1 (M146V) mutations are heterozygous in human populations. In the laboratory, homozygous iPSCs matched to isogenic controls have shown utility for the dissection of a mutation’s molecular effects due to clear phenotypic signal over noise^34, 36^. We verified that all iPSC lines (DIV0) were karyotypically normal (**Fig. S1b**), exhibited standard morphology indicative of the pluripotent cellular state (**Fig. S1c**), and homogeneously expressed known markers of pluripotency (e.g. OCT3/4 (also known as POU5F1) and SSEA4) (**Fig. S1c, Supplementary Methods**).

*In vitro* cellular models of AD can recapitulate different pathological hallmarks of disease onset and progression (**Fig. 1b**)^40, 41^. To focus on the early, neuron-restricted, cell autonomous molecular events in FAD, we first implemented a well-established, multi-stage protocol for neuronal differentiation of iPSCs in monolayer tissue culture (**Fig. 1c**)^36, 37^. Using immunostaining, we verified that early-stage iPSC-derived neural progenitor cells (eNPCs, DIV8) and late-stage iPSC-derived NPCs (lNPCs, DIV35) homogenously expressed the hallmark neural progenitor marker NESTIN and the forebrain marker FOXG1 (**Fig. S1d-e**). Moreover, DIV65 iPSC-neurons exhibited the expected post-mitotic spindle-like morphology and homogenously expressed pan-neural marker MAP2 (**Fig. S1f**). We confirmed that differentiation efficiency, homogeneity, and cell viability/proliferation were unchanged across all three genotypes as previously reported (**Fig. S1c-f)**^34, 36^.

To assess known differentiation markers^42^, we conducted bulk RNA-seq analysis in WT iPSCS (DIV0), eNPCs (DIV8), lNPCs (DIV35), and post-mitotic neurons (DIV65) (**Fig. S2a-b**). In DIV35 lNPCs, we observed increased levels of the FOXG1 forebrain marker and transcription factors known to be expressed in the neocortex (e.g. PAX6, LHX2). We also observed that ventral telencephalon (e.g. GSX2, NKX2-1) and radial glia (e.g. VIM, HES1, FABP7/BLBP) markers peak at DIV35 and remain lowly expressed in post-mitotic neurons as expected. Finally, during the transition from DIV35 lNPC to DIV65 post-mitotic neurons, we observed increased expression of immature and mature neuronal markers, including: TUBB3, DCX, DLG4/PSD95, SYP, and GAP43. These data together confirmed that we can create homogeneous populations of four stages of cell types representing early neurodevelopment using isogenically-matched, karyotypically normal WT, APP(Swe), and PSEN1(M146V) iPSCs.

**Figure 1:**
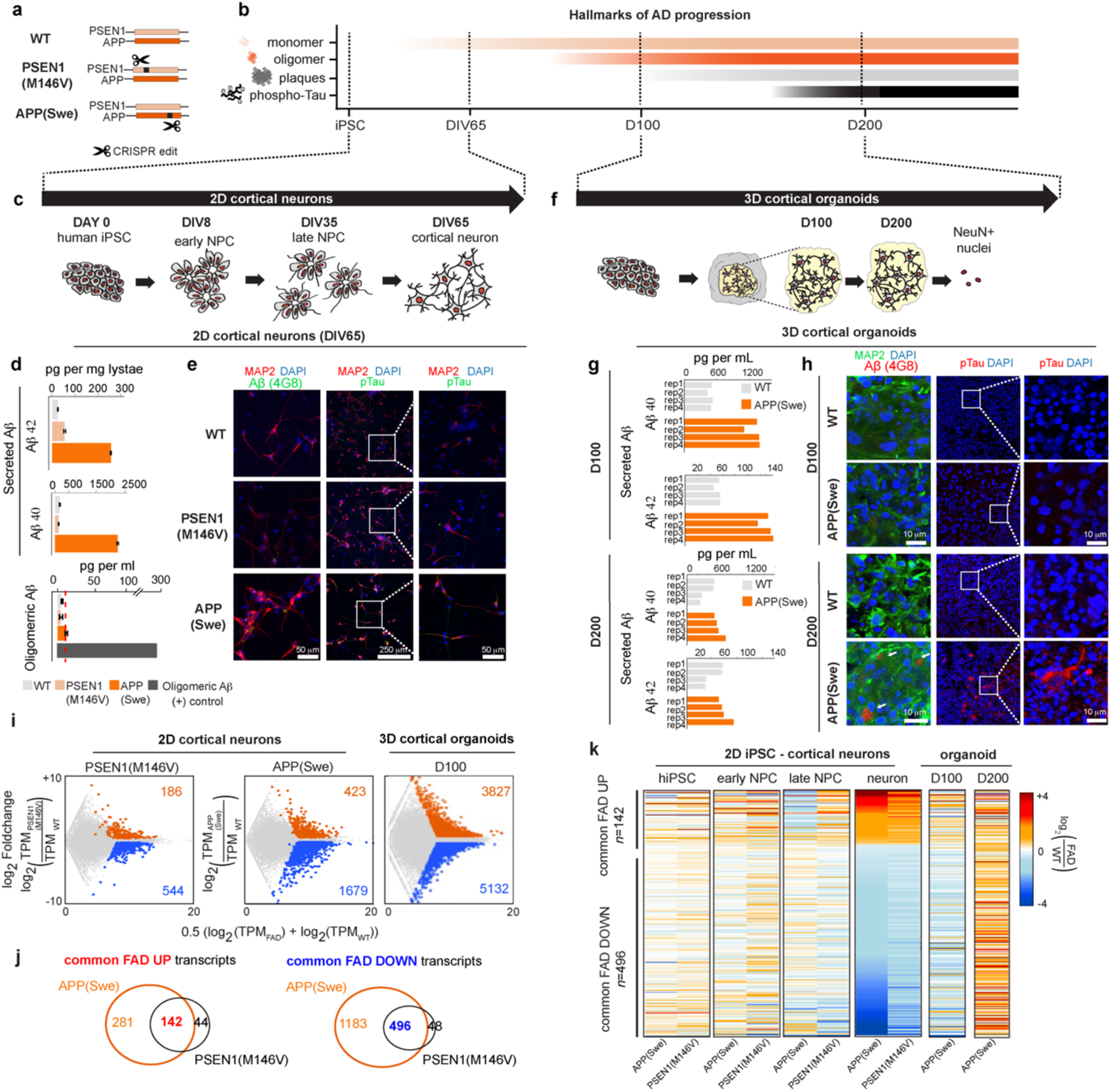
Human iPSC-derived neurons with rare, autosomal dominant FAD mutations exhibit widespread transcriptional dysregulation prior to Aβ accumulation and phosphorylated tau. (**a-b**) Schematics representing (**a**) CRISPR/Cas9-mediated introduction of homozygous APP(Swe) or PSEN1(M146V) mutations in human iPSCs and (**b**) hallmark protein markers of FAD, (**c**) 2D monolayer model of chemically induced iPSC differentiation into neurons, (**d**) genotype-dependent changes in Aβ40, Aβ42, and oligomeric Aβ detected from cell supernatant in FAD and WT DIV65 iPSC-derived neurons. Red line, assay detection limit, (**e**) Representative immunofluorescence images from WT and FAD iPSC-derived neurons (DIV65) stained for neuron marker MAP2 (red), DAPI (blue), Aβ (green, 4G8 antibody), and phosphorylated tau (green, AT8 antibody). Scale bars, 250μm and 50μm. (**f**) human iPSC-derived cortical organoid model, (**g**) genotype-dependent changes in Aβ40, Aβ42, and oligomeric Aβ in DIV100 and DIV200 iPSC-derived cortical organoids, (**h**) confocal images in DIV100 and DIV200 WT and APP(Swe) organoids stained for mature neuron marker (MAP2 (green)), Aβ accumulation (red), and phosphorylated tau (red). Scale bar, 10μm. (**i**) M-A plot of RNA-seq data from iPSC-derived DIV65 neurons and DIV100 organoids from WT, PSEN1(M146V), and APP(Swe) genotypes. (**j**) Venn diagrams of (*left*) significantly upregulated mRNA transcripts in human iPSC-derived neurons with APP(Swe) or PSEN1(M146V) genotypes compared to WT: “common FAD UP” (*n=142*) and (*right*) significantly downregulated mRNA transcripts in human iPSC-derived neurons with APP(Swe) or PSEN1(M146V) genotypes compared to WT: “common FAD DOWN” (*n= 496*), (**k**) Heatmap of fold-change mRNA levels for common FAD UP and common FAD DOWN transcripts across iPSCs, early NPCs, late NPCs, DIV65 post-mitotic neurons, and DIV100/DIV200 organoids.

We next set out to understand the extent to which iPSC-neurons in monolayer culture produce pathologic hallmarks of AD, including increased levels of Aβ40 and Aβ42 peptides, Aβ oligomers, extracellular Aβ accumulation, and intracellular phosphorylated tau^14, 43–45^. Consistent with literature^46–50^ and previous reports robustly differentiating neurons from our iPSC lines^34, 36^, we observed more than a ten-fold increase in Aβ40 and Aβ42 levels in APP(Swe)-neurons (**Fig. 1d**), as well as a two-fold increase in the Aβ42:40 ratio in PSEN1(M146V)-neurons (**Fig. S3a**), compared to isogenic WT iPSC-neurons. Increases in Aβ40 and Aβ42 levels in FAD genotypes occurred in both lNPC and neurons (**Fig. S3a, Table S1**). As previously established^51, 52^, we observed negligible or undetectable levels of Aβ oligomers (**Fig. 1d, Fig. S3a, Table S1**), extracellular Aβ accumulation, or tau phosphorylation (**Fig. 1e, Fig. S3b-e, Table S1**), confirming that iPSC-neurons in monolayer with endogenous Aβ40/42 peptide generation exhibit negligible evidence of advanced protein hallmarks of FAD. Thus, we verify previous reports^36, 37^ that iPSC-neurons with homozygous FAD mutations in monolayer culture represent a robust model of an early stage of AD onset marked by increase in Aβ monomeric species but prior to Aβ oligomerization, extracellular accumulation of Aβ, or pathological tau phosphorylation.

To confirm that our model system has the capability of accumulating Aβ and/or phosphorylated tau in long-term 3D culture, we generated cortical organoids with established methods^53, 54^ (**Fig. 1f, Fig. S4a**). We focused on the APP(Swe) FAD genotype because it is well documented to cause Aβ aggregates in early-onset FAD patients with severe cognitive defects^55,56^. We observed SOX2-positive rosette-like structures indicative of the NPC subtypes of neuroepithelial stem cells and ventricular radial glial cells in similar morphology and levels with both WT and APP(Swe) genotypes at DIV50 (**Fig. S4b**). Pan-neural (e.g. MAP2) and cortical layer markers (e.g. CTIP2 and TBR1) were detected in both genotypes by DIV100 as previously reported^53, 54^ (**Figs. S4c-d**). By DIV200, CTIP/TBR1+ neurons had organized into a tissue structure indicating potential multi-laminar cortical layer organization. Cells positive for astrocyte marker GFAP had increased substantially (**Figs. S4e-f).** We did not observe major differences in cortical layer structure or neural density between genotypes at DIV100 or DIV200 (**Fig. S4c-f**). Using immunohistochemistry and biochemistry experiments, we observed increased Aβ40 and Aβ42 in APP(Swe) organoids (**Figs. 1g-h, S4g-i).** By DIV200, staining patterns consistent with accumulated Aβ and phosphorylated tau were detectable in APP(Swe) organoids (**Fig. 1h, Fig. S4g-h, Movie S1, Table S1).** Together, these data reveal that biochemical and immunofluorescence staining patterns consistent with Aβ accumulation and phosphorylated tau can be detected in human APP(Swe) iPSC-derived DIV200 organoids, illustrating the need of a 3D tissue environment and long timescales to form advanced pathologic AD hallmarks.

#### FAD-mutant iPSC-neurons in 2D monolayer culture exhibit widespread transcriptional dysregulation prior to Aβ accumulation and tau phosphorylation

We next used RNA-seq to understand the extent to which mRNA levels change due to rare FAD mutations. We observed only minor changes in gene expression due to rare FAD mutations in iPSC, eNPC, or lNPC cell types (**Fig. S5a-b**). By contrast, we observed a substantial increase in the number of dysregulated genes due to FAD mutations in post-mitotic neurons (**Fig. 1i, Fig. S5a-b, Table S2**). The APP(Swe) FAD line showed a more severe expression phenotype (n=1,679 downregulated and n=423 upregulated transcripts compared to WT, respectively) than the PSEN(M146V) FAD line (n=544 downregulated and n=186 upregulated transcripts compared to WT, respectively) (**Fig. 1i, Fig. S5b**). Importantly, although the two FAD lines have different rare variants, we found 638 common differentially expressed transcripts that were dysregulated in both the FAD genotypes in post-mitotic neurons compared to isogenic WT neurons, hereafter termed “common FAD DOWN” (n=496) and “common FAD UP” (n=142) (**Fig. 1j, Table S3**). Together these data demonstrate widespread and reproducible dysregulation in mRNA levels due to single FAD mutations in post-mitotic neurons in DIV65 monolayer culture.

Our monolayer 2D and 3D organoid models afforded us the opportunity to ensure that the gene expression patterns were not an artifact of 2D monolayer culture. We performed RNA-seq on NeuN+ sorted nuclei from DIV100 (negligible amyloid accumulation) and DIV200 (amyloid accumulation + hyperphosphorylated tau) organoids to compare with monolayer DIV65 neurons (**Fig. S5C, S6, Table S4)**. We found that the reproducibly downregulated and upregulated transcripts due to FAD mutations in DIV65 monolayer neurons (common FAD DOWN, common FAD UP) also showed similar trends for the same genes in NeuN+ sorted neurons from DIV100 organoids (**Fig. 1k, Fig. S5c)**. In NeuN+ neurons from DIV200 organoids, the gene expression changes had almost no resemblance to DIV100 organoids or DIV65 monolayer (**Fig. 1k, Fig. S5c**), which is consistent with recent reports of a second wave of dysregulated gene expression coincident with accumulation of Aβ aggregates, tau tangles, and severe disruption of cell viability^12^. Our data confirm that the dysregulated genes common to DIV65 neurons with APP(Swe) and PSEN mutations were similarly misexpressed in DIV100 organoids and thus not an artifact of 2D monolayer culture.

To assess the potential physiologic relevance of the dysregulated genes in FAD-mutant neurons, we conducted gene ontology (GO) analysis. The genes that are “common FAD UP” (n=142) represent GO terms of GABAergic synaptic transmission, neural lineage commitment, axon development, and neuronal projections and patterning (**Fig. S5d, left**). The genes that are “common FAD DOWN” (n=496) exhibit GO terms for common cellular functions such as metabolism, cell adhesion, and membrane localization of proteins (**Fig. S5d, right**). Genes commonly dysregulated in both DIV65 monolayer and DIV100 organoids represent cell adhesion, neural lineage commitment, neuron projections, and axon development ontology (**Fig. S5e**). Moreover, dysregulated genes unique only to DIV100 organoids encode synapse maturation indicative of the 3D environment provided by organoids (**Fig. S5e**). Together, these analyses underscore the physiologic relevance of the dysregulated genes in FAD-neurons and highlight the importance of further studying the chromatin mechanisms governing their misexpression in early stages of neurodevelopment and degeneration.

#### Transcriptional dysregulation in FAD neurons is not associated with changes in chromatin accessibility at gene promoters

As a first step toward elucidating the mechanisms driving transcriptional dysregulation due to FAD mutations, we focused our study on our pure population of DIV65 neurons with no evidence of confounds from amyloid accumulation or phosphorylated tau. We first measured chromatin accessibility genome-wide in WT, PSEN1(M146V), and APP(Swe) iPSC-derived neurons using ATAC-seq (**Table S5**). We unexpectedly observed that the promoters of common FAD Down and common FAD Up genes showed no change in chromatin accessibility in neurons (**Fig. 2a-b, Fig. S7a**). Although a substantial proportion of ATAC-seq peaks changed between APP(Swe) or PSEN1(M146V) FAD neurons compared to WT (**Fig. S7b-c, Table S6),** nearly all were located distal from TSSs (**Fig. 2c, Fig. S7d**). These observations suggest that transcriptional dysregulation in FAD neurons cannot be explained by chromatin regulatory mechanisms at the core promoter.

#### Long-range chromatin looping interactions are miswired in neurons with FAD mutations

Distal non-coding regulatory elements such as enhancers can regulate their target genes via long-range looping interactions^57–59^. Therefore, we next investigated the extent to which topologically associating domains (TADs)^60–63^, subTADs^64^, and loops^65^ were altered genome-wide. We created *in situ* Hi-C libraries sequenced to 10 kb resolution with ∼450 million uniquely mapped interactions across both replicates in WT, APP(Swe), and PSEN1(M146V) DIV65 iPSC-neurons. Neurons from all three genotypes achieved the cis/trans read distribution expected for high-quality libraries and exhibited high concordance between biological replicates (**Fig. S8a-b, Table S7)**. Using our graph theory-based method 3DNetMod^66–68^, we identified similar numbers of 14,620, 14,873 and 14,881 TADs/subTADs in WT, APP(Swe), and PSEN(M146) post-mitotic neurons, respectively (**Table S8, Supplementary Methods**). Visual inspection and quantitative analysis of the genome-wide insulation score further confirmed that TAD/subTAD boundaries were in large part unchanged across WT and FAD genotypes (**Fig. S8c-e**). These data indicate that TAD and subTAD boundaries are generally unchanged due to FAD mutations in post-mitotic neurons.

We previously developed statistical methods to identify dot-like features in Hi-C data characteristic of bona fide long-range chromatin loops exhibiting significantly higher interaction frequency compared to the local TAD/subTAD structure^65, 67–71^ (**Table S9, Supplementary Methods**). We previously developed a negative binomial likelihood ratio test – the 3DeFDR algorithm – to find statistically significant differential loops across biological conditions^70, 71^. Because the most severe changes in gene expression were observed in the APP(Swe) condition, we focused our loop analysis on APP(Swe) and WT neuron Hi-C. We applied 3DeFDR to identify 1,680 APP(Swe)-specific loops, 3,346 WT-specific loops, and 29,527 genotype-invariant loops in post-mitotic neurons (**Fig. 2d-f, Table S10**). We stratified FAD-dynamic and genotype-invariant loops into those connecting promoters to distal promoters, promoters to non-promoter regions, and non-promoter regions to other distal non-promoter regions **(Fig. 2g-h, Fig. S7e-g, Table S11**). We observed moderate changes in chromatin accessibility only at distal non-promoter anchors **(Fig. 2i, Fig. S7i)**. Negligible changes in chromatin accessibility were apparent at promoters whether they engage in loops with non-promoter regions (**Fig. 2i, Fig. S7i)**, distal promoters (**Fig. S7f-h)**, or do not loop at all (**Fig. 2j**). Together, our data indicate that thousands of loops change in FAD-mutant neurons. Chromatin accessibility exhibits dynamic changes in FAD-mutant neurons primarily at the non-coding anchor of promoter-enhancer loops.

**Figure 2.**
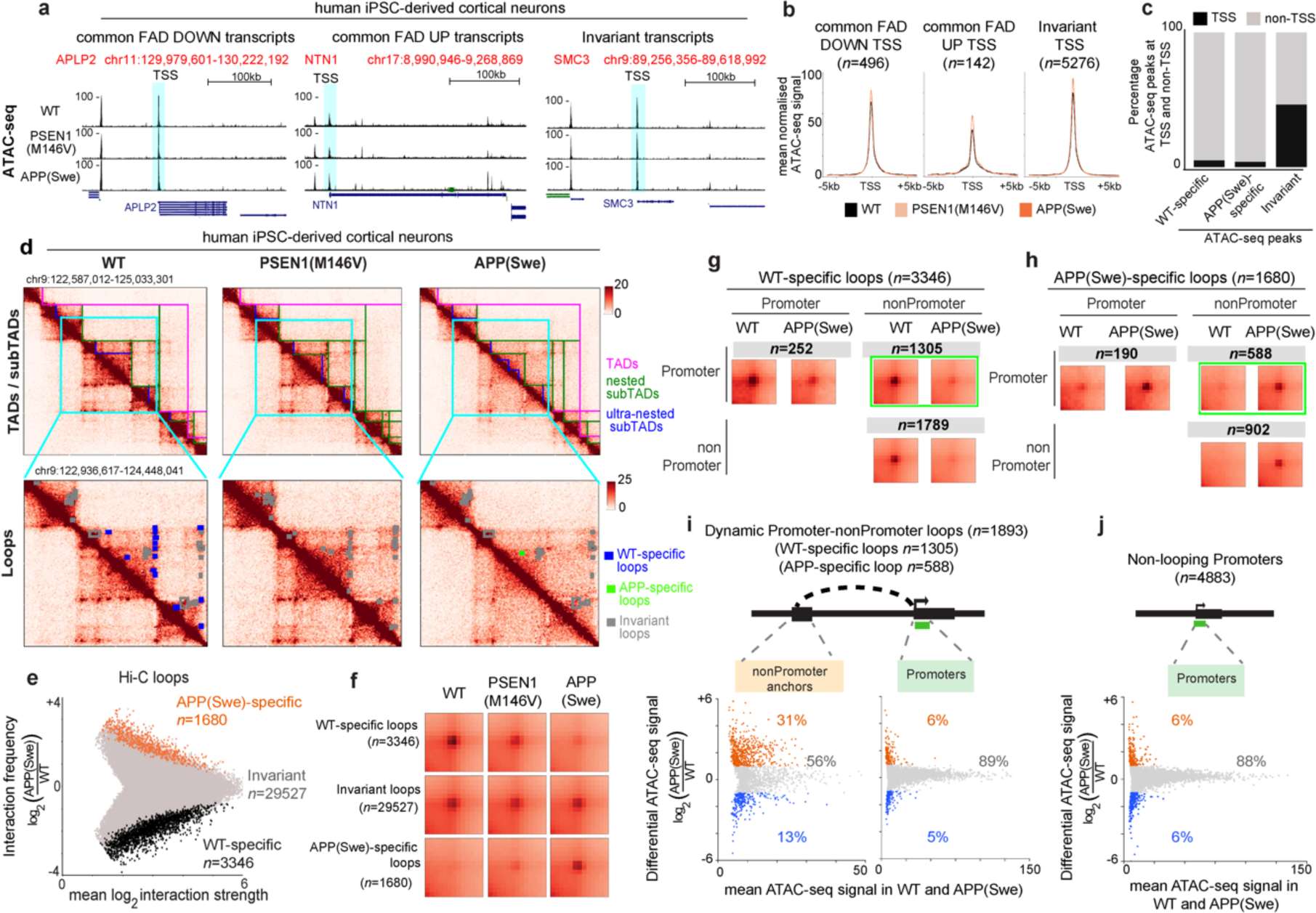
Chromatin loops are miswired in neurons with FAD mutations. **(a)** ATAC-seq tracks in WT, PSEN1(M146V), and APP(Swe) iPSC-derived neurons at genes that are common FAD downregulated (*APLP2*), common FAD upregulated (*NTN1*), and invariantly expressed (*SMC3*) as defined in (Fig. 1j**, Table S3**). Promoters highlighted by cyan box. **(b)** Aggregate ATAC-seq peak signal at TSSs ± 5kb at common FAD downregulated (*n=*496), common FAD upregulated (*n=*142), and invariantly expressed genes (*n=*5276). **(c)** WT-specific, APP(Swe)-specific, and genotype-invariant ATAC-seq peaks at TSS ± 2kb (black) and non-TSS (grey) regions genome-wide. **(d)** Hi-C heatmaps at a representative 2.5 Mb region in WT, APP(Swe) and PSEN1(M146V) iPSC-neurons across TADs (magenta lines), subTADs (green, blue lines), and loops (green dots). Bin size, 10 kb, hg38 genome. Cyan arrows highlight miswired loops. **(e-f)** M-A plot and APA analysis of cell type-specific and genotype-invariant loops between APP(Swe) and WT post-mitotic neurons (**Supplementary Methods**). **(g)** Aggregate peak analysis of looping interaction frequency signal in WT and APP(Swe) iPSC-neurons across all WT-specific loops classified into Promoter-Promoter (*n=*252), Promoter-nonPromoter (*n=*1305), and nonPromoter-nonPromoter classes (*n=*1789), **(h)** Aggregate peak analysis of looping interaction frequency signal in WT and APP(Swe) iPSC-neurons across all APP(Swe)-specific loops that are classified into Promoter-Promoter (*n=*190), Promoter-nonPromoter (*n=*588), and nonPromoter-nonPromoter classes (*n=*902). **(i-j)** M-A plot of differential ATAC-seq signal between WT and APP(Swe) iPSC-derived neurons at **(i)** the nonPromoter anchors (*left*) and at promoters (*right*) for the FAD dynamic Promoter-nonPromoter loops and **(j)** the TSS ± 2kb promoters of all non-looping genes.

#### The severity and direction of changes to mRNA levels and single-cell burst frequency strongly correlates with the number of dynamic loops per gene

A fundamentally important unanswered question is the extent to which loops control gene expression. Recent studies have called the functional role for loops into question based on the minimal effect of acute loop disruption on the maintenance of gene expression^72, 73^. Studies to date have focused on dividing cell lines, and it is unknown how loops relate to gene expression in post-mitotic neurons or when transcriptional programs are altered due to changes in cellular state.

To dissect the relationship between chromatin loop miswiring and gene expression changes in our neuronal FAD model, we first classified promoters by their engagement in WT-specific loops (FAD-lost, Class 1), APP(Swe)-specific loops (FAD-gained, Class 2), or only genotype-invariant loops (Class 3) with non-coding (NonPromoter) regions of the genome, as well as those that are not looping (Class 4) (**Fig. 3a, Table S12**). Importantly, presence and changes to single loops did not correlate with gene expression dysregulation. We observed a dynamic range of up to 32-fold (log_2_(32)=5) increased or decreased mRNA levels in APP(Swe) compared to WT neurons no matter if a promoter engaged in a single invariant loop, a single dynamic loop, or no loop at all (**Fig. 3b**). We instead observed a striking pattern where the number of FAD-dynamic loops per promoter was strongly predictive of a consistent direction and degree of gene expression dysregulation (**Fig. 3b, S9a-b).** Genes with multiple disease-lost Promoter-to-NonPromoter loops were consistently downregulated by 9-16-fold in FAD-mutant neurons in a manner commensurate with the number of lost loops (**Fig. 3b, S9a-b)**. Similarly, genes engaged in multiple disease-gained Promoter-to-NonPromoter loops were consistently upregulated in FAD-mutant neurons compared to WT (**Fig. 3b, S9a-b)**. Our results revealed that single loop changes were poor predictors of gene expression changes, however, multiple gained or lost loops between promoters and their distal non-coding region targets exhibited consistently severe changes in mRNA levels in post-mitotic FAD neurons.

**Figure 3:**
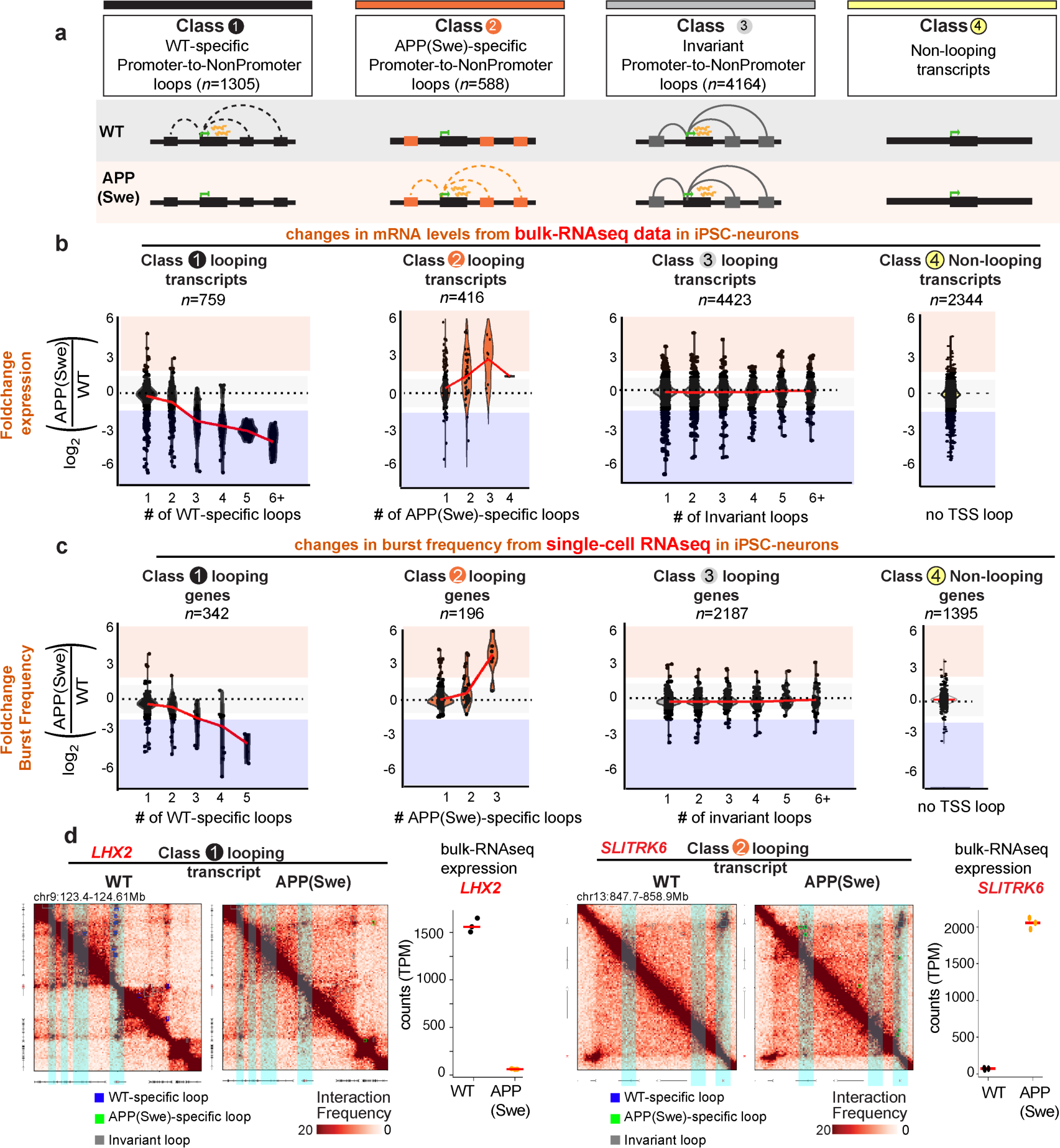
The severity and direction of fold-change in mRNA levels and single-cell burst frequency strongly correlates with the number of changing Promoter-to-NonPromoter loops in FAD-mutant neurons. **(a)** Schematic depicting genes classified by their connection in WT-specific (FAD-lost, Class 1), APP(Swe)-specific (FAD-gained, Class 2), and genotype-invariant (Class 3) Promoter-to-NonPromoter loops as well as not looping (Class 4). **(b-c)** Strip plot of the fold change in **(b)** bulk RNA-seq mRNA levels and **(c)** single-cell RNA-seq burst frequency stratified by the number of WT-specific, APP(Swe)-specific, or genotype-invariant Promoter-to-NonPromoter loops formed per gene. **(d)** Hi-C interaction frequency heatmaps and normalized mRNA levels from bulk-RNAseq for the *LHX2* and *SLITRK6* genes. *LHX2* represents a Class 1 multi-looper with severe downregulated mRNA levels and *SLIRTK6* represents a Class 2 multi-looper with severe upregulation of mRNA levels in FAD neurons. Blue circles, WT-specific loops. Green circles, APP(Swe)-specific loops. Grey circles, genotype-invariant loops.

Transcription occurs in pulses (or ‘bursts’) of mRNA production over time, alternating between active “on” states and inactive “off” states. In single cells, the frequency of transcriptional bursts can be calculated as the metric ‘burst frequency’. To assess the role for multi-looping genes on transcriptional burst frequency, we generated single-cell RNA-seq (scRNA-seq) data in WT and FAD neurons. We first confirmed with independently generated scRNA-seq data that the number of WT-specific or APP(Swe)-specific loops strongly correlated with the degree of fold-change in mRNA levels (**Fig. S9c**). We further uncovered a strong correlation between the number of FAD-dynamic loops and transcriptional burst frequency in single cells (**Fig. 3c, Table S13**). The number of WT-specific and APP(Swe)-specific loops strongly correlated with consistently decreased and increased transcriptional burst frequency, respectively. Burst size showed no correlation with loop changes (**Fig. S9d**). The striking multi-loop link to single-cell changes in mRNA levels and burst frequency is unique to Promoter-to-NonPromoter loops, as Promoter-to-Promoter loops did not correlate with gene expression dysregulation in WT and APP(Swe) human iPSC-derived neurons (**Fig. S10a-e**). Locus-specific examples confirmed our quantitative genome-wide findings. Multi-looping genes *LHX2* and *SLITRK6* exhibit up to a 500-fold change in mRNA levels coincident to the number of WT-specific and APP(Swe)-specific loops, respectively (**Fig. 3d**). Together, our data demonstrate that the number of disrupted or newly formed loops connecting promoter and distal non-promoter regions are a strong predictor of the severity and direction of fold change in mRNA levels and burst frequency in FAD neurons.

#### Architectural proteins CTCF and cohesin exhibit unchanged occupancy at the genomic anchors of WT-specific and APP(Swe)-specific promoter-enhancer multi-loops

We next set out to understand the mechanism governing the formation of WT-specific and APP(Swe)-specific multi-loops. Understanding this mechanism is important given that such loops connect physiologically relevant genes dysregulated in FAD neurons.

**Figure 4.**
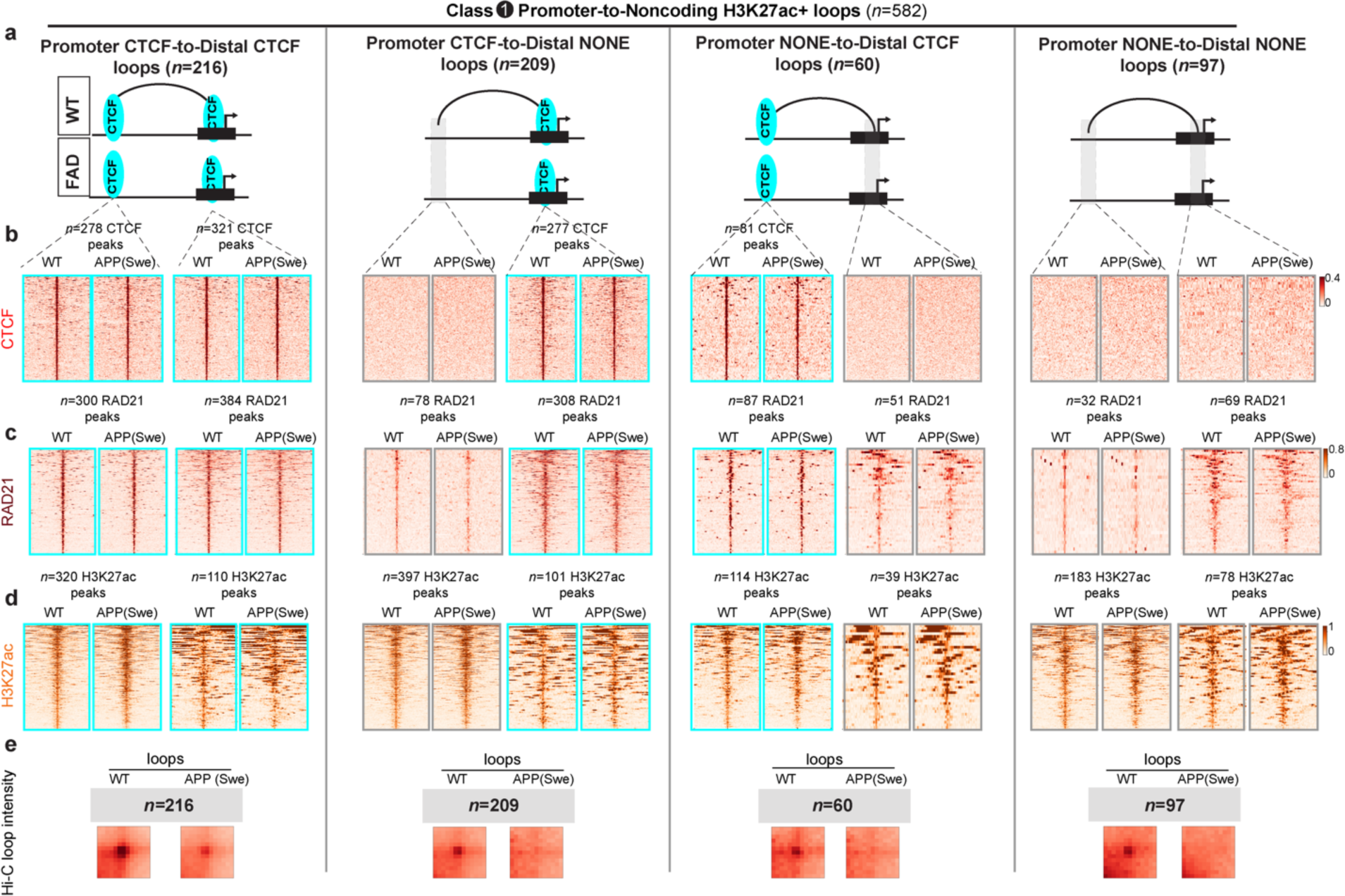
Promoter-enhancer loops lost in FAD neurons are anchored by unchanging occupancy of the architectural proteins CTCF and cohesin. (**a**) Stratification of WT-specific Promoter-NoncodingH3K27ac-positive(+) loops into those anchored by (column 1) Promoter CTCF-to-Distal CTCF, (column 2) Promoter CTCF-to-Distal NONE, (column 3) Promoter NONE-to-Distal CTCF, and (column 4) Promoter NONE-to-Distal NONE. (**b-d**) ChIP-seq heatmaps of (**b**) CTCF occupancy, (**c**) RAD21 occupancy, and (**d**) H3K27ac signal in WT and APP(Swe) iPSC-derived neurons. (**e**) Aggregate peak analysis of Hi-C interaction frequency at WT-specific loops in WT and APP(Swe) iPSC-derived neurons.

We performed ChIP-seq for the known architectural proteins CTCF and a subunit of the cohesin complex RAD21 (**Table S14**). We focused on WT-specific and APP(Swe)-specific loops connecting promoters to non-coding putative enhancer regions marked by H3K27ac signal as assayed by ChIP-seq (**Fig. 4a, Fig. S11a-c, Table S15**). We further sub-stratified the WT-specific and APP(Swe)-specific promoter-enhancer loops into those with (i) CTCF occupancy on both anchors, (ii) CTCF occupancy at the promoter anchor, (iii) CTCF occupancy at the non-coding enhancer anchor, and (iv) no CTCF occupancy on either anchor (**Fig. 4a, Fig. S11c, Table S11**). Unexpectedly, we observed no change in CTCF occupancy at either the promoter or distal non-coding loop anchor between WT and APP(Swe) genotypes (**Fig. 4b, Fig. S11d**). We also observed strong binding of the cohesin subunit RAD21 at both noncoding and promoter anchors whether CTCF was present or not (**Fig. 4c, Fig. S11e**). RAD21 occupancy was also unchanged across genotypes. Altogether, our data unexpectedly reveal that promoter-enhancer loops markedly change interaction frequency in FAD neurons (**Fig. 4e, Fig. S11g**) independent from the known mechanisms of altered architectural protein occupancy.

#### RNA Polymerase II signal at non-coding H3K27ac+ enhancers changes substantially and is strongly correlated with FAD gained and lost promoter-enhancer loops

We next set out to understand if aspects of noncoding enhancers were changing in FAD dynamic loops. We first examined the classic enhancer histone modification H3K27ac and found it unchanged between WT and APP(Swe) neurons (**Fig. 4d, Fig. S11f**). We also assessed chromatin accessibility at distal noncoding putative enhancers and did not find changes in ATAC-seq signal that were consistent with loop loss or gain (**Fig. 5a-b, S12a-b**). Thus, we did not see correlated loss in chromatin accessibility or H3K27ac at either anchor for WT-specific or APP(Swe)-specific promoter-enhancer loops in FAD-mutant neurons.

What then changes at FAD dynamic loops? We recently reported that RNA Polymerase II (RNAPII) occupancy at promoters correlates with loops that are gained during lineage commitment of human iPSCs to NPCs and NPCs to neurons^74^. Using ChIP-seq, we unexpectedly observed a strong loss of RNAPII signal in FAD-mutant neurons at H3K27ac-positive enhancers anchoring WT-specific promoter-enhancer loops (**Fig. 5c, Table S14**). Consistent with this pattern, we also observed strong gain of RNAPII signal at H3K27ac-positive enhancers in APP(Swe)-specific gained promoter-enhancer loops (**Fig. S12c, Table S14**). We confirmed that RNAPII occupancy did not exhibit such changes between WT and APP(Swe) neurons at non-looping H3K27ac+ enhancers or non-looping promoters (**Fig. S13a-f**). Together, our data reveal that RNAPII signal at putative enhancers exhibits significant changes in occupancy commensurate with the loss and gain of promoter-enhancer loops in FAD neurons.

#### An unexpected role for DLX homeodomain transcription factors anchoring miswired promoter-enhancer chromatin loops in FAD-mutant neurons

To identify candidate factors contributing to changes in RNAPII and loop interaction frequency, we next performed motif enrichment analysis at the ATAC-seq peaks at distal non-coding H3K27ac+ enhancer regions (**Supplementary Methods**). At WT-specific promoter-enhancer loops, we found a striking enrichment of multiple homeodomain transcription factor (TF) TAATTA motifs under the WT-specific lost RNAPII signal at enhancers (**Fig. 5d**). Moreover, at APP(Swe)-specific promoter-enhancer loops, we also observed slight enrichment of single TAATTA motifs under the APP(Swe)-specific gained RNAPII signal at enhancers (**Fig. S12d**).

Homeodomain TFs represent a family of evolutionarily conserved proteins that bind DNA and have widespread roles in regulating development^75^. We queried mRNA levels in the RNA-seq data from our model system to identify candidate homeodomain TFs which might differentially bind to the TAATTA motif, and observed that *LHX2*, *LHX9*, and *EMX2* mRNA levels were high in WT and silenced in APP(Swe) neurons (**Fig. 5e**). We further discovered that *DLX1, DLX5*, and *DLX6* mRNA levels exhibited negligible expression in WT and were strongly upregulated in APP(Swe)-mutant neurons (**Fig. 5f**). These data provide the initial observation that marked changes in mRNA levels of multiple homeodomain TFs occur coincident with loop changes between WT and APP(Swe) genotypes.

We elected to focus on DLX TFs because they are master regulators of GABAergic interneuron lineage commitment and play critical roles in gene expression regulation in the developing brain^75, 76^. A major technical challenge is that custom ChIP assays for DLX proteins are unattainable due to the lack of availability of epitope-specific ChIP-grade antibodies. For this reason, we re-analyzed published data representing ChIP-seq with custom-made, validated DLX1 and DLX5 antibodies in micro-dissected brain tissue at stage E16.5 in mouse development and lifted over to the human genome build^76^ (**Supplementary Methods**). Importantly, we found strong DLX1 and DLX5 peaks at non-coding WT-specific RNAPII signal anchoring WT-specific promoter-enhancer loops (**Fig. 5g**). We also found DLX1 and DLX5 peaks at non-coding APP(Swe)-specific RNAPII signal in APP(Swe)-specific promoter-enhancer loops (**Fig. S12e**). As predicted by our motif search, we observed that DLX1/DLX5 peaks were directly localized over TAATTA motifs at putative enhancers anchoring the base of both WT-specific and APP(Swe)-specific loops (**Fig. 5h-i, Fig. S12f**). Because *DLX1/5/6* gene expression only occurred in APP(Swe)-mutant neurons, and these proteins can bind to both WT-specific and APP(Swe)-specific looped enhancers, this provided the rationale for our initial working model that DLX TFs can operate as context-specific anti-loopers at WT-specific loops and looping facilitators at APP(Swe)-specific loops.

**Figure 5.**
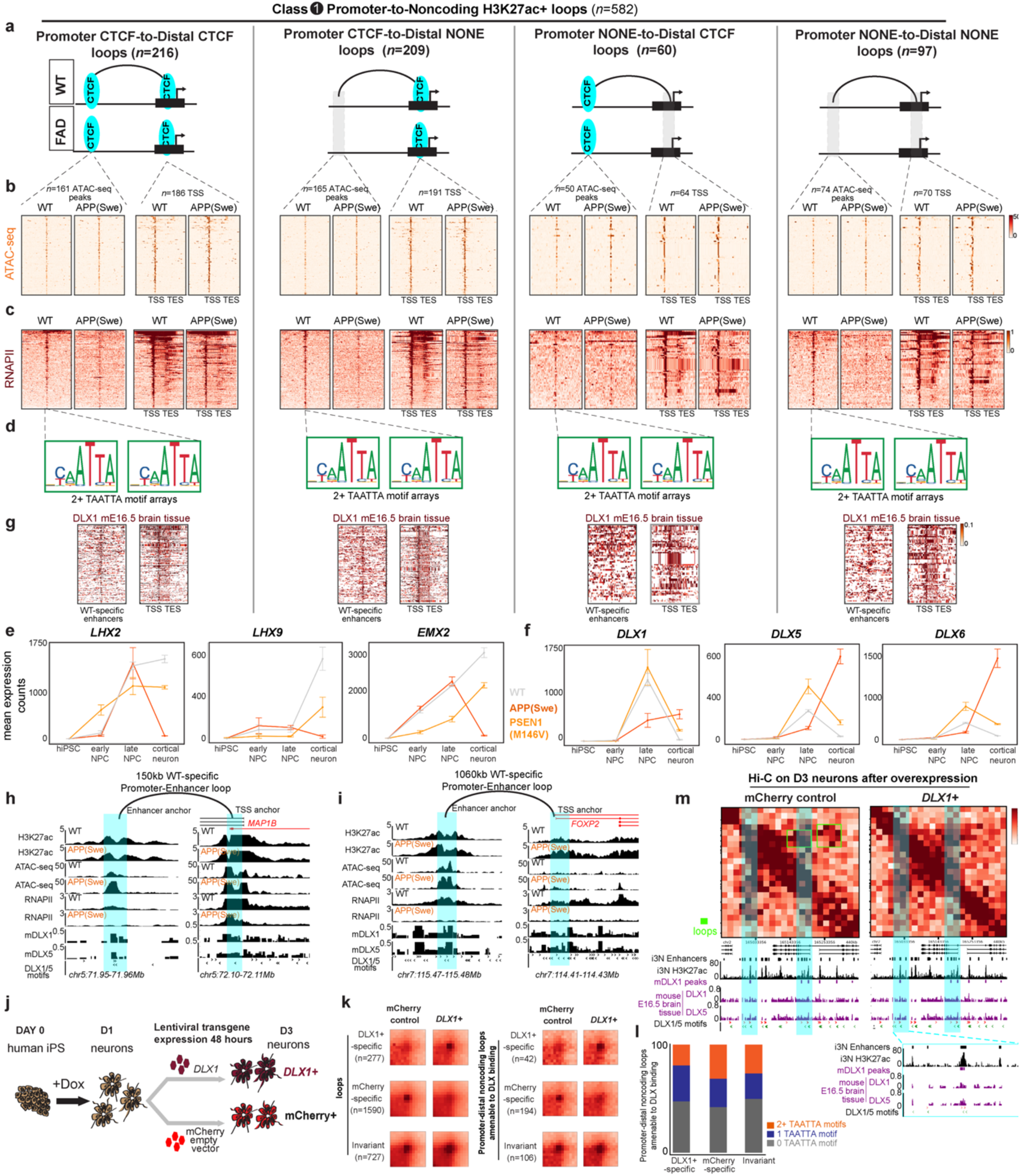
Reduced RNA polymerase II signal at tandem arrays of TAATTA motifs amenable to DLX binding at promoter-enhancer loops lost in FAD-mutant human neurons. (**a**) Stratification of WT-specific (class 1) Promoter-NoncodingH3K27ac-positive(+) loops into those anchored by (i) Promoter CTCF-to-Distal CTCF, (ii) Promoter CTCF-to-Distal NONE, (iii) Promoter NONE-to-Distal CTCF, and (iv) Promoter NONE-to-Distal NONE. **(b)** Heatmaps of ATAC-seq signal at WT-specific loops in WT and APP(Swe) human iPSC-derived neurons. **(c)** Heatmaps of RNAPII ChIP-seq signal at WT-specific loops in WT and APP(Swe) human iPSC-derived neurons. **(d)** Logograms of TAATTA motifs identified as strongly enriched at enhancers in WT-specific promoter-enhancer loops after MEME motif analysis (**Supplementary Methods**). (**e-f**) Mean normalized expression levels in bulk RNA-seq data for iPSC, early NPC, late NPC, and cortical neurons in WT, PSEN1(M146V) and APP(Swe) genotypes. Error bars represent standard deviation with n = 3 biological replicate experiments. **(e)** *LHX2* (NM_004789.4), *LHX9* (NM_001014434.2), and *EMX2* (NM_004098.4) isoforms. **(f)** *DLX1* (NM_001038493.2), *DLX5* (NM_005221.6), and *DLX6* (NM_005222.4) isoforms. **(g)** Heatmaps of DLX1 ChIP-seq signal from mouse E16.5 brain tissue at distal non-coding H3K27ac+ anchors in WT-specific promoter-enhancer loops (**Supplementary Methods**). **(h-i)** Locus-specific examples of enhancer and promoter loop anchors for WT-specific promoter-enhancer loops anchored by enhancers with WT-specific RNAPII, 2+ TAATTA motifs, and DLX1/5 chromatin occupancy signal, **(j)** Schematic of doxycycline-induced neural differentiation of i3N iPSCs overexpressing *DLX1* and *mCherry* lentiviral transgenes. **(k)** Aggregate peak analysis of Hi-C interaction frequency at mCherry-specific, *DLX1*-specific, and invariant loops between mCherry overexpression and *DLX1*+ overexpression conditions in human neurons. **(l)** Stacked barplot of the proportion of mCherry-specific, *DLX1*-specific, and invariant loops anchored by non-coding DLX peaks with 2+ TAATTA motifs, 1 TAATTA motif, or 0 TAATTA motifs. **(m)** Locus-specific example of a prototypical promoter-enhancer loop clearly abolished upon *DLX1* overexpression (mCherry-specific loop) in human neurons and anchored by 2+ TAATTA motifs and DLX1/5 chromatin occupancy signal at the non-coding enhancer.

We next set out to ascertain if there was a genomic feature which could distinguish DLX binding sites in WT-specific versus APP(Swe)-specific looped enhancers. Strikingly, we observed that DLX peaks at looped WT-specific enhancers exhibited a tandem array of 2 or more TAATTA motifs (**Fig. 5h-i, Fig. S12g).** By contrast, looped APP(Swe)-specific enhancers typically contained either 1 DLX-bound TAATTA motif or no TAATA motif or no DLX peak at all (**Fig. S12f-g).** These important observations led us to hypothesize that when DLX proteins bind to enhancers with multiple (2+) TAATTA motifs arranged in a tandem array they can attenuate loop interaction frequency (so-called anti-looping) and when they bind to enhancers with 0 or 1 TAATTA motifs they can form loops (so-called loop facilitation).

#### Overexpression of DLX1 in wild type neurons results in a similar pattern of miswired chromatin loops as in FAD-mutant neurons

Testing DLX’s role in looping is challenging via classic knock-down experiments because multiple homeodomain TFs are redundant and compete for the TAATTA motif^75, 76^. Therefore, to test our hypothesis and mimic *DLX1/5/6*’s strong upregulation in FAD-mutant neurons, we prioritized the overexpression of the DLX1 TF in a rapid neural differentiation model system (**Fig. 5j)**. We overexpressed a lentiviral mCherry transgene as a negative control. We cultured human i3N iPSCs expressing the Ngn2 transgene and initiated neural induction upon doxycycline addition (**Supplementary Methods**)^77, 78^. We transduced i3Ns at DIV1 with lentiviruses encoding the DLX1 and mCherry transgenes and used microscopy to confirm that i3Ns receiving the lentiviral constructs were mCherry-positive indicating high levels of transgene expression after 48 hours (**Fig. S13g)**. We confirmed that cells homogeneously exhibited morphology indicative of post-mitotic neurons (**Fig. S13g)**. After 48 hours of overexpression, we fixed and sorted neurons using fluorescence-activated cell sorting (FACs), quantified transduction efficiency (61% *mCherry*+, 24% *DLX1*+), and normalized for the number of transduced cells per condition (**Supplementary Methods**) (**Fig. S13h**).

We created Hi-C libraries at 20kb resolution in both conditions (*mCherry* control overexpression and *DLX1* overexpression (*DLX1+*)) (**Fig. 5j-m**). We identified 1,590 mCherry-specific loops, 277 *DLX1+-*specific loops, and 727 invariant loops between *mCherry+* and *DLX1+* conditions, thus suggesting more than a five-fold skew toward lost loops upon *DLX1* overexpression (**Fig. 5k, Supplementary Methods**). We stratified noncoding DLX1/5 peaks into those that bind to noncoding regions of the genome and have potential to bind DLX with either 0, 1, or 2+ TAATTA motifs. In cases of mCherry-specific loops lost upon *DLX1* overexpression, we observed a skew toward putative non-coding enhancers with 2+ TAATTA motif arrays (**Fig. 5l, Fig. S12g)**. By contrast, in cases of *DLX1+*-specific loops gained upon *DLX1* overexpression, we observed a skew toward putative non-coding enhancers with 0 or 1 TAATTA motifs (**Fig. 5l, Fig. S12g)**. Locus-specific examples confirmed the genome-wide predictions of multiple TAATTA motifs at mCherry-specific loops and a single TAATTA motif at DLX1+-specific loops after *DLX1+* overexpression in human neurons (**Fig. 5m, Fig. S12h**). Together, our results provide preliminary functional data suggesting that DLX homeodomain TFs can influence loop interaction frequency even in cases where architectural proteins CTCF and cohesin are bound and unchanging. Our data are consistent with our hypothesis that increased *DLX1* mRNA levels, mimicking what is seen in FAD-mutant neurons in vitro, can have two distinct effects on the contacts between promoters and enhancers: (i) attenuation of promoter-enhancer loops (as anti-loopers) anchored by 2+ TAATTA motifs and (ii) formation of promoter-enhancer loops (as loop facilitators) when anchored by 0-1 TAATTA motifs.

#### Single-cell RNA-seq reveals a shift from excitatory to inhibitory gene expression in FAD-mutant neurons coincident with DLX mRNA upregulation and miswiring of promoter-enhancer loops

The physiological relevance of promoter-enhancer loops remains a critical unanswered question. Using our FAD-mutant neurons as the model system, we finally set out to assess if increased *DLX1/5/6* mRNA levels and miswired loops coincided with changes to morphology or phenotype. Using phase contrast microscopy as well as immunostaining for pan-neural markers, we found no discernable change in morphology, density, viability, or the presence of non-neural populations in WT versus APP(Swe) neurons (**Fig. 6a)**.

We next used scRNA-seq in a matched number of n=5,539 cells for each genotype to assess the extent to which changes in cellular state might be occurring at the level of gene expression signatures (**Supplementary Methods**). Consistent with previous reports, we found that the large majority (87%) of cells in the WT genotype exhibited gene expression profiles characteristic of excitatory neurons (EX, cluster 0 and cluster 3), including cluster-defining expression of the vesicular glutamate transporter *SLC17A6*/*VGLUT2* and the glutamate receptor *GRID2* (**Fig. 6b-c, Fig. S14a**). These results confirm our ability to replicate previous reports and further demonstrate the purity of our differentiation protocol^36, 79^. In neurons with the APP(Swe) mutation, we observed a marked shift from excitatory to inhibitory expression profiles, including 50% neurons with cluster-defining expression of *GABRB2/GABRG2/TAC1* (cluster 1), 23% neurons with cluster-defining expression of *SSTR2/PAX6* (cluster 2), 17% neurons with cluster-defining expression of *GAD2/SSTR2* (cluster 4), and 7% neurons with cluster-defining expression of *GABRA2* (cluster 6) (**Fig. 6b-c, Fig. S14a**). Inhibitory clusters 1, 2, and 4, representing 90% of neurons, did not express *SLC17A6* and also expressed high levels of *DLX5* or *DLX6* (**Fig. S14a-b**). Our findings reveal a shift from excitatory to inhibitory gene expression in iPSC-derived neurons with homozygous FAD mutations coincident with upregulation of *DLX* mRNA levels.

We investigated whether single cells arising from the APP(Swe) genotype exhibited markers suggestive of a pure inhibitory subtype or if they showed evidence of a heterogeneous gene expression patterns suggestive of de-differentiation or reprogramming correlated with upregulation of *DLX1/5/6* mRNA. We identified two subpopulations that express both inhibitory and excitatory markers (cluster 4, cluster 6) (**Fig. 6b-c, Fig. S14b**). Both the *GAD2/SSTR2*-positive inhibitory neurons (cluster 4) and the *GABRA2*-positive inhibitory neurons (cluster 6) also express excitatory marker *GRID2* in the same single cells, suggesting presence of an intermediate/mixed Exc+Inh cellular state (**Fig. 6b, Fig. S14b**). Along with expression of the *GRID2* excitatory neuron marker, clusters 4 and 6 also expressed fewer inhibitory markers than clusters 1 and 2. Furthermore, single neurons in cluster 4 co-expressing *GRID2* and *GAD2/SSTR2* were also likely to express DLX5/6 **Fig. S14c**). Our observations suggest the possibility that at least some FAD-mutant neurons overexpressing *DLX1/5/6* are in a hybrid state expressing both excitatory and inhibitory markers.

**Figure 6.**
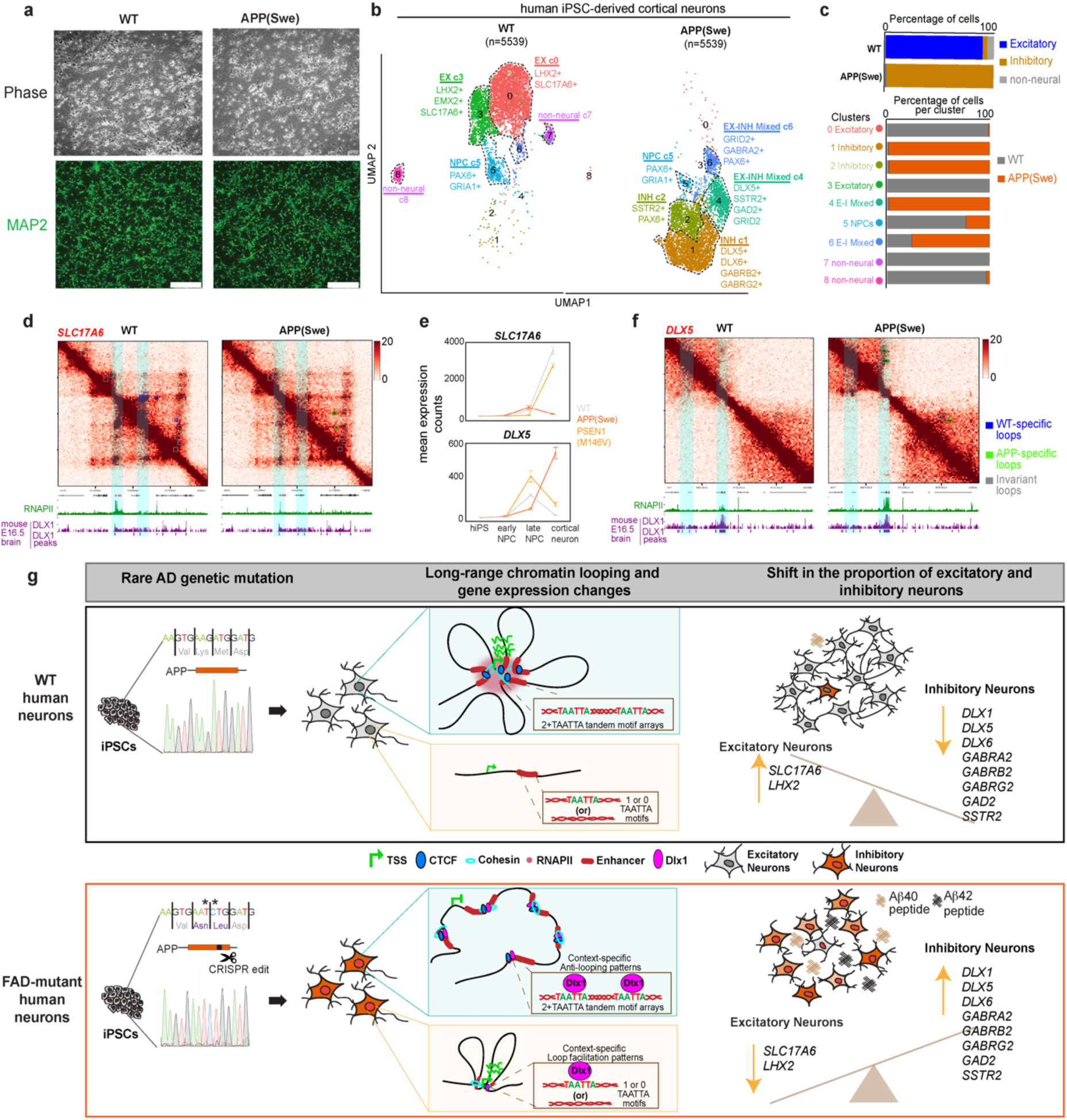
Single-cell RNA-seq reveals an excitatory to inhibitory shift in FAD-mutant neurons coincident with upregulation of *DLX1/5/6* mRNA and miswiring of promoter-enhancer multi-loops. **(a)** Representative phase contrast and immunofluorescence images from WT and APP(Swe) iPSC-neurons (DIV65) stained for mature cortical neuron marker MAP2 (green). Scale bars; 100μm. **(b-c)** Single-cell RNA-seq based clustering of n=5,549 WT and APP(Swe) iPSC-neurons. **(b)** Clusters are distinguished by color and labelled based on relevant cluster markers. **(c)** (top) Proportion of excitatory, inhibitory, and non-neural cells each cell type or (bottom) proportion of cells in all 8 clusters. **(d-f)** Hi-C heatmaps and bulk RNA-seq data in WT and APP(Swe) iPSC-derived neurons demonstrating multi-looped promoter-enhancer contacts anchored by DLX at **(d-e)** downregulated excitatory neural marker gene *SLC17A6* and **(e-f)** upregulated inhibitory neural marker *DLX5*. **(e)** For RNA-seq, mean normalized expression levels are shown for iPSC, early NPC, late NPC, and cortical neurons in WT, PSEN1(M146V), and APP(Swe) genotypes. **(g)** Schematic of multi-loop miswiring and transcriptional dysregulation linked to excitatory-inhibitory imbalance in FAD-mutant human neurons.

We finally set out to understand if WT-specific or APP(Swe)-specific loops anchored by enhancers with dynamic RNAPII and TAATTA motifs might be linked to changes in expression of excitatory or inhibitory neural subtype markers. We found that the excitatory neuron marker genes *SLC17A6* and *LHX2* are connected in WT-specific promoter-enhancer multi-loops anchored by DLX1/5 peaks and multi-TAATTA motif arrays (**Fig. 6d, Fig. S12i**). Such loops were disrupted in parallel with loss of RNAPII at the enhancers and downregulation of *SLC17A6* and *LHX2* in FAD-mutant neurons (**Fig. 6e top, Fig. S12j**). We also observed that the inhibitory neuron marker genes *DLX5/6* connected in APP(Swe)-specific promoter-enhancer multi-loops also anchored by DLX1/5 and 0-1 TAATTA motif as predicted by our computational analyses (**Fig. 6f**). Such loops were gained in parallel with gain of RNAPII signal at enhancers and upregulation of *DLX5/6* in FAD-mutant neurons (**Fig. 6e bottom**). Together, these locus-specific examples provide further evidence toward our working model in which *DLX1/5/6* overexpression occurs coincident with multi-loop miswiring, downregulation of multi-looped excitatory markers, and upregulation of multi-looped inhibitory markers in FAD-mutant neurons. Although we cannot use our model system to elucidate the order of molecular and cellular events, it is reasonable to conclude that loop miswiring is a critical step in a molecular cascade in which a rare, highly penetrant genetic mutation causes loop miswiring leading to pronounced shifts in excitatory to inhibitory gene expression programs.

## Discussion

How chromatin loops change and orchestrate gene expression in physiologically relevant cellular contexts and disease states in the mammalian brain represents an unanswered question with broad impact in genome biology^80^. Here, we set out to understand the genome’s structure-function relationship in post-mitotic human neurons using rare, penetrant, high effect size genetic mutation found in nature with physiologically relevant effects on neurodegeneration. We found genome-wide miswiring of loops in FAD-mutant neurons (**Fig. 6g**). When genes connect in FAD dynamic multi-loops, they exhibit strong changes in expression, including the upregulation of inhibitory neural marker genes and the downregulation of excitatory neural marker genes. Given their potential physiologic relevance and functional role, we explored the mechanisms underlying FAD-gained and -lost loops. We unexpectedly found that promoter-enhancer loops miswire despite negligible changes in chromatin occupancy patterns of the classic architectural proteins CTCF and cohesin. Instead, we observed that genes encoding the homeodomain TFs *DLX1/5/6* are strongly upregulated in FAD-mutant neurons. We overexpressed *DLX1* in wildtype neurons and demonstrated that it was sufficient to cause a similar pattern of loop miswiring. Using published ChIP-seq data, we found that TAATTA motif at enhancers anchoring both FAD-gained and -lost promoter-enhancer loops are amenable to DLX1/5 TF binding. At WT-specific promoter-enhancer loops lost in FAD-mutant neurons, we found lost occupancy of RNAPII and a tandem array of 2+ TAATTA motifs capable of binding DLX1/5 at enhancers (**Fig. 6g, top**). Such patterns are consistent with a role for DLX1/5 in context-specific anti-looping even when CTCF/cohesin occupancy remains unchanged (**Fig. 6g, top**). Moreover, at APP(Swe)-specific promoter-enhancer loops gained in FAD-mutant neurons, we found gained occupancy of RNAPII and 0-1 TAATTA motifs capable of binding DLX1/5 at enhancers (**Fig. 6g, bottom**). Such patterns are consistent with a role for DLX1/5 in context-specific loop facilitation even in cases where CTCF/cohesin binding is already present (**Fig. 6g, bottom**). Together, our work uncovers a link between multi-loop miswiring and dysregulated excitatory and inhibitory transcriptional programs during lineage commitment of human FAD-mutant neurons prior to formation of Aβ plaques and hyperphosphorylated tau tangles.

One plausible interpretation of our data is that gene expression and looping changes between wildtype and FAD-mutant neurons are only a consequence of the shift from excitatory to inhibitory neuron profiles. Here, we observe that the excitatory-inhibitory gene expression shift occurs in parallel with marked upregulation of *DLX1/5/6*. We also observe that DLX TFs can exhibit strong DNA binding signal at both WT-specific and APP(Swe)-specific looped enhancers, including at the base of FAD-disrupted *SLC17A6* and *DLX5/6* promoter-enhancer loops. Recently, Rubenstein and colleagues also provided data consistent with DLX TFs binding at the base of looped enhancers during early mouse brain development^75^. They emphasize that DLX TFs work in a context-specific manner to govern both down and upregulation of gene expression via distal enhancers during early mammalian brain development^75^. Similarly, here, we find data consistent with a possible role for DLX TFs as either anti-looper or loop facilitator depending on genomic context. Although we cannot use our model system to elucidate the order of molecular and cellular events, it is reasonable to conclude that *DLX* upregulation and DLX-mediated loop miswiring of key inhibitory and excitatory genes are critical steps in a molecular cascade in which a rare, highly penetrant genetic mutation causes gene dysregulation linked to a shift in excitatory-to-inhibitory gene expression programs. Future work should prioritize answering new lines of inquiry raised by this work; in particular, single-variable experiments dissecting how the rare genetic FAD mutations themselves versus changes in Aβ40/42 levels can kick-start miswiring of the genome’s structure-function relationship.

Excitatory-inhibitory (E-I) imbalance and defects in synaptic transmission are well-established phenomena observed during preclinical stages of AD^81–83^. For example, Aβ-induced increases in inhibitory GABAergic neurotransmission have been reported in transgenic mice heterozygously expressing a human APP gene with FAD mutations^84, 85^. Although such studies are focused on neural function, it is plausible that in models of FAD there could be changes in the proportion of excitatory, inhibitory, or hybrid E-I single neurons. However, depletion of cellular subtypes is underexplored and difficult to quantify in FAD due to the paucity of in vivo tissue across neurodevelopmental timepoints. Here, using isogenically-matched neurons with homozygous APP(Swe) mutations *in vitro*, we observe a shift from excitatory-to-inhibitory gene expression patterns coincident with *DLX1/5/6* overexpression and miswired DLX-bound multi-loops. The APP(Swe) mutation is heterozygous in FAD patients; therefore, the effect size of *in vivo* changes in biological pathways are diluted by the presence of the mutation on only one allele and further confounded by a myriad of additional biological pathways and cell types influencing neuronal cell type-specific gene expression. To overcome these challenges and focus on cell autonomous neuron-specific changes in FAD, we chose to employ isogenic iPSC-derived neurons engineered with homozygous APP(Swe) mutations and a differentiation system achieving pure post-mitotic neuron populations. Our model allows for the quantification of signal above noise given the high effect size and clear direction of effect of a homozygously-engineered rare variant, thus allowing us to find clear molecular and cellular disruptions in early neural lineage commitment prior to accumulation of amyloid and phosphorylated tau. We also hypothesize that our ability to observe a shift from excitatory to inhibitory gene expression in vitro may be enhanced by the presence of homozygous mutations inducing unusually high levels of amyloid peptides and thus represent hyper-accelerated dysregulation of molecular pathways. Our model may amplify cellular phenomena that are slightly different or of lower effect size in human tissue or transgenic models, thus allowing us to clearly link a genetic rare variant important for human disease to chromatin folding disruptions, gene expression dysregulation, and a potentially physiologically relevant phenotype of shifting excitatory-inhibitory neuronal populations.

There is a gap in knowledge of how genetic mutations and chromatin marks mechanistically interplay to give rise to the severity and timing of cellular phenotypes in earliest stages of neurodegenerative disease^86^. Transcriptomic and epigenomic mapping studies have led to the identification of thousands of differentially expressed genes in neural and non-neural single cells from SAD post-mortem brain tissue^13, 28, 87, 88^. However, late-stage tissue profiling ultimately conflates signal from multiple cell types and obscures the mechanisms that drive the initial onset of pathologic phenotypes. Often the primary affected cell type will not be present due to degeneration. In the case of FAD, chromatin and gene expression studies in post-mortem brain tissue are rare^33^. Therefore, FAD insights are driven by transgenic mouse models and human iPSC models with rare mutations in the *APP*, *PSEN1*, or *PSEN2* genes. Consistent with our current data, classic early studies differentiating iPSCs with rare FAD mutations to neurons confirmed the expected known preclinical phenotypes of increased total Aβ, shifts in the ratio of Aβ42:40 peptides, increased levels of Aβ oligomers, and the absence of Aβ aggregates in monolayer culture^34–36, 89–91^. In the last five years, two additional studies reported thousands of dysregulated genes in iPSC-neurons with FAD mutations, with overlap of many of the same dysregulated genes among lines with different mutants^34, 35^. Although limited in number, the published results confirm the veracity of the gene expression phenomenon reported here. Only one FAD iPSC-neuron chromatin study is available to our knowledge, and it focuses on classic epigenetic modifications to the linear 10 nm chromatin fiber and mutations only in the *PSEN1* gene^35^. Thus, there is a critical need to understand how individual cell types undergo changes to chromatin and higher-order folding during the early onset of FAD, and our observations add unique insight to a major knowledge gap in the neurodegenerative disease field.

A decade or more of imaging, genomics, and computational technology development has enabled the mapping of loops genome-wide, but their functional role in controlling gene expression has remained an unresolved question. Seminal molecular tagging and Chromosome-Conformation-Capture (3C) studies provided the initial evidence that non-coding loci with enhancer activity can physically contact distal genes in parallel with their upregulation^57–59^. Contemporary studies disrupting loops genome-wide via short-term degradation of architectural proteins initially showed minor direct effects on mRNA levels and transcription^72, 92^. Based on these results, an initially controversial conclusion emerged that loops were unnecessary for gene expression maintenance in dividing cell lines. Here we focus on the use of non-dividing, post-mitotic neurons and a perturbation found in nature (rare FAD mutations) to assess the genome’s structure-function relationship. We find that the severity of fold change in expression in FAD neurons is directly correlated with the number of disrupted loops the promoter engages; FAD upregulated genes encoding inhibitory neural markers connect in multiple gained loops and FAD downregulated genes encoding excitatory neural markers lose multiple loops. Together, our data suggest that a genome structure-function relationship can be found if one uses physiologically relevant model systems of human neurons with mutation occurring in known neurological disorders.

Altogether, our results suggest that miswiring of multi-loops is a critical contributor to dysregulated excitatory-to-inhibitory gene expression programs in FAD-mutant human neurons. Although ascertaining the order of events is beyond the scope of the current study, our work uncovers new observations to inform a testable model of the genome’s structure-function relationship bridging scales of genetics, chromatin, gene expression, and neural cell phenotype.

## Acknowledgments

We thank members of the Cremins lab for helpful discussions. H. Ryu is the recipient of the Michael S. Brown Graduate Research Fellowship. A. Nikish is the recipient of an NIH T32 training grant. K. Titus is the recipient of a Fontaine fellowship. J. E. Phillips-Cremins was a New York Stem Cell Foundation Robertson Investigator and an Alfred P. Sloan Foundation Fellow during this project.

## Author contributions

Conceptualization: ZS, HChan, JEPC

Methodology/Visualization: HChan, ZS, HChoi, AW, AN, HR, SM, WG, DP

Investigation: HChan, ZS, JEPC

Funding: DP, JEPC Administration: JEPC

Writing & Editing: HChan, ZS, AW, AN, HR, HChoi, DP, JEPC

Reagents: DP

## Data and materials availability

Data and code are provided to reviewers via confidential links and tokens.

All data and code will be made freely available to the scientific community upon publication.

