## Supplemental Methods and Figures for "A multi-looping chromatin signature predicts dysregulated gene expression in neurons with familial Alzheimer’s disease mutations"

### Supplementary Tables

Supplementary Table 1: Soluble and oligomeric amyloid beta quantification from human iPSC-derived 2D neurons and 3D organoids.

Supplementary Table 2. Genome-wide normalized counts and differential mRNA levels across stages of human iPSC to neuron 2D differentiation

Supplementary Table 3. Normalized counts and differential mRNA levels for Common FAD Down, Common FAD UP, and genotype-invariant expressed mRNA transcripts in human iPSC-derived neurons

Supplementary Table 4. Genome-wide normalized counts and differential expression in NeuN+ sorted nuclei from human iPSC-derived D100 and D200 WT and APP(Swe) brain organoids

Supplementary Table 5. ATAC-seq peaks identified in WT, APP(Swe), and PSEN1(M146V) human iPSC-derived neurons

Supplementary Table 6. Differential ATAC-seq peaks between WT and APP(Swe) human iPSC-derived neurons

Supplementary Table 7. Sequencing depth and mappability for all datasets generated in this study

Supplementary Table 8. TADs/subTADs genome-wide from Hi-C in WT, APP(Swe), and PSEN1(M146V) human iPSC-derived neurons

Supplementary Table 9. Loops genome-wide from Hi-C in WT, APP(Swe), and PSEN1(M146V) human iPSC-derived neurons

Supplementary Table 10. WT-specific, APP(Swe)-specific, and Genotype-invariant loop calls genome-wide between WT and APP(Swe) human iPSC-derived neurons

Supplementary Table 11. WT-specific, APP(Swe)-specific, and Genotype-invariant loops stratified into Promoter-to-Promoter, Promoter-to-NonPromoter, and NonPromoter-to-NonPromoter loop subclasses and further substratified by CTCF and H3K27ac signal in human iPSC-derived neurons

Supplementary Table 12. Gene isoforms with promoters anchoring WT-specific loops, APP(Swe)-specific loops, genotype-invariant loops, and not looping in human iPSC-derived neurons

Supplementary Table 13. Burst size and burst frequency computed genome-wide from single-cell RNA-seq data produced in WT and APP(Swe) human iPSC-derived neurons

Supplementary Table 14. Genome-wide CTCF, H3K27ac, RAD21, and RNA Polymerase II peaks identified in WT, APP(Swe), and PSEN1(M146V) human iPSC-derived neurons

Supplementary Table 15. Genome-wide putative non-coding cis regulatory elements positive for H3K27ac signal called in WT and APP(Swe) human iPSC-derived neurons

#### Supplementary Movies

Movie S1. Representative movie from confocal z-stack immunofluorescence images from DIV200 organoids derived from WT iPSCs stained for neuron marker MAP2 (red), DAPI (blue), Aβ (green, 4G8 antibody). A 50x50μm image is shown as acquired with a 63x objective.

Movie S2. Representative movie from confocal z-stack immunofluorescence images from DIV200 organoids derived from APP(Swe) iPSCs stained for neuron marker MAP2 (red), DAPI (blue), Aβ (green, 4G8 antibody). A 50x50μm image is shown as acquired with a 63x objective.

#### Supplementary Figures

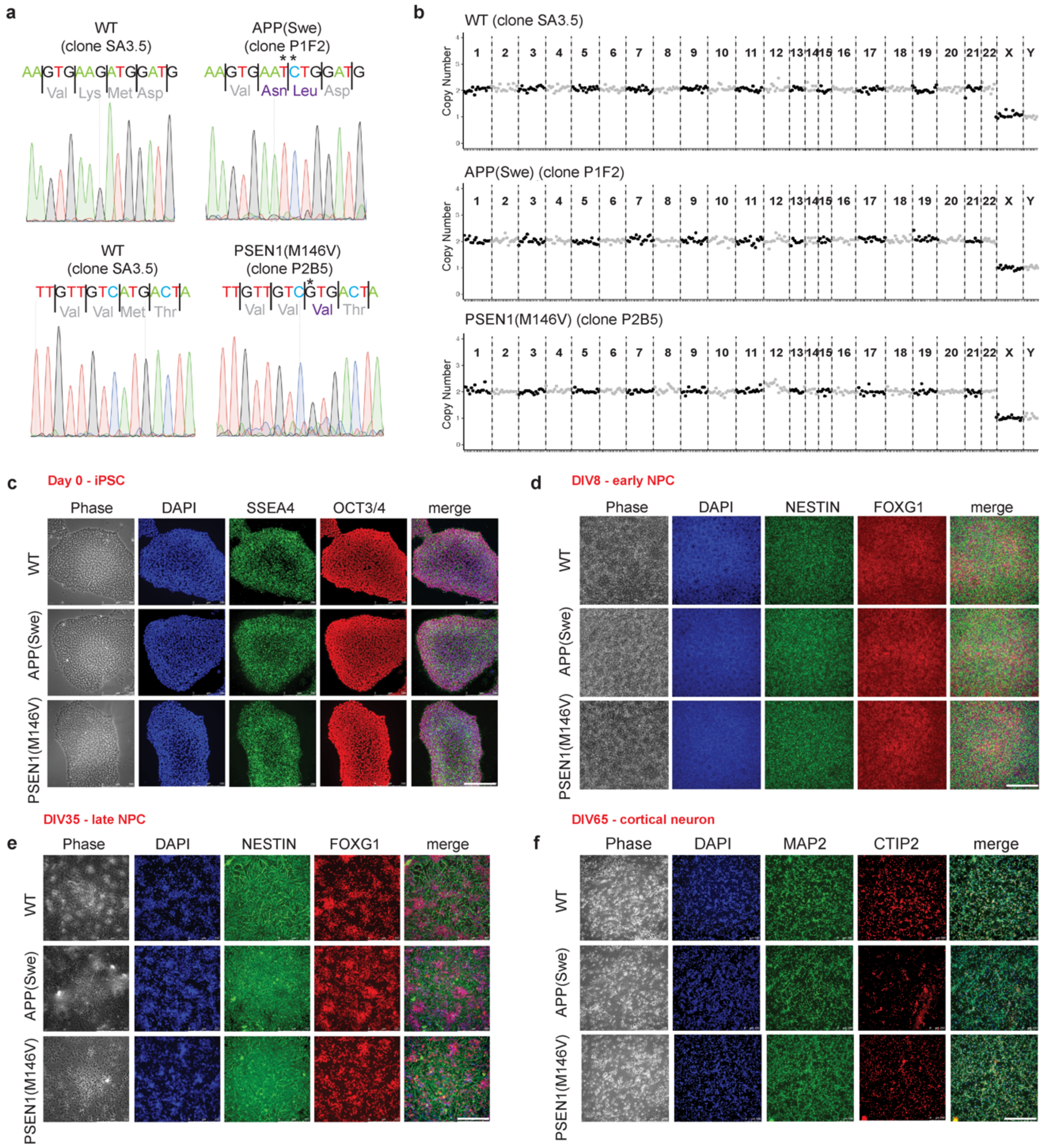


**Supplementary Figure 1: *In vitro* characterization of human iPSC-derived neural progenitors and cortical neurons with WT, APP(Swe), and PSEN1(M146V) genotypes.** **(a)** Sanger sequencing reads of APP(Swe) and PSEN1(M146V) CRISPR edited human iPSC lines, including: SA3.5 (WT), P1F2 (rare APP(Swe) mutation), and P2B5 (rare PSEN1(M146V) mutation). **(b)** iPSC lines were assessed as karyotypically normal using the nCounter human karyotype panel assay containing 338 probes across the 24 human chromosomes. **(c)** Representative immunofluorescence images from WT and FAD iPSCs (Day 0), **(d)** early neural progenitor cells (eNPC, DIV8), **(e)** late NPC (lNPC, DIV35) and **(f)** neurons (DIV65) stained for stem cell (SSEA4 (green), OCT3/4(red)) markers; anterior forebrain (FOXG1(red)) and pan-NPC (NESTIN (green)) markers; and post-mitotic cortical neuron (MAP2(green), CTIP2(red)) markers. Scale bars; 100µm.

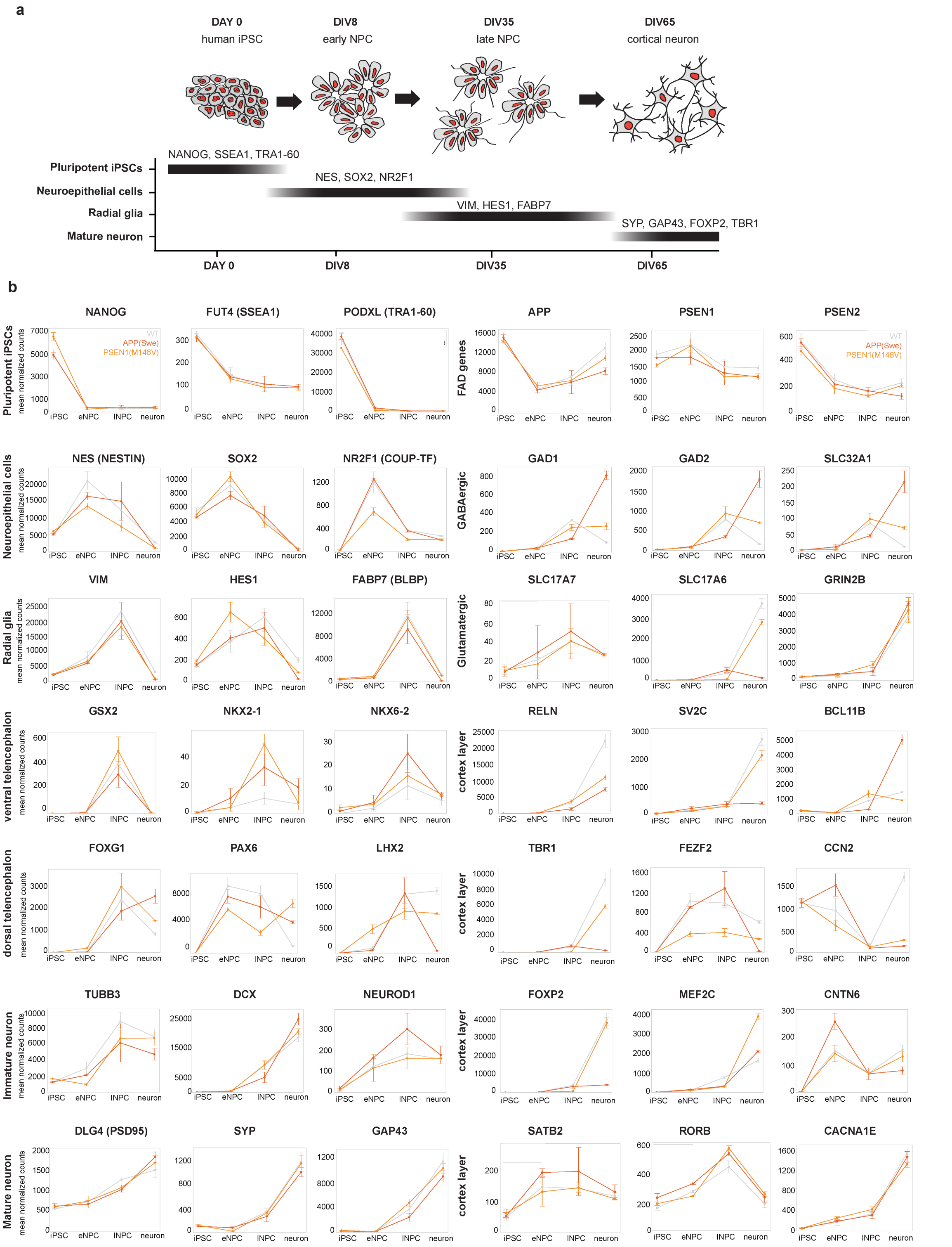


**Supplementary Figure 2. RNA-seq characterization of mRNA levels in human iPSC-derived neural progenitors and cortical neurons with WT, APP(Swe), and PSEN1(M146V) genotypes. (a)** Cartoon schematic representing expected gene expression patterns for known markers of the 4 major differentiation states. **(b)** Normalized mRNA levels (TPM) for iPSC, early NPC, late NPC, and cortical neuron cellular states in WT, PSEN1(M146V), and APP(Swe) genotypes. Error bars, standard deviation using n = 3 biological replicates.

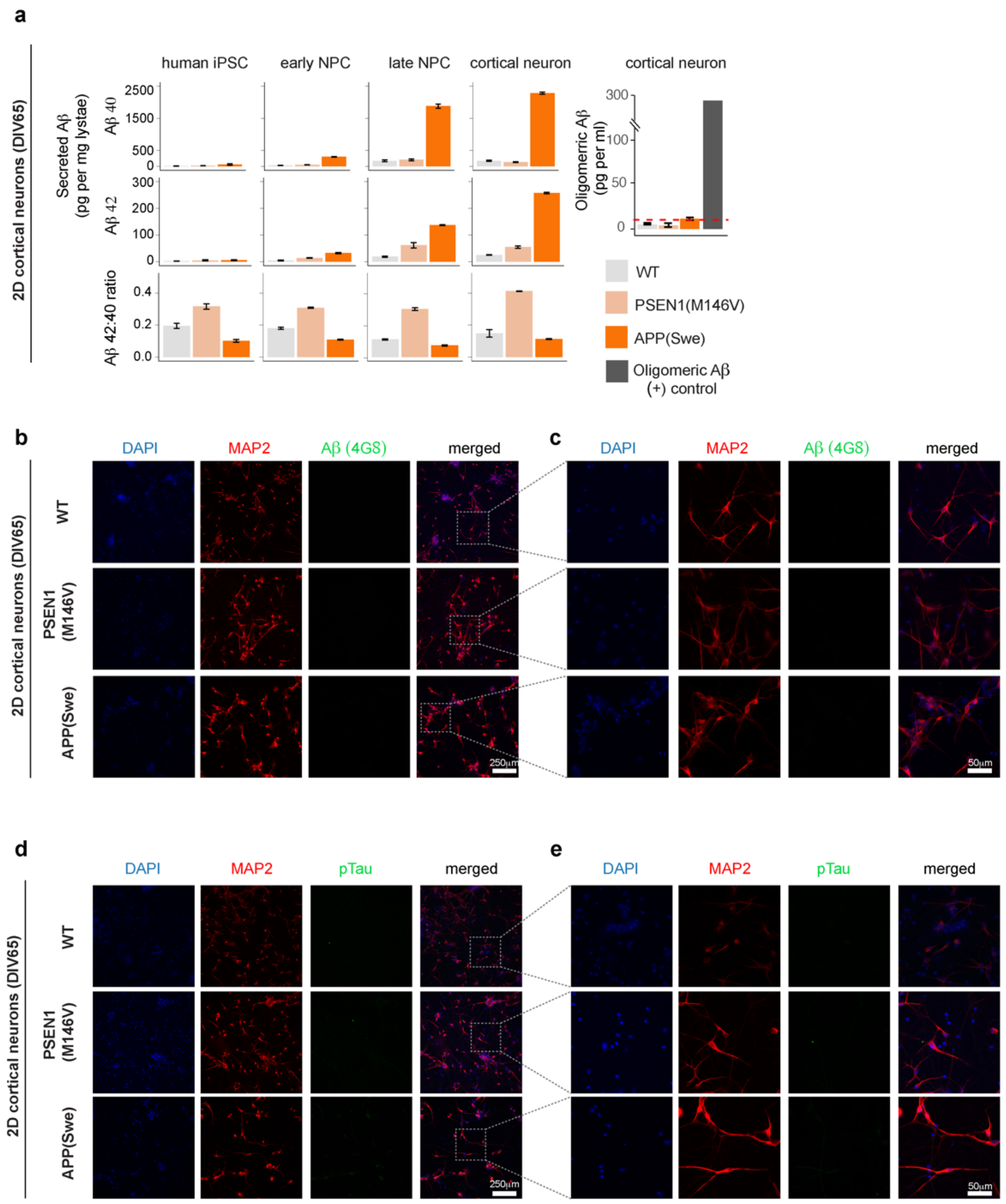


**Supplementary Figure 3: Characterization of amyloid-β (Aβ) secretion and Tau phosphorylation in 2D human iPSC-derived neurons.** (**a**) Aβ40, Aβ42, and Aβ42:40 ratios in WT, APP(Swe), and PSEN1(M146V) genotypes across iPSC, early NPC, late NPC, and neurons. Oligomeric Aβ measured across genotypes in neurons. Red line indicates the assay's detection limit. (**b, c**) Representative immunofluorescence images from WT, APP(Swe), and PSEN1(M146V) 2D neurons (DIV65) stained for nuclei (DAPI, blue), MAP2 (red), Aβ (4G8 antibody, green). Scale bars: (**b**) 250μm, (**c**) 50μm. (**d, e**) Representative immunofluorescence images from WT, APP(Swe), and PSEN1(M146V) 2D neurons (DIV65) stained for nuclei (DAPI, blue), MAP2 (red), pTau (AT8 antibody, green). Scale bars: (**d**) 250μm, (**e**) 50μm.

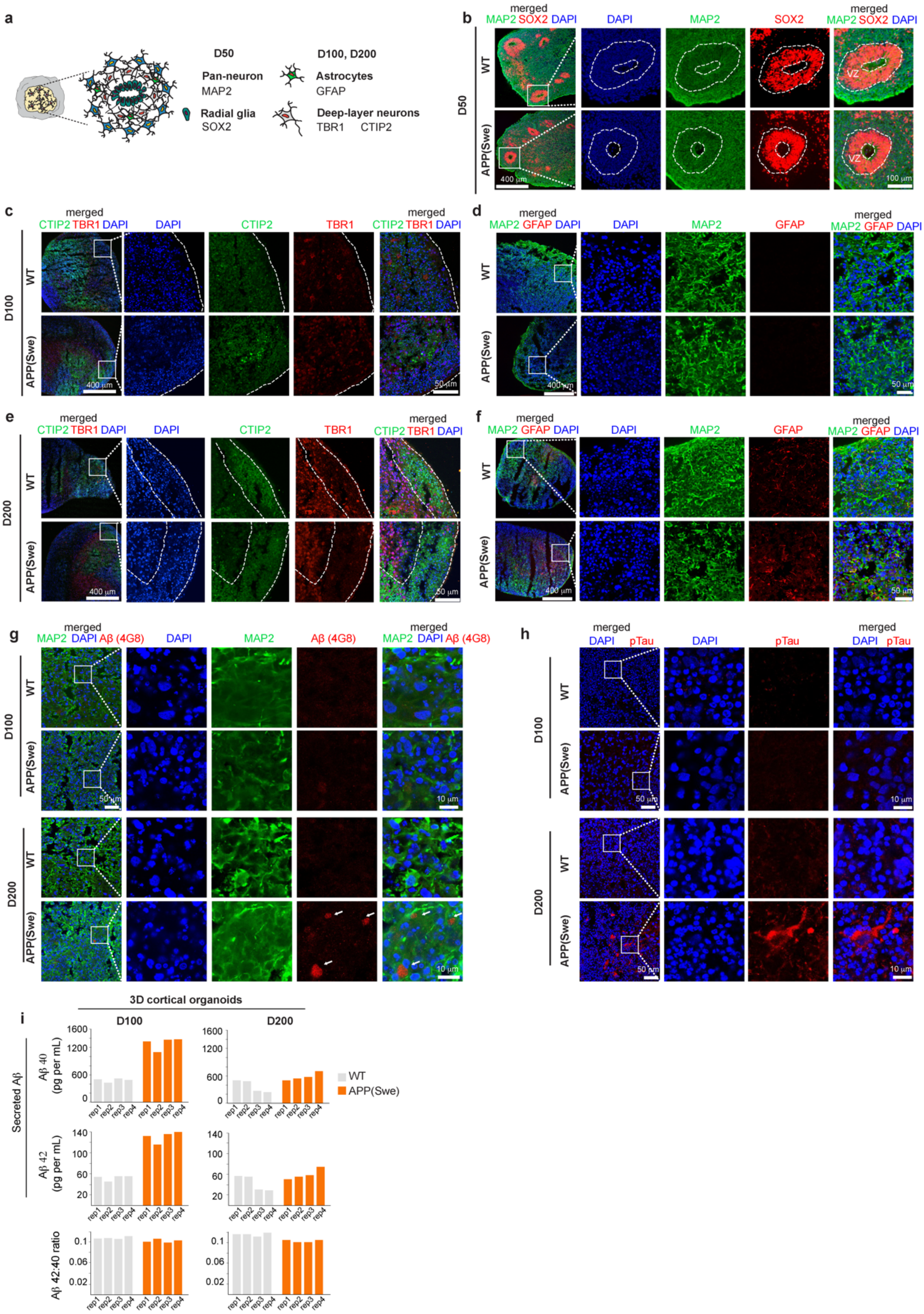

**Supplementary Figure 4: Characterization of cellular markers, Aβ- and phospho-tau accumulation in WT and APP(Swe) human iPSC-derived 3D brain organoids. (a)** Schematic representation of the organoid architecture and cellular markers at D50, D100, and D200 in culture. **(b)** Confocal images in WT and APP(Swe) organoids at D50 of differentiation showing the ventricular zone (VZ)-like structure formed by newborn neurons (MAP2, green) and neural progenitor cells (SOX2, red). Scale bars, 400µm and 100µm; (**c-**

**f)** Confocal images in WT and APP(Swe) organoids at (**c-d**) D100 and (**e-f**) D200 of differentiation showing (**c, e**) the expected cortical layer structure by D200 using CTIP (green) and TBR1(red) to label deep layer cortical neurons and (**d, f**) the expected emergence of astrocytes by D200 using GFAP (red). Scale bars, 400μm and 50μm. (**g-h**) Confocal images in WT and APP(Swe) organoids at D100 and D200 of differentiation stained for (**g**) mature neuron marker (MAP2, green), Aβ (4G8 antibody, red), and DAPI (blue) or (**h**) pTau (AT8 antibody, red) and DAPI (blue). Scale bars, 10μm. (**i**) Aβ40, Aβ42, and Aβ42:40 ratios in supernatants collected from D100 and D200 organoids produced from WT and APP(Swe) iPSCs. Each condition, n = 4 biological replicates.

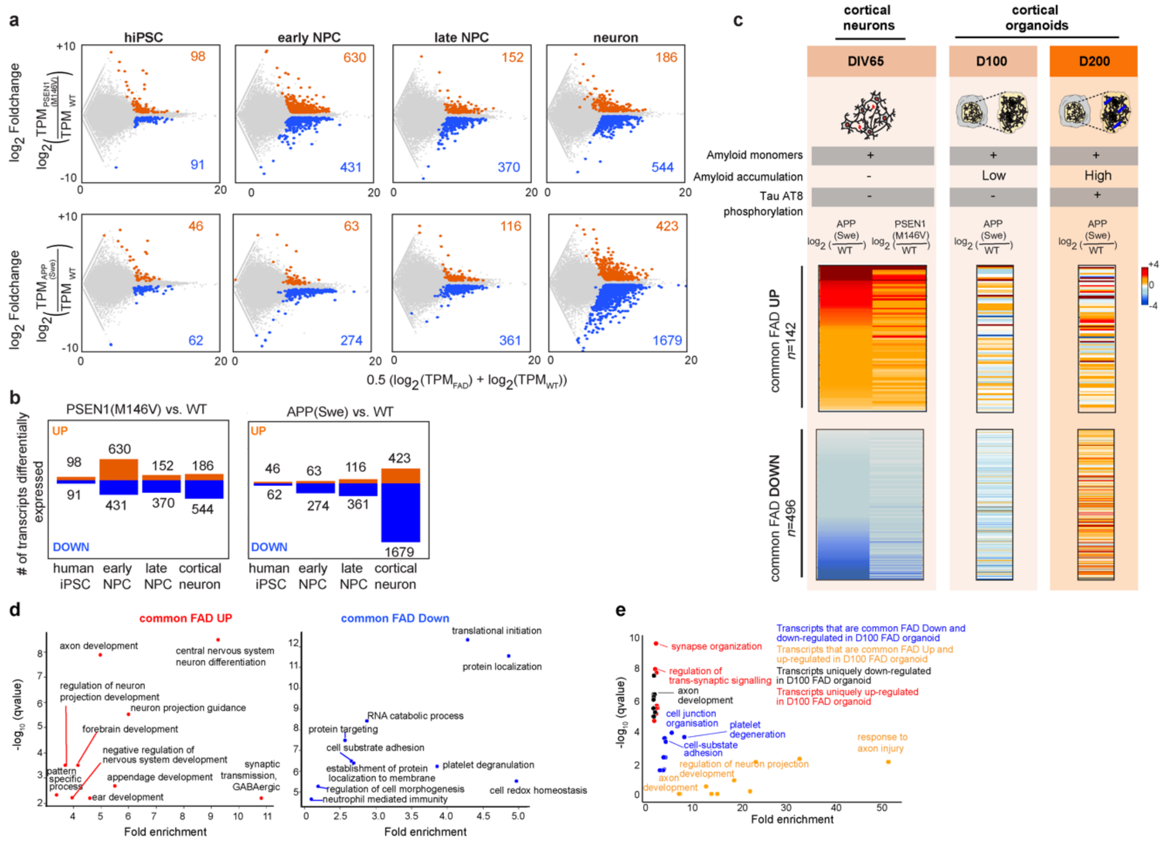

**Supplementary Figure 5: Comparison of differential gene expression in 2D monolayer versus 3D organoid cultured iPSC-derived neurons with rare familial AD mutations.** **(a)** MA plot displaying RNA-seq data in human iPSCs, early NPCs, late NPCs, and neurons comparing PSEN1(M146V) (top row) or APP(Swe) (bottom row) genotypes to isogenic WT control. Differentially expressed mRNA transcripts called at q-value<0.05 and |log2(fold-change)|>0.32. **(b)** Number of differentially expressed mRNA transcripts identified in PSEN1(M146V) and APP(Swe) across the neural differentiation stages. **(c,** ***top*****)** Schematic summary of observed patterns of amyloid monomers, amyloid accumulation, and tau phosphorylation in iPSC-derived neurons cultured in 2D monolayer and in D100/D200 cortical organoids. (**c,** ***bottom*****)** Heatmap comparison of fold change in gene expression of common FAD UP transcripts (*n*=142) and common FAD DOWN transcripts (*n*=496) in APP(Swe) or PSEN1(M146V) compared to isogenic WT iPSC-derived neurons cultured in (column 1) 2D monolayer or in (columns 2/3) APP(Swe) D100/D200 cortical organoids. (**d**) Gene Ontology analysis for transcripts for common FAD UP transcripts (*n*=142) and common FAD DOWN transcripts (*n*=496). (**e**) Gene Ontology analysis for transcripts dysregulated in 2D monolayer and D100 FAD organoids.

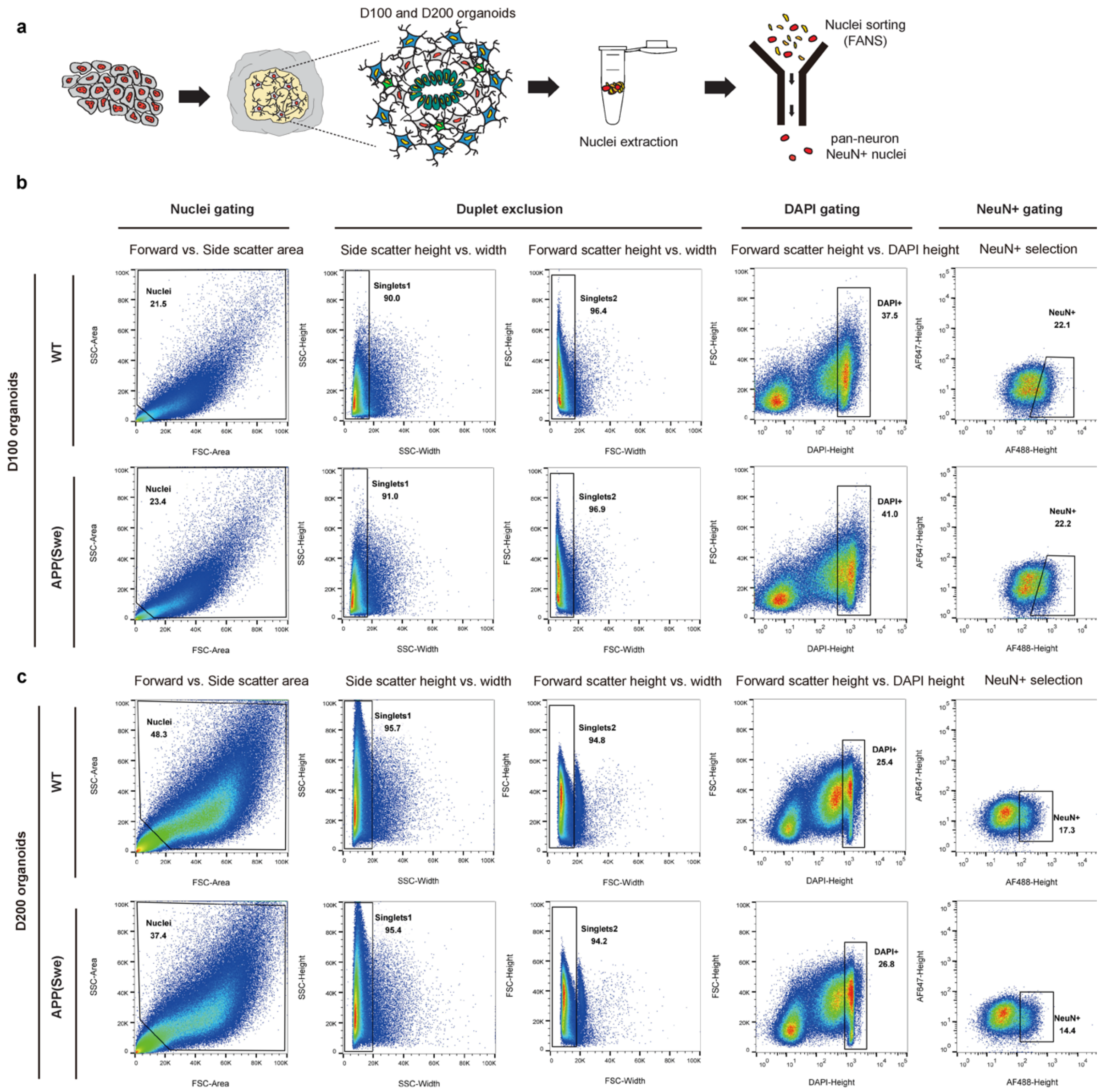


**Supplementary Figure 6: Isolation and purification of neuronal nuclei from human iPSC-derived organoids by FANS. (a)** Workflow summarizing differentiation of human iPSC-derived organoids and Fluorescent Activated Nuclei Sorting (FANS). **(b, c)** Overview of the NeuN+ nuclei sorting strategy for WT and APP(Swe) (**b**) D100 and (**c**) D200 organoids. forward scatter (FSC) versus Side scatter (SSC) dot plot showing gating on live, average-sized cells, followed by sequential gating on SSC width versus height and FSC width versus height for duplet exclusion. Single cells were then gated to collect DAPI+, NeuN+ nuclei.

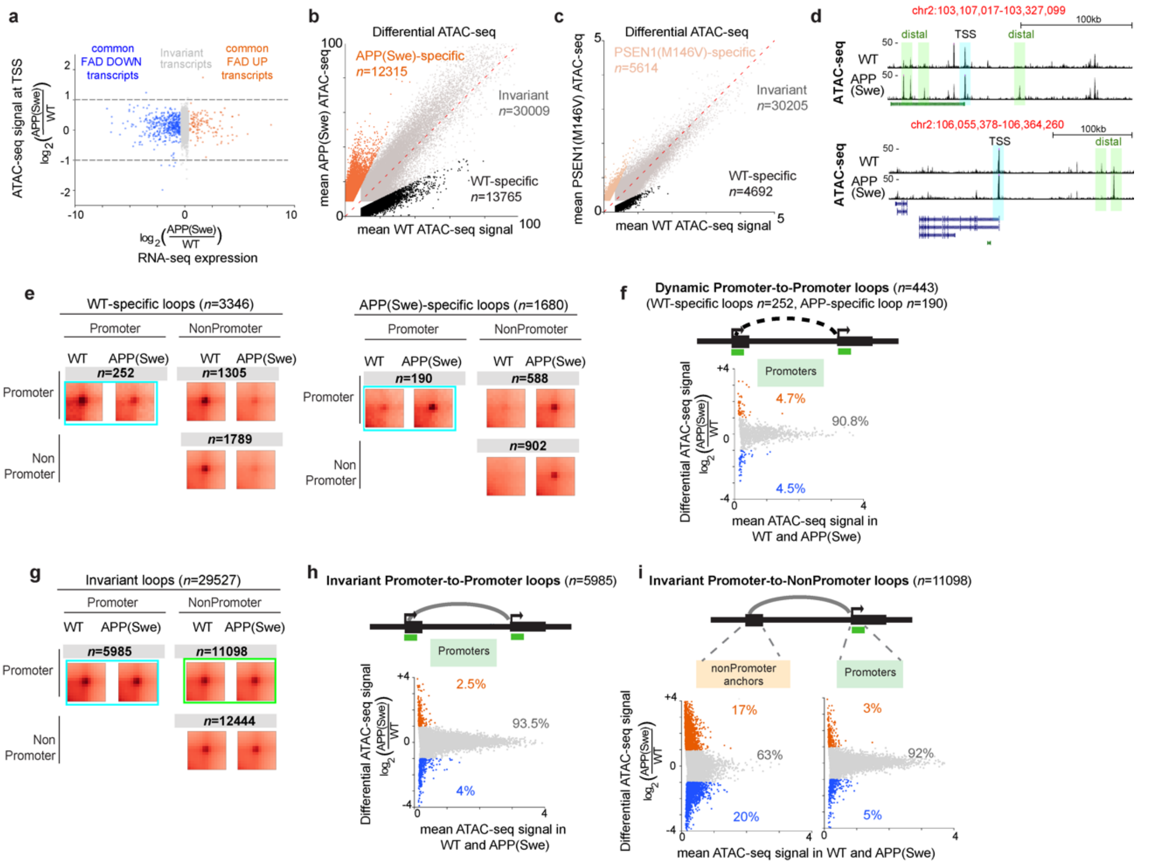


**Supplementary Figure 7: Chromatin accessibility at promoters is invariant and does not correlate with changes in gene expression or loop interaction frequency due to rare FAD mutations in neurons. (a)** Scatterplot of fold change in ATAC-seq signal and gene expression in WT and APP(Swe) iPSC-derived neurons for promoters of common FAD DOWN transcripts (*blue*), common FAD UP transcripts (*orange*), and invariantly expressed transcripts (*grey*). **(b-c)** Scatterplot of ATAC-seq signal at peaks of chromatin accessibility in WT human iPSC-derived neurons versus those with (**b**) APP(Swe) or (**c**) PSEN1(M146V) rare FAD mutations. WT-specific (black), Genotype-invariant (grey), and APP(Swe)- or PSEN1(M146V)-specific (orange) ATAC-seq peaks were computed as in the Supplementary Methods. **(d)** Genome browser example of ATAC-seq tracks in WT and APP(Swe) human iPSC-derived neurons highlighting changes in ATAC-seq signal at distal non-coding regions (green) but not at peaks in promoters (cyan). **(e)** Aggregate Peak Analysis (APA) across all (left) WT-specific loops and (right) APP(Swe)-specific loops in each of the Promoter-to-Promoter, Promoter-to-nonPromoter, and nonPromoter-to-nonPromoter loop classes comparing Hi-C interaction frequency in WT and APP(Swe) human iPSC-derived neurons. **(f)** MA plot of differential ATAC-seq signal between WT and APP(Swe) human iPSC-derived neurons in a ±2kb window around TSSs anchoring WT-specific or APP(Swe)-specific dynamic Promoter-to-Promoter loops. **(g)** Aggregate Peak Analysis (APA) across all genotype-invariant loops in each of the Promoter-to-Promoter, Promoter-to-nonPromoter, and nonPromoter-to-nonPromoter loop classes comparing Hi-C interaction frequency in WT and APP(Swe) human iPSC-derived neurons. **(h-i)** MA plot of differential ATAC-seq signal between WT and APP(Swe) human iPSC-derived neurons in a ±2kb window around TSSs anchoring (**h**) genotype-invariant Promoter-to-Promoter loops or (**i**) genotype-invariant Promoter-to-NonPromoter loops.

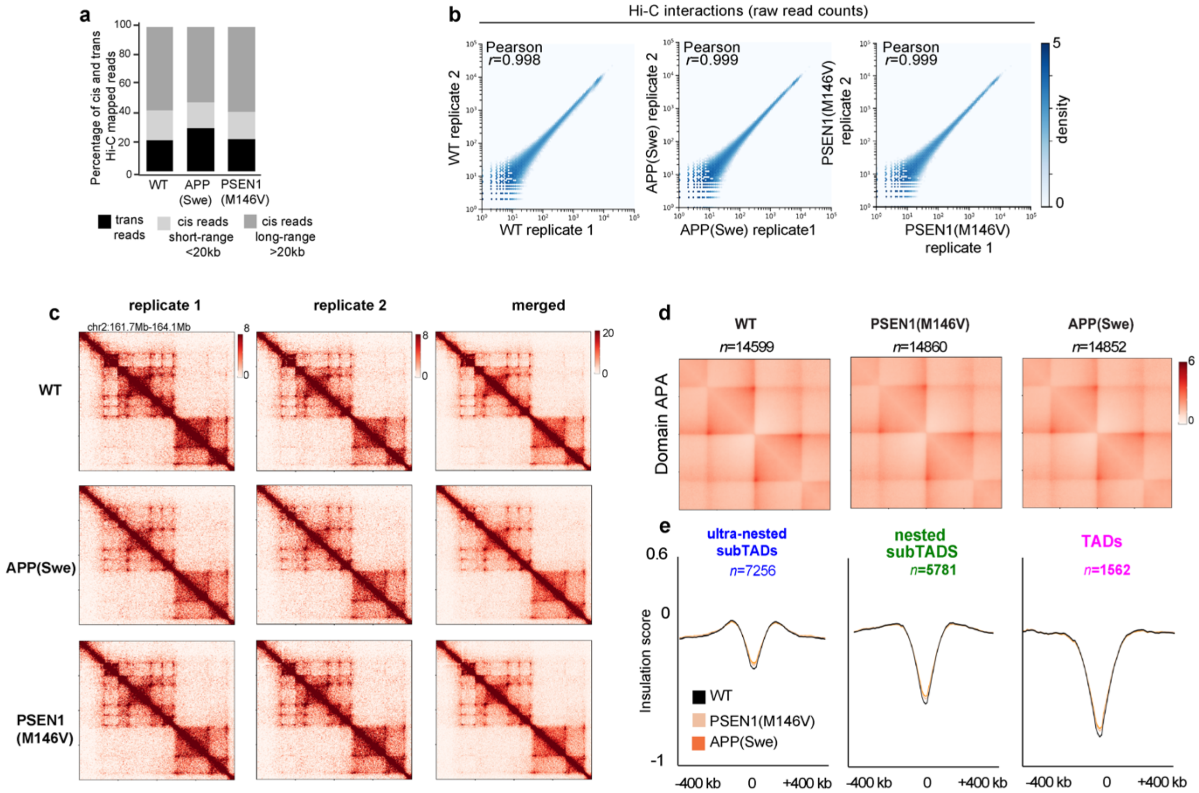


**Supplementary Figure 8**: **Hi-C data demonstrates that TADs and subTADs are unchanged in human iPSC-derived neurons with FAD mutations. (a)** Distribution of cis and trans Hi-C reads across WT, APP(Swe), and PSEN1(M146V) human iPSC-derived neurons. **(b)** Hi-C libraries for two biological replicates are highly correlated for all 3 genotypes in iPSC-derived neurons. Bin size, 250 kb. **(c)** Hi-C interaction frequency heatmap for a representative region (*chr2:161.7-164.1Mb*) across all 3 genotypes in individual biological replicates and in merged. Bin size, 10 kb. **(d)** Genome-wide TAD/subTAD APA plots for each of the three genotypes; (**e**) Mean insulation score computed in WT, APP(Swe), and PSEN1(M146V) iPSC-derived neurons computed for TAD and subTAD boundaries called in WT neurons.

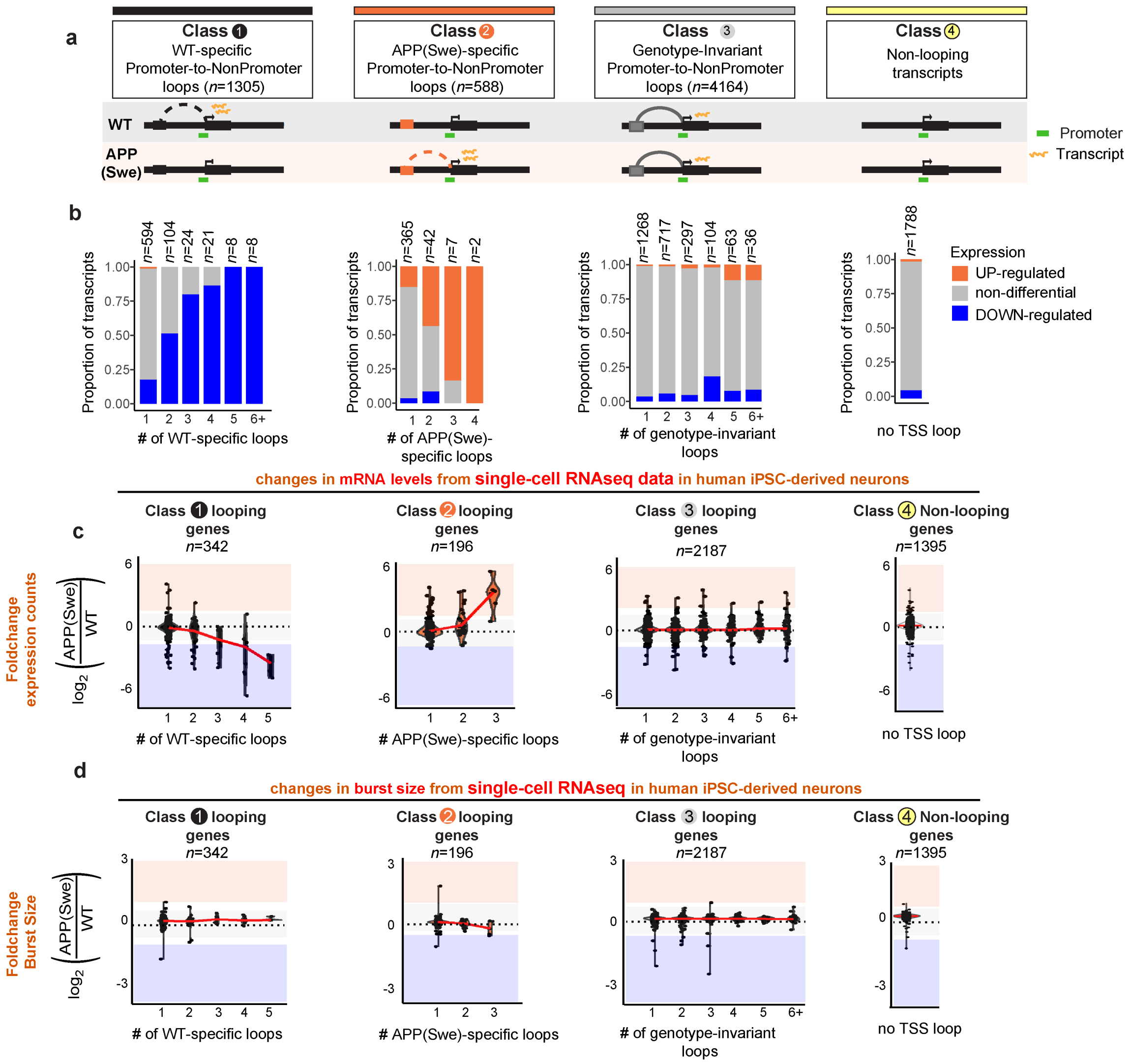


**Supplementary Figure 9: The number of broken Promoter-to-NonPromoter loops correlates with the severity and direction of the dysregulation of single-cell mRNA levels but not burst size. (a)** Cartoon representation of transcripts in WT-specific (class 1), APP(Swe)-specific (class 2), and genotype-invariant (class 3) Promoter-to-NonPromoter loops as well as non-looping promoters (class 4). **(b)** Stacked bar plot showing proportion of mRNA transcripts that are upregulated (orange), downregulated (blue), or unchanged (grey) in APP(Swe) versus WT human iPSC-derived neurons. (**c-d**) Strip plot of the fold change in two single-cell RNA-seq metrics stratified by the number of genotype-dynamic or -invariant Promoter-to-NonPromoter loops formed per transcript. (**c**) mean single-cell RNA-seq normalized count per gene (log2(APP(Swe)/WT)). (**d**) mean single-cell burst size per gene (log2(APP(Swe)/WT)). Red line, median.

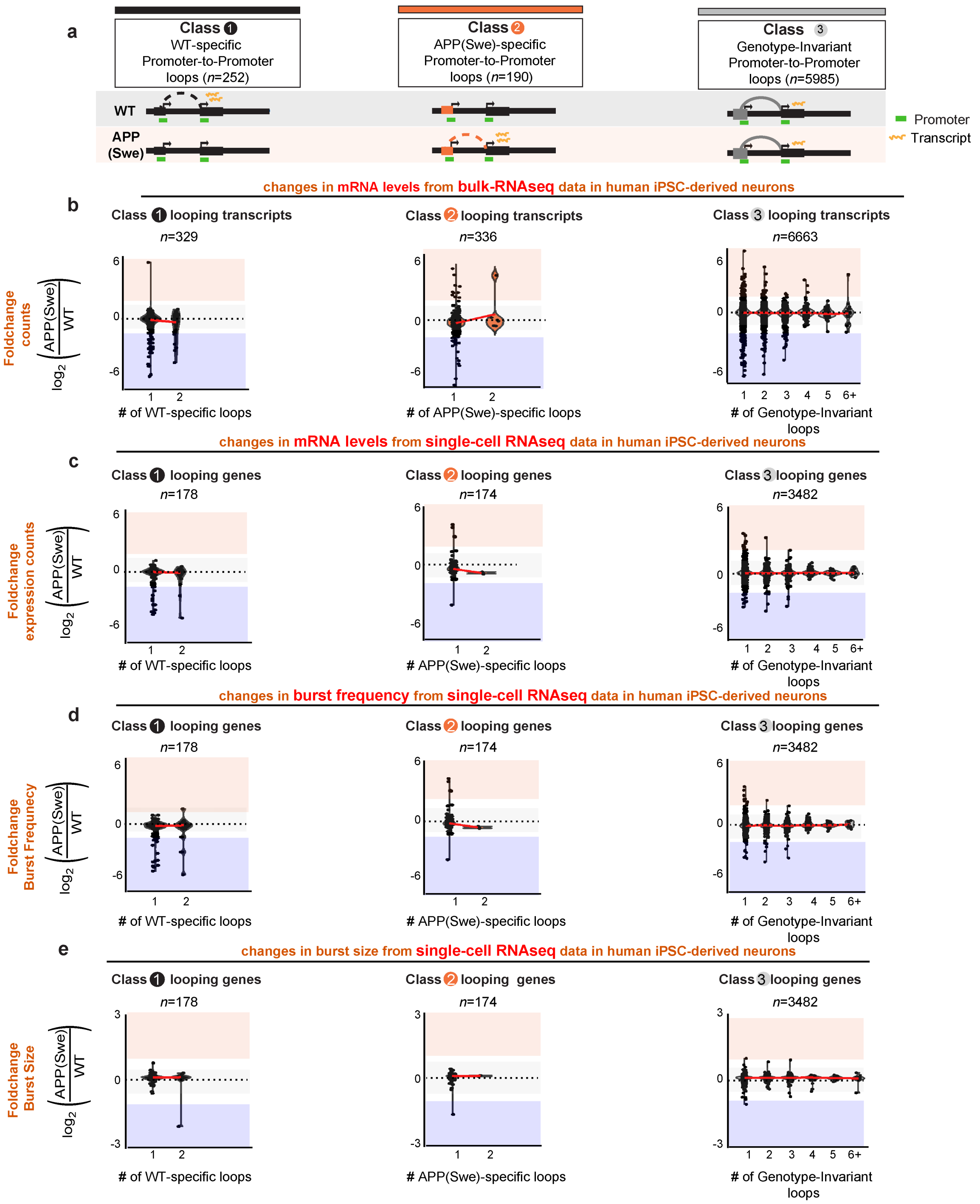


**Supplementary Figure 10: The number of broken Promoter-to-Promoter loops does not correlate with gene expression dysregulation in WT and APP(Swe) human iPSC-derived neurons. (a)** Cartoon representation of transcripts in WT-specific (class 1), APP(Swe)-specific (class 2), and genotype-invariant (class 3) Promoter-to-NonPromoter loops. **(b)** Strip plot of the fold change in bulk RNA-seq mRNA levels stratified by the number of genotype-dynamic or -invariant Promoter-to-NonPromoter loops formed per transcript. **(c-e)** Strip plot of the fold change in three single-cell RNA-seq metrics stratified by the number of genotype-dynamic or -invariant Promoter-to-NonPromoter loops formed per transcript. **(c)** mean single-cell RNA-seq normalized count per gene (log2(APP(Swe)/WT)). **(d)** mean single-cell burst frequency per gene (log2(APP(Swe)/WT)). **(e)** mean single-cell burst size per gene (log2(APP(Swe)/WT)). Red line, median.

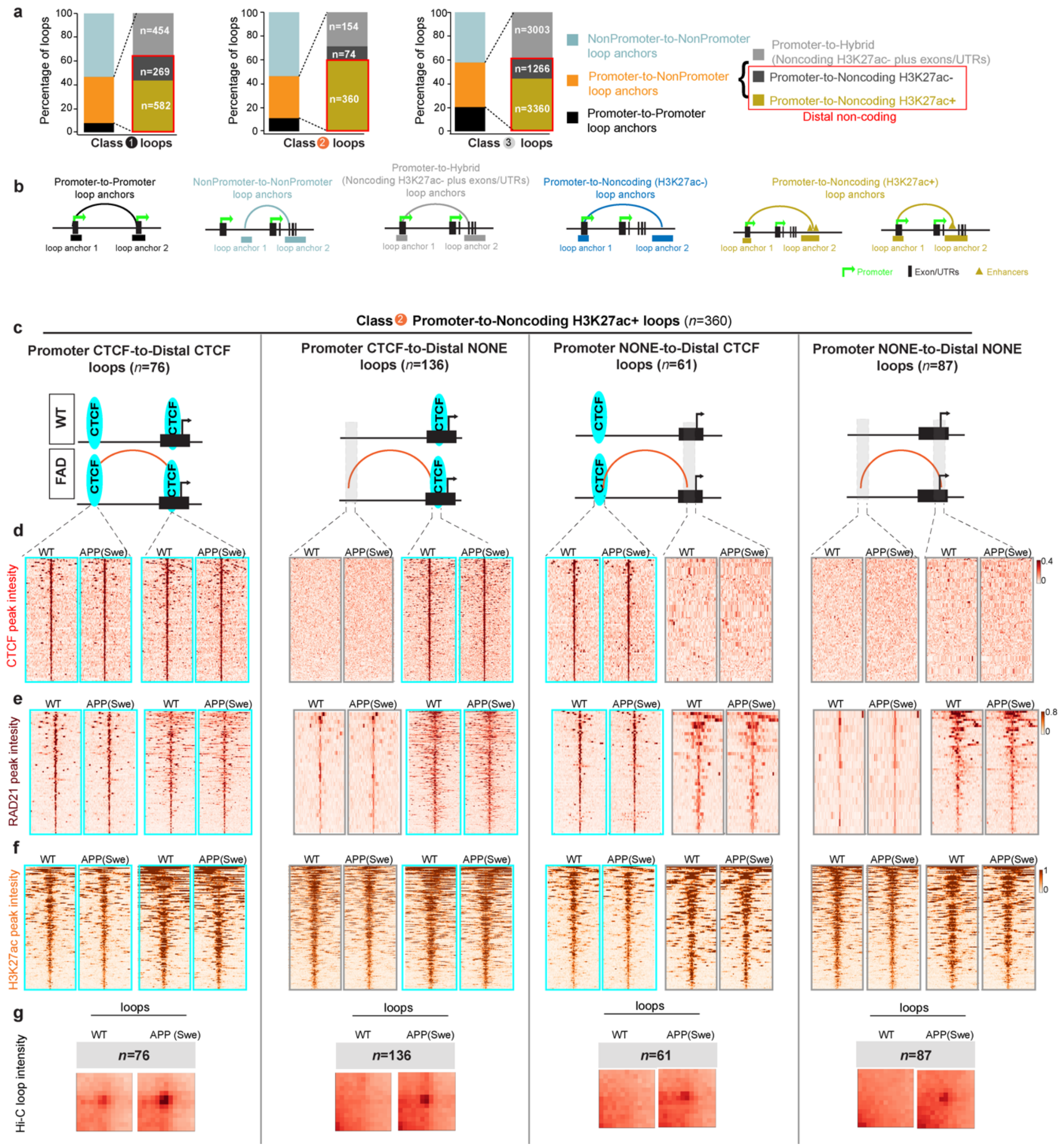


**Supplementary Figure 11: Enhancer-promoter loops gained in FAD-mutant neurons are anchored by unchanging occupancy of the classic architectural proteins CTCF and cohesin. (a)** Stacked barplot showing the stratification of WT-specific (class 1), APP(Swe)-specific (class 2), and genotype-invariant (class 3) loops into NonPromoter-to-NonPromoter, Promoter-to-Promoter, and Promoter-to-NonPromoter subclasses. The Promoter-to-NonPromoter subclass was further substratified into those NonPromoter regions containing noncoding H3K27ac+ peaks characteristic of putative enhancers and those devoid of H3K27ac signal. **(b)** Schematic representation of each of the loop types as in (a). (**c**) Stratification of APP(Swe)-specific (class 2) Promoter-to-Noncoding H3K27ac-positive(+) loops into those anchored by (i) Promoter CTCF-to-Distal CTCF, (ii) Promoter CTCF-to-Distal NONE, (iii) Promoter NONE-to-Distal CTCF, and (iv) Promoter NONE-to-Distal NONE. (**d-f**) ChIP-seq heatmaps of (**d**) CTCF occupancy, (**e**) RAD21 occupancy and (**f**) H3K27ac signal in WT and APP(Swe) in human iPSC-derived neurons. (**g**) Aggregate peak analysis (APA) of the APP(Swe)-specific loop Hi-C interaction frequency in WT and APP(Swe) in human iPSC-derived neurons.

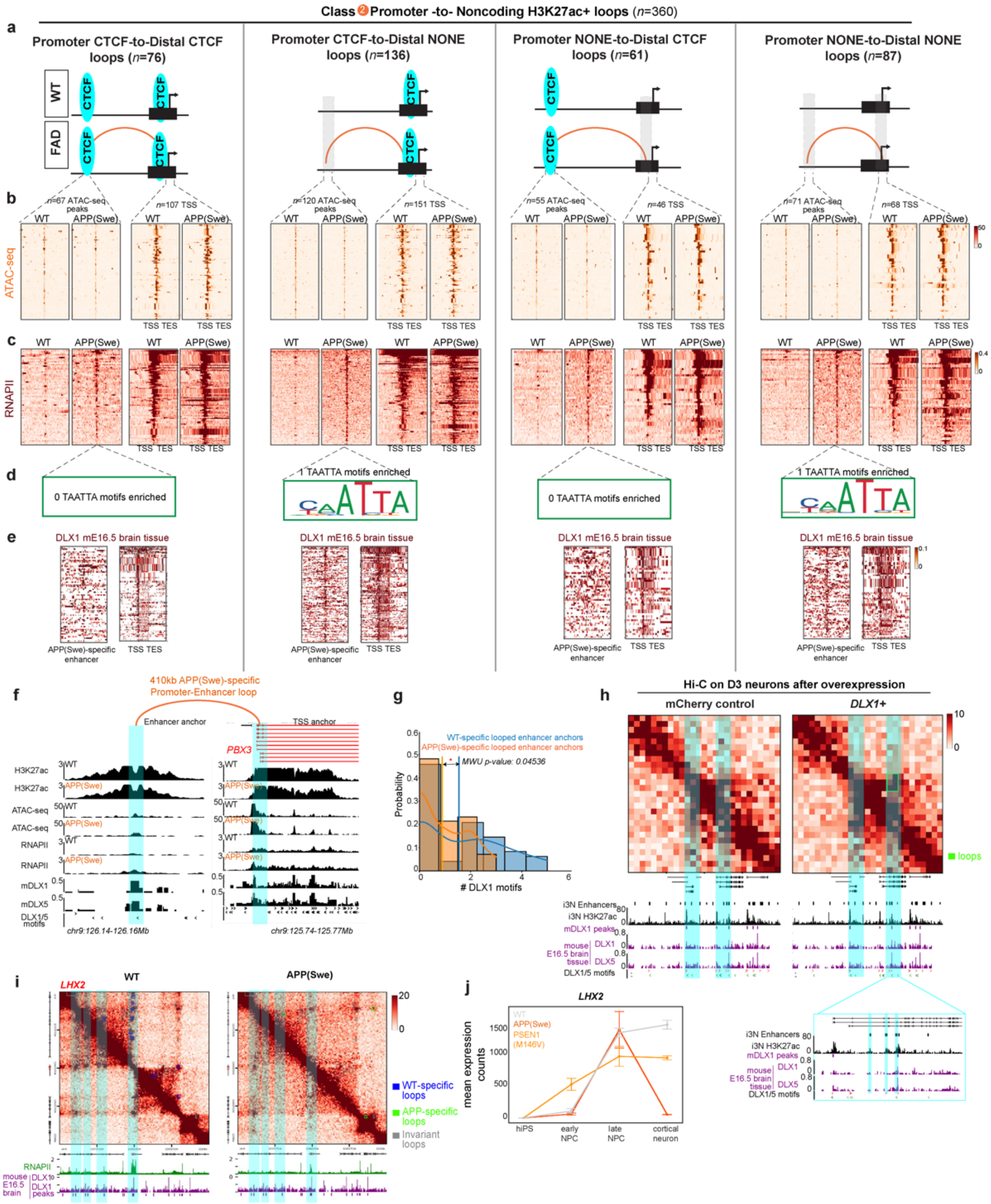


**Supplementary Figure 12: Gained RNA polymerase II signal at single TAATTA motifs at APP(Swe)-specific promoter-enhancer loops gained in FAD-mutant human neurons.** (**a**) Stratification of APP(Swe)-specific (Class 2) Promoter-NoncodingH3K27ac-positive(+) loops into those anchored by (i) Promoter CTCF-to-Distal CTCF, (ii) Promoter CTCF-to-Distal NONE, (iii) Promoter NONE-to-Distal CTCF, and (iv) Promoter NONE-to-Distal NONE. (**b**) Heatmaps of ATAC-seq signal at APP(Swe)-specific promoter-enhancer loops in WT and APP(Swe) human iPSC-derived neurons. (**c**) RNAPII ChIP-seq signal at APP(Swe)-specific promoter-enhancer loops in WT and APP(Swe) human iPSC-derived neurons. (**d**) Logograms of single TAATTA motifs slightly enriched at enhancers in APP(Swe)-specific promoter-enhancer loops after MEME motif analysis (Supplementary Methods). (**e**) Heatmaps of DLX1 ChIP-seq signal from mouse E16.5 brain tissue at distal non-coding H3K27ac+ anchors at APP(Swe)-specific promoter-enhancer loops (**Supplementary Methods**). (**f**) Locus-specific example of an enhancer and promoter loop anchor for an APP(Swe)-specific promoter-enhancer loop anchored by an enhancer with APP(Swe)-specific RNAPII, 1 TAATTA motif, and DLX1/5 chromatin occupancy signal.

(**g**) Probability distribution of the number of TAATTA motifs at WT-specific enhancers anchoring WT-specific promoter-enhancer loops and at APP(Swe)-specific enhancers anchoring APP(Swe)-specific promoter-enhancer loops. Orange and Blue lines indicate *mean*. (**h**) 20kb resolution Hi-C heatmaps at a representative locus in mCherry control, and DLX1+ overexpression conditions. Each track underneath the heatmap corresponds to: H3K27ac+ non-coding putative enhancers and H3K27ac ChIP-seq from D35 glutamatergic i3Ns, DLX1/5 peaks called and DLX1, DLX5 ChIP-seq track in mouse E16.5 brain tissue, DLX1/5 positive and negative direction motif track. (**i**) Hi-C heatmaps at *LHX2* locus in WT, and APP(Swe) cortical neurons. Bin size, 10 kb. **(j)** Mean normalized expression levels of *LHX2* (NM_004789.4) in bulk RNA-seq data for iPSC, early NPC, late NPC, and cortical neurons in WT, PSEN1(M146V) and APP(Swe) genotypes. Error bars represent standard deviation with n = 3 biological replicate experiments.

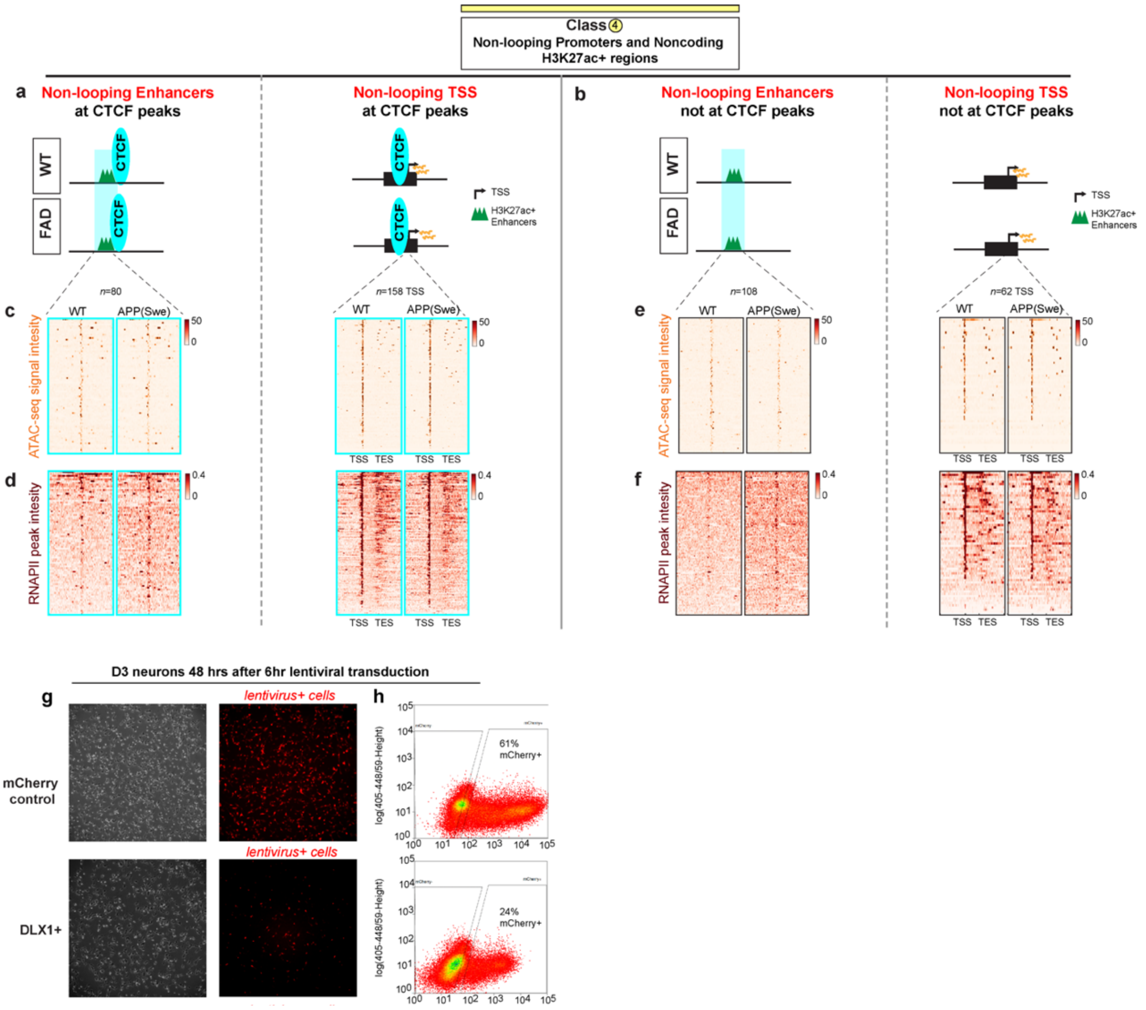


**Supplementary Figure 13: Non-looping mRNA transcripts and non-looping putative H3K27ac+ enhancers in human iPSC-derived neurons do not show changes in ATAC-seq signal or RNA Polymerase II occupancy between WT and APP(Swe) genotypes. (a-b)** Schematic showing non-looping mRNA transcripts and non-looping distal non-coding H3K27ac+ peaks **(a)** co-localized with CTCF peaks and **(b)** devoid of CTCF occupancy. **(c-f)** ChIP-seq heatmaps of (**c, e**) ATAC-seq signal and (**d, f**) RNA Polymerase II (RNAPII) occupancy in WT and APP(Swe) in human iPSC-derived neurons for **(c-d)** non-looping mRNA transcripts and **(e-f)** non-looping putative H3K27ac+ enhancers. (**g**) Representative phase contrast and fluorescence images upon lentiviral transduction of mCherry control, and DLX1-mCherry vectors into i3Ns after 48hours. (**h**) FACS gating used to sort for mCherry+ cells in mCherry control and DLX1 overexpression conditions.

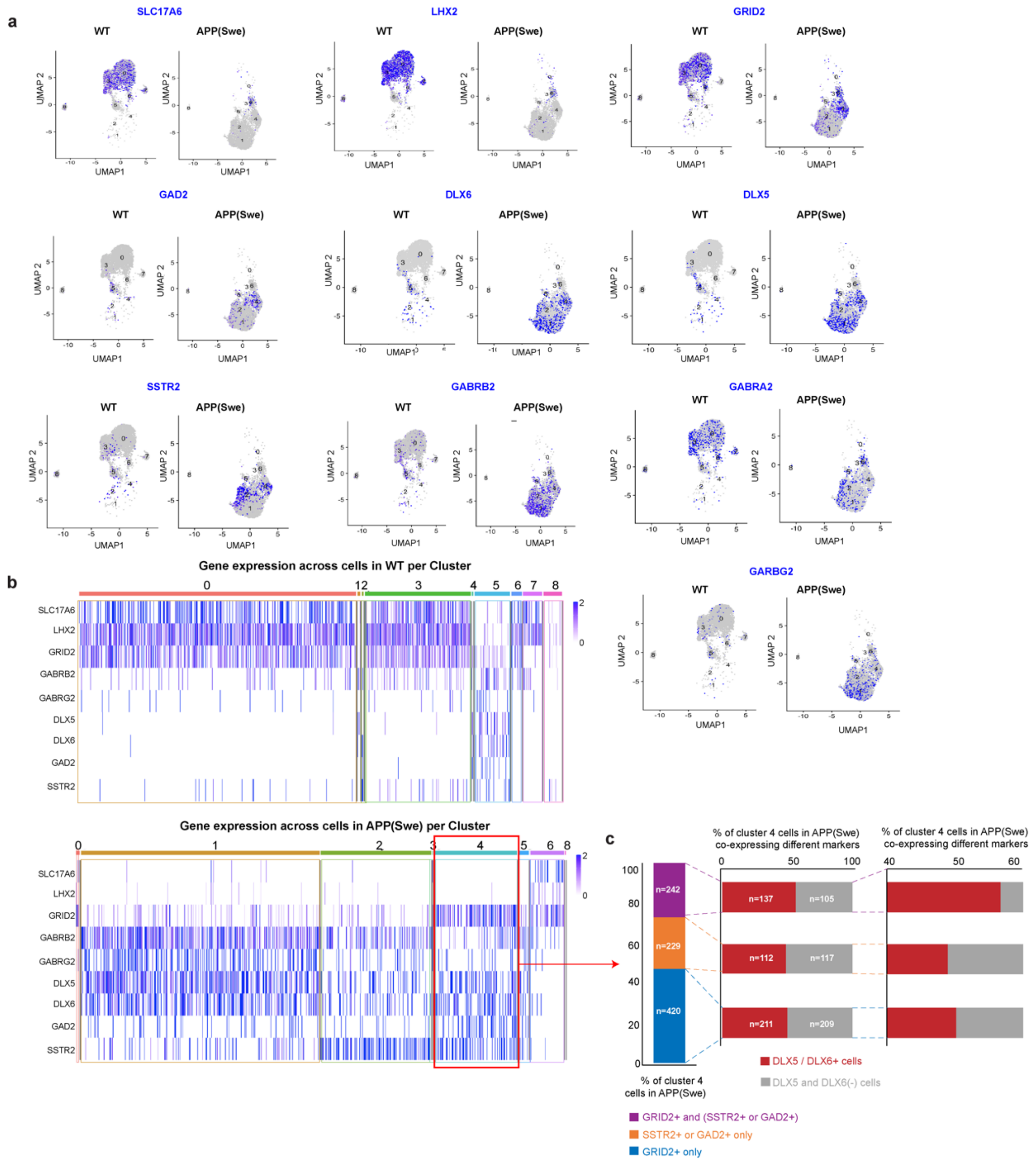


**Supplementary Figure 14: Single-cell analyses reveal a shift in excitatory-to-inhibitory gene expression profiles and presence of a hybrid cluster expressing both excitatory and inhibitory markers in FAD-mutant neurons. (a)** Uniform manifold approximation and projection (UMAP) visualization representing 5,539 cells in WT and APP(Swe) iPSC-derived neurons scRNA-seq data with the expression patterns of genes *SLC17A6*, *GRID2*, *LHX2*, *DLX5*, *DLX6*, *GAD2*, *SSTR2*, *GABRB2*, *GABRA2* and *GABRG2* across all the clusters. (**b***)* Heatmap of gene expression across all single-neurons in WT (*top*) and APP(Swe) (*below*) separated by clusters. (**c**) (*left*) Stacked bar plot of percentage of cells in APP(Swe) Cluster 4 expressing excitatory neuron marker *GRID2* only (*blue*), inhibitory neuron markers *SSTR2* or *GAD2* only (*orange*), and co-expressing both *GRID2* and *SSTR2/GAD2* (*purple*). (*right*) Stacked bar plot indicating the percentage of neurons in each of the groups in (**c**, *left*) further subdivided by co-expression of *DLX5/DLX6* in the same single neurons.

#### Supplementary Methods

##### Human iPSC tissue culture and characterization

We obtained CRISPR/Cas9-engineered human induced pluripotent stem cell (iPSC) lines previously established and validated to carry rare homozygous APP(Swe) and PSEN1(M146V) Alzheimer's disease mutations in an isogenic non-diseased (WT) background[1]. The WT line SA3.5 was derived as a single-cell clone from the 7889SA iPSC line, which as previously reported was derived from commercially available AG07889 male fibroblasts (Coriell Institute), and also served as parent line for CRISPR engineering[1]. Coriell Institute collected the AG07889 fibroblast cells under IRB approval and patient informed consent. We expanded the WT, APP(Swe), and PSEN1(M146V) iPSC lines upon arrival and generated frozen stocks at early passage for early passage-matched experiments.

We coated 10 cm tissue culture plates (Corning, #430167) with 6 mL hESC-Qualified Geltrex (Thermo Fisher Scientific, #A1569601) for at least 1 hour at 37 °C. We thawed and maintained all iPSC lines in StemFlex culture media (Thermo Fisher Scientific, #A3349401) containing 1 % (v/v) penicillin-streptomycin (Thermo Fisher Scientific, #15140122) at 37 °C and 5 % $CO_2$. To passage, we detached cells from plates using 4 ml of Versene (Thermo Fisher Scientific, 15040066) and placing the plates in 37 °C incubator for 3 minutes before inactivation by adding 10 ml of culture media and seeding cells onto freshly prepared Geltrex-coated plates. We split cells every 2-4 days in a 1:3 to 1:8 dilution and passaged cells only up to 3 times prior to experiments.

To verify our human iPSC lines, we routinely characterized the early passage stock vials for (1) karyotype abnormalities, (2) morphology expected from pluripotent cells, (3) presence of proteins used as markers for pluripotent cells, (4) mycoplasma contamination, and (5) confirmation of successful genome editing. To characterize pluripotency markers, we used immunofluorescence staining and microscopy for OCT4 (Santa Cruz, SC9081) and SSEA4 (R&D, MAB1435) as described under "Immunocytochemistry and microscopy (2D culture)". To confirm absence of mycoplasma contamination, we used the universal mycoplasma detection kit (ATCC, 30-1012K). Briefly, we scraped $10^5$ iPSCs into the existing culture media and processed the samples following the manufacturer's instructions.

To assess karyotype of the iPSC lines, we used the Human Karyotype Panel (NanoString, XT-CSO-KAR15-012), containing 338 probes across the 24 human chromosomes. After phenol:chloroform and ethanol precipitation-based extraction of DNA from the iPSCs, we submitted DNA samples to the NanoString service provided by the

Genomics Facility at The Wistar Institute, Philadelphia, PA. Reference value for the karyotype assessment was established by averaging signal intensity of each probe across three karyotypically normal WT iPSC lines (CS0002iCTR, CS0179iCTR, CS0702iCTR), obtained from the Cedars-Sinai Biomanufacturing Center. For each probe, a copy number estimate was calculated by dividing the counts obtained from individual cell lines by the averaged counts of the reference value.

To verify the presence of homozygous APP(Swe) and PSEN1(M146V) point mutations, we used PCR amplification coupled with Sanger sequencing of the PCR product. We performed phenol:chloroform and ethanol-based DNA extraction and precipitation and used 50 ng DNA as an input for PCR. We used Phusion High-Fidelity DNA Polymerase (NEB, M0530S) as recommended by the manufacturer for PCR amplification of the targeted sequences using the following primers:

APPswe_R (ATCCTATAGGCAAGCATTGTATTTTA)
APPswe_F (GGGTAGGCTTTGTCTTACAGTGTTAT)
PSEN1_M146V_F (GGTGAGTTGGGGAAAAGTGA)
PSEN1_M146V_R (CTGGCATTACACATGCACCT)

We purified the amplicons using a MinElute PCR Purification Kit (Qiagen, Cat. No. 28004) and submitted samples for Sanger sequencing at Genewiz (South Plainfield, NJ).

Upon successful confirmation of karyotypically normal iPSCs homogeneously expressing proper pluripotency markers, we continued to conduct (1) daily visual assessment to ensure the expected pluripotent colony morphology and (2) intermittent staining for pluripotent markers.

**iPSC differentiation to DIV65 neurons (2D culture)**

We differentiated iPSC-derived cortical neurons as established and detailed previously with some minor modifications[1-3]. The experiments are again described in detail here to ensure reproducibility given we implemented wet lab procedures as trained by the Paquet lab. The specific details below have also been provided in the published recent references of the authors and should be expected to be similar due to reproducibility of the methodological steps[1-3].

One day (24 hrs) prior to starting differentiation, we replaced StemFlex media with Essential 6 (E6) media (Thermo Fisher Scientific, A1516401) on iPSCs growing on 10 cm tissue culture plates. To generate early neural precursor cells (eNPCs), we detached iPSCs by adding 4 ml Accutase (Thermo Fisher Scientific, A1110501) and triturated to single cell suspension. We added 16 ml of E6 media to inactivate Accutase, counted cells, and centrifuged for 4 minutes at 200 x g. We resuspended cells at a 1 million cell/ml cell concentration in E6 media supplemented with ROCK inhibitor (Stem Cell Technologies, #72304) and seeded iPSCs on flat bottom 12-well tissue culture plates (Falcon, #353043) pre-coated with 0.75 ml Geltrex hESC-Qualified Reduced Growth Factor Basement Membrane Matrix (Thermo Fisher Scientific, A1569601). When cells were 100% confluent, we replaced medium with neural induction (NI) medium (235 ml Neurobasal media (Thermo Fisher Scientific, #21203-049), 235 ml DMEM/F12 GlutaMax (Thermo Fisher Scientific, #10565-018), 10 ml Penicillin-Streptomycin (Thermo Fisher Scientific, #15140-122), 5ml B27 supplement (Thermo Fisher Scientific, #17504-044), 5 ml GlutaMax supplement (Thermo Fisher Scientific, #35050) 5 ml Non-Essential Amino Acid (Thermo Fisher Scientific, #11140-050), 2.5 ml N-2 supplement (Thermo Fisher Scientific, #17502048), 0.25 ml of 500µg/ml Insulin (Sigma, #I1882), 450 uL of 2-mercapto-ethanol (Thermo Fisher Scientific, #21985-023)) supplemented with SB431542 (Selleckchem, #S1067, 1:1000), and LDN-193189 (Selleckchem, #S2618, 1:10000). We counted the day of this first media change as day *in vitro* 0 (DIV0) and maintained cells in NI media for 8 days (DIV8), changing media daily. Throughout the manuscript, cells were harvested at DIV8 for the condition of eNPCs.

To generate late neural precursor cells (lNPCs), we dissociated DIV8 eNPCs using Accutase (Thermo Fisher Scientific, #A1110501) and resuspended at 30 million cells per ml in NI medium supplemented with ROCK inhibitor. We seeded cells onto dried poly-L-ornithine (Sigma-Aldrich, #P4957) and laminin-coated (Thermo Fisher Scientific, #23017015, 20x dilution) 6-well plates in 250-µl spots. We left cells within the spots to adhere for ~45 minutes and then we added 2 ml NI medium supplemented with ROCK inhibitor. On DIV10 we replaced NI media with neural maintenance (NM) media (235 ml Neurobasal media (Thermo Fisher Scientific, #21203-049), 235 ml DMEM/F12 GlutaMax (Thermo Fisher Scientific, #10565-018), 10 ml Penicillin-Streptomycin (Thermo Fisher Scientific, #15140-122), 5 ml B27 supplement (Thermo Fisher Scientific, #17504-044), 5 ml GlutaMax supplement (Thermo Fisher Scientific, #35050) 5 ml Non-Essential Amino Acid (Thermo Fisher Scientific, #11140-050), 2.5 ml N-2 supplement (Thermo Fisher Scientific, #17502048), 0.25 ml of 500µg/ml Insulin (Sigma, #I1882), and 450µl 2-mercapto-ethanol (Thermo Fisher Scientific, #21985-023)). Upon the appearance of neural rosettes, we added 20 ng ml FGF-2 (Stem Cell Technologies Inc., #78003) for 2 days. Upon the emergence of neurons, we manually isolated rosettes under the microscope after 1 hr treatment with STEMdiff Neural Rosette Selection Reagent (Stem Cell Technologies Inc., #5832), washed them, and re-plated on poly-L-ornithine/laminin-coated 6-well plates. Throughout the manuscript, cells were harvested at DIV35 upon the confirmation of homogeneous rosettes for the condition of lNPCs.

To generate terminally differentiated post-mitotic neurons, we further cultivated lNPCs or froze them down between DIV30 and DIV36 in NM media containing 10% DMSO. After trituration of lNPC rosettes into single cells, we seeded ~200,000–500,000 lNPCs on 24-well poly-L-ornithine/laminin-coated plates and cultured in Neurobasal medium containing B-27 serum-free supplement (Thermo Fisher Scientific, #17504044), 2 mM GlutaMax (Thermo Fisher Scientific, #35050061), 100 U per ml penicillin, and 100 microgram per ml streptomycin. During the first week after plating, we treated cells with 10 μM γ-secretase inhibitor DAPT (Sigma-Aldrich, #D5942) and 5-FU to promote neuronal maturation. After DIV43 we maintained cells by replacing half of the media to fresh media every second day. Throughout the manuscript, post-mitotic cortical neurons were harvested at DIV65 and confirmed for homogenous morphology and neuronal markers as described below.

**Amyloid-β measurements**

We measured secreted Aβ monomers ($Aβ_{1-38}$, $Aβ_{1-40}$, and $Aβ_{1-42}$) from cell supernatants collected from human iPSC-derived 2D neurons or 3D organoid cultures using MSD Human Aβ V-PLEX kits (Meso Scale Discovery, #K15200G) as recommended by the manufacturer. This measurement was conducted utilizing the Meso Scale Discovery MESO QuickPlex SQ 120 instrument at the Translational Core Lab facility at the University of Pennsylvania. For the analysis of growth factor-induced human iPSC-derived neurons, we collected 2 biological replicates of media conditioned on each genotype for 24 hours. We stored the supernatants and cell pellets at −80 °C until further processing. To normalize Aβ levels, we measured total protein levels from iPSC and neuron lysates using the BCA assay (Pierce, #23227). To analyze secreted monomers from cortical organoids, we incubated organoids in 150 uL of media for 24 hours and pooled media from 3 size-matched organoids per condition. We did not perform normalization to total protein levels for the cortical organoid samples.

**Thawing DIV35 iPSC-derived NPCs for differentiation to DIV65 post-mitotic neurons**

Thawed cells from frozen cultures at the DIV35 stage of differentiation were plated on polyornithine/Laminin coated plates by resuspending in Neural Maintenance Medium as described[1, 2]. Briefly, we plated approximately 5 million cells onto one well of a pre-coated 6-well plate. After plating, we allowed the cells to recover for a day after thawing and replaced with Neural Maintenance Medium supplemented with 1:5000 bFGF at 48 hours post-thaw. We then changed media daily for five days until NPCs reached full confluency, at which point we dissociated cells using Accutase and plated cells onto 12-well plates containing sterilized 100 mm glass coverslips coated with poly-l-ornithine and laminin, at a density of 200,000 cells per well. 24 hours after replating, we replaced media with 1:1000 DAPT and 1:1000 5-Fluorouracil (Sigma Aldrich, F6627-1G) - Uridine (Sigma Aldrich, U3750-1G) (5FU) (5 mM in cell culture grade water). We then maintained cells by changing half volume of the media three days a week (remove 900 uL, add 1 mL media) with NB/B27 with 1:1000 DAPT/ 1:1000 5-FU. After the sixth feed, we fed cells with half media changes with NB/B27 every 2 days.

**Immunohistochemistry and microscopy (2D culture)**

Using cellular preps from the continuous iPSC to DIV65 neuronal differentiation protocol, we performed immunofluorescence analysis of iPSC markers (OCT4 and SSEA4), NPC markers (NESTIN, FOXG1), and cortical neuronal markers (CTIP2, SATB2) at Day0, DIV8, DIV35, and DIV65. Briefly, we fixed cells in 4% paraformaldehyde (Sigma-Aldrich, #F8775), and permeabilized cell membranes using PBS/0.1% Triton X-100 prior to staining with primary and secondary antibodies. We used anti-OCT4 (Santa Cruz, #SC9081), anti-SSEA4 (R&D, #MAB1435), anti-Nestin (R&D System, #MAB1259), anti-FoxG1 (Abcam, #ab18259), anti-CTIP2 (Abcam, #ab18465), and anti-SATB2 (Abcam, #ab51502) as primary antibodies. As secondary antibodies, we used Goat Anti-Mouse IgG Alexa Fluor® 488 (Abcam, #ab150113), Goat Anti-Rat IgG Alexa Fluor® 647 (Abcam, #ab150159), and Goat Anti-Rabbit IgG Alexa Fluor® 594 (Abcam, #ab150080)). We imaged cells on a Leica DMi8 using software Leica LAS X and processed the images in ImageJ and Adobe Photoshop by altering brightness and contrast levels equivalently across all images to give higher quality phase-contrast images. Parameters were selected after ensuring that secondary antibody only controls did not show signal.

Using cellular preps from the thawed DIV35 to DIV65 neuronal differentiation protocol, we immunostaining for classic amyloid and tau features. At DIV65, 30 days after replating cells onto coverslips, we aspirated media and washed cells with room temperature 1xPBS (Fisher, 20-040-CV) twice to remove dead cells. We diluted 16% paraformaldehyde (Thermo 28908) to 4% in 1xPBS and added to cells for 12 minutes, after which we washed cells with 1xPBS three times, incubating for five minutes each time. We then simultaneously blocked and permeabilized cells for 45 minutes using Blocking Buffer (0.3 % Triton X-100 solution (Sigma, 93443), 5% BSA in 1xPBS). During blocking, we prepared primary antibody solutions in Antibody Buffer (0.3% Triton, 1% BSA in 1xPBS). To stain phospho-Tau we used anti-pTau (AT8, Thermo Fisher Scientific, MN1020, 1:50 dilution, mouse) and to show dendritic morphology we used anti-MAP2 (Abcam, #ab5392, 1:1000 dilution, chicken). To stain amyloid plaques and cell morphology we used anti-amyloid (4G8, BioLegend, #800704,1:1000 dilution, mouse) and anti-MAP2, respectively. After dilution, we centrifuged primary antibody solutions for 5 minutes at 11,600 x g to remove precipitates. After blocking and permeabilization, we aspirated media, rinsed permeabilized the cells with PBS, and added the primary antibody solutions. We then incubated cells overnight at 4C in a humidified chamber. The next day, we prepared secondary antibody solutions in Antibody Buffer, containing. 1:1000 dilution of Goat Anti-Mouse IgG Alexa Fluor 488 (Abcam, ab150113) and Goat Anti-Chicken IgY Alexa Fluor 555 (ThermoFisher, A-21437). We removed primary antibody solutions, and washed cells with 1xPBS three times for 5 minutes. We added secondary antibody solutions and incubated cells for two hours in the dark at room temperature. We then aspirated the secondary antibody buffer and washed cells with 1xPBS 3x5 minutes. We then mounted the coverslips onto microscope slides using VECTASHIELD Antifade Mounting Medium with DAPI (Vector Laboratories, H-1200) and sealed them with nail polish, and imaged slides on a Lecia DMi8 microscope as described above.

**Total RNA-seq from DIV65 iPSC-derived neurons**

We conducted total RNA-seq as we have detailed previously with some minor modifications[1]. The experiments are again described in detail here to ensure reproducibility and should be expected to be similar due to reproducibility of the methodological steps[1].

We isolated total RNA from 3 biological replicates of iPSCs, eNPCs, lNPCs, and neurons from WT, APP(Swe), and PSEN(M146V) genotypes using the Direct-zol total RNA isolation kit (Zymo, #R2061) following the manufacturer's recommendation. The RNA isolated using this approach included non-coding and coding RNA. We assessed purity and integrity of the RNA samples using Agilent RNA 6000 Pico reagent kit on the Bioanalyzer 2100 (Agilent Technologies, Santa Clara, CA, USA). We used 500 ng of isolated RNA for RNA-seq library preparation and after ribosomal RNA depletion we used the RNA for double stranded cDNA preparation using 0.8 U of SuperScript II RT (Thermo Fisher Scientific, #4376600) and A-tailing end repair. We generated strand-specific RNA libraries using the TruSeq Stranded Total RNA LT sample preparation kit (Illumina, #RS-122-2301). To enable multiplex sequencing, we ligated TruSeq RNA Single Indexes to cDNA (Illumina, #20020492). We performed size selection for ~300 bp fragments and performed a clean-up using 42.5 uL of cDNA and 42 uL of Agencourt AMPure XP magnetic beads (Beckman

Coulter, #A63881). To amplify the purified cDNA and enable index incorporation during library amplification, we performed a PCR with 15 cycles. We purified the PCR product using Agencourt AMPure XP beads (Beckman Coulter, #A63881) and assessed library quality using the Agilent DNA 1000 reagent kit (Agilent, 5067–1504) on the Agilent Bioanalyzer 2100 (Agilent Technologies, Santa Clara, CA, USA). We used the Qubit high sensitivity RNA assay kit (Thermo Fisher Scientific, Q32852) on the Qubit Fluorometer to determine the quantity. We used Illumina NextSeq500 platform and the 150-cycle v2.5 High Output kit with 2 x 75bp, paired-end reads for sequencing the pooled libraries.

**iPSC differentiation into organoids (3D culture)**

To confirm that the genomic observations we observe are reproducible in both 2D and 3D cultured neurons, we generated 3D human cortical organoids from WT and homozygous APP(Swe) iPSCs as described previously[4, 5]. Over several months, we transitioned and habituated low-passage stocks of undifferentiated iPSCs to mTeSR Plus (STEMCELL Technology, #05825) media according a published gold-standard protocol (https://www.stemcell.com/how-to-transition-hpscs-into-mtesr-plus-introduction.html). We expanded all iPSC lines on Matrigel-coated plates (Sigma, #CLS354277-1EA) at low passage number, confirmed homogeneity of pluripotency morphology and markers, and established independent Matrigel/mTeSR-adapted iPSC lines that we specifically used for organoid differentiation.

To generate 3D organoids, we expanded undifferentiated human WT and homozygous APP(Swe) iPSCs by seeding cells grown on a 6-well plate onto a 10 cm culture dish and cultured them until reaching 80–90% confluency. On Day -1, we dissociated iPSC colonies into a single-cell suspension with 4 ml Accutase. We added 6 ml of EB formation media (Essential 6™ Medium, Life Technologies, #A1516401) with Y-27632 (10 uM, Selleckchem, #S1049) and, after counting and dilution, we seeded 3 million cells/1ml EB formation media with Y-27632 to one well of a 24-well AggreWell plate (StemCell Technologies, #34815). At Day 0, after 1 day in culture, we transferred embryoid bodies from the microwells to ultra-low attachment 100 mm plates (Corning, #3262) in EB media (E6™ medium containing 2.5 µM Dorsomorphin (Sigma, #P5499) and 10 µM SB-431542 (R&D Systems, #161410)). We replaced EB media daily until neural induction on Day 6.

We induced neural differentiation at Day 6 by changing media to Neural Medium (Neurobasal-A media (Thermo Fisher Scientific, #10888022) containing B-27 supplement without vitamin A (50x) (Thermo Fisher Scientific, #12587010) and GlutaMax in 1:100

dilution) with EGF2 (20ng/ml, R&D, #236-EG-200), FGF-2 (20 ng/ml, R&D, #233-FB-500). We changed media daily for the first 10 days and then every other day for the subsequent 9 days. To promote neural differentiation from Day 25, we added NT-3 (20 ng/ml, Peprotech, #450-03) and BDNF (20 ng/ml, Peprotech, #450-02) to the Neural Medium and removed the growth factors FGF2 and EGF2 and changed media every other day until Day 43. From Day 43, we kept the growing cortical organoids in Neural Medium without the growth factors, BDNF and NT-3 and changed media every 4 days until the harvest time points of Days 100 (D100) and 200 (D200).

**NeuN+ nuclear sorting from cortical organoids**

We performed fluorescent activated nuclei sorting (FANS) and flow cytometry as previously described with modifications[6]. We collected and pooled 6 organoids per replicate (at D100 and at D200) into 1.5 ml Eppendorf tube per replicate and flash froze tissue in liquid nitrogen followed by storage at -80C until nuclear isolation. We prepared a sucrose cushion solution (for 50mL, we used 45 ml 2M sucrose, 4.45 ml H2O, 0.5 ml 1M Tris-HCl pH8.0, 50 µl 3M MgAc2 and 1× protease inhibitor cocktail (Roche)) and poured 14 ml into centrifuge tube (Beckman Coulter, #344058) on ice for each sample. We added 8 ml homogenization buffer (0.32 M sucrose, 10 mM Tris-HCl, (pH 8.0), 5 mM $CaCl_2$, 3 mM $MgAc_2$, 100 microM EDTA, 0.1% v/v Triton X-100, and 1× protease inhibitor cocktail (Roche)) slowly onto the top of the sucrose cushion, collected organoids into 1.5 ml Eppendorf centrifuge tubes, and removed media.

To begin nuclear isolation, we homogenized organoids by adding 1 ml of homogenization buffer with 10ul of RNase inhibitor (Promega, #N2615). We triturated organoids to single cell suspension using sequential pipetting with 1 ml, 200 µl, and 100 µl pasteur pipette tips. We then transferred the 1ml of homogenized sample atop of the 8 ml homogenization buffer + 14 ml sucrose cushion. We washed the Eppendorf tubes with 1ml homogenization buffer and added atop of the 8+14+1 ml. Finally, we also added 8 ml homogenization buffer atop of the 8+14+2ml sample, reaching a total volume of 32 ml.

To isolate nuclei using the sucrose cushion, we performed ultracentrifugation at 25,700 rpm for 120 min at 4°C in a SW43Ti swinging-bucket rotor (Acceleration 6 and Deceleration 6). After removing most of the supernatant (leaving ~100-300 µL), we added 10µL of RNasin (Promega RNasin Plus RNase Inhibitor, #N2615) and 1.5 ml wash buffer (19 ml 1xPBS, 1 ml 10% BSA, 1× protease inhibitor cocktail (Roche)) on ice for 20 min. We then triturated nuclei with a 1 mL capacity mechanical pipette for 30-60 s until clumps were visibly reduced and removed a 10 µL aliquot of each sample for nuclei counting. We spun the remaining nuclei for 15 min at 5,000 rpm on a tabletop centrifuge to obtain a loosely packed pellet of nuclei. We resuspended nuclei in wash buffer containing RNasin (~7-10 million nuclei/mL) for 20 min by rotating the tubes at 4°C, followed by incubation of nuclei with primary antibody for 40 min at 4°C while rotating in the dark. Primary antibody solutions included: (1) anti-NeuN antibody conjugated with Alexa Fluor 488 (Millipore, MAB377x, 1:1000) diluted in wash buffer and (2) Alexa Fluor 488 only diluted in wash buffer as a 'no antibody' negative control. We also added 1µL of DAPI (Sigma, #D9542-1MG) per mL for 5 min at 4°C while rotating in the dark. Following primary antibody incubation, we spun samples for 10 min at 5000 rpm at 4°C, washed for 10-20 min at 4°C by rotating the tubes in the dark, and then subjected samples to a final spin for 10 min at 500 g at 4°C. Finally, we resuspended the samples at 5 million nuclei per 1 mL of FACS buffer (1xPBS, 1% BSA, 1× protease inhibitor cocktail (Roche)) with 10 µL of RNasin ribonuclease inhibitor.

To prepare neural nuclei for sorting, we filtered samples into a polystyrene holding tube (BD Falcon 5mL tubes w/cell-strainer cap, #352235). We used a Moflo Astrios sorter with a 100 µm nozzle and Summit software to perform sorting. We gated on the FSC-Area (forward scatter, proportional to size) and SSC-Area (side scatter, proportional to cell complexity) parameters to remove debris, followed by gating on SSC-Height and SSC-width as well as FSC-Height and FSC-width to select single nuclei. Finally, we gated on DAPI+ and NeuN+ signal compared to the "Secondary antibody only" negative control and collected ~120,000 NeuN+ nuclei per replicate directly into 1 mL TRIzol.

**Immunohistochemistry and microscopy (3D organoid characterization)**

We analyzed differentiation markers, including the canonical ventricular zone (SOX2), cortical layer (TBR1, CTIP2), pan-neural (MAP2), astrocyte (GFAP), and mature cortical neuron (CTIP2, SATB2) markers by immunofluorescence staining at organoid D50, D100, and D200. We fixed organoids in 4% paraformaldehyde (Thermo Fisher Scientific, #28908) by placing samples at 4°C for 15 min. Following the fixation step, we washed organoids three times with 1xPBS, incubating the samples at room temperature after each wash for 10 min. We transferred organoids into 30% sucrose solution and kept overnight at 4°C. Following the removal of the sucrose solution, the organoids were equilibrated with O.C.T (optimal cutting temperature) compound (Sakura, #4583) at room temperature for 15 min. We then transferred organoids to tissue base molds and embedded within O.C.T compound. We stored the organoid blocks at -80°C until cryosectioning 15 µm slices.

To prepare cryosectioned slides for immunostaining, we washed 3 times with 1xPBS and permeabilized and blocked the slices by incubating in PBS/0.1% Triton X-100 containing Normal Goat Serum Blocking Solution (Vector Laba, #S-1000). We stained the samples with primary antibodies (anti-MAP2 (Abcam, #ab5392, 1:1000 dilution), anti-GFAP (Invitrogen, #130300, 1:100 dilution), anti-amyloid (4G8, BioLegend, #800704, 1:1000 dilution), anti-pTau (AT8, Thermo Fisher Scientific, MN1020, 1:50 dilution), anti-SOX2 (Cell Signaling Technologies, #3579S, 1:200 dilution), and anti-TBR1 (Abcam, #ab31940, 1:200 dilution) for overnight at 4°C and secondary antibodies (Goat Anti-Mouse IgG Alexa Fluor® 488 (Abcam, #ab150113, 1:1000 dilution), Goat Anti-Rat IgG Alexa Fluor® 647 (Abcam, #ab150159, 1:1000 dilution), Goat Anti-Rabbit IgG Alexa Fluor® 594 (Abcam, #ab150080, 1:1000 dilution)) for 1 hour. We used aa TCS SP8 Multiphoton Confocal (Leica TCS SP8 Multiphoton Confocal) microscope and Leica LAS X software for imaging. We processed images in ImageJ/FIJI and Adobe Photoshop, where we altered brightness and contrast levels equivalently across all images to give higher quality phase-contrast images. Parameters were selected after ensuring that secondary antibody only controls did not show signal.

**Total RNA-seq from NeuN+ nuclei sorted from organoids**

We conducted total RNA-seq as we have detailed previously with some minor modifications[1]. The experiments are again described in detail here to ensure reproducibility and should be expected to be similar due to reproducibility of the methodological steps[1]. We isolated total RNA from ~120,000 NeuN+ nuclei sorted from a pool of 6 organoids per genotype and time point as described above. We sorted N=2 separate populations of ~120,000 NeuN+ nuclei from WT and APP(Swe) D100 organoids and N=1 population of ~120,000 NeuN+ nuclei from WT and APP(Swe) D200 organoids. We isolated total RNA using the Direct-zol total RNA isolation kit (Zymo, #R2061) as recommended by the manufacturer and assessed purity and integrity of the RNA samples using Agilent RNA 6000 Pico reagent kit on the Bioanalyzer 2100 (Agilent Technologies, Santa Clara, CA, USA). We used NEBNext® rRNA Depletion Kit v2 (NEB, #E7405L) to deplete ribosomal RNA and the NEBNext® Ultra™ II Directional RNA Library Prep kit (NEB, #E7765S) for library preparation. To facilitate multiplex sequencing, we ligated cDNA to unique indexes using the NEBNext Ultra II DNA Library Prep (NEB, #E7645S). We amplified adaptor-ligated products for 15 cycles and performed size selection and magnetic bead clean-up. We sequenced pooled samples on an Illumina NextSeq500 instrument using a 150-cycle (2 x 75) v2.5 High Output kit.

**Single-cell RNA-seq from growth factor-induced iPSC-derived neurons**

We obtained a suspension of viable single cells from iPSC-derived neurons at the same time point as bulk genomics (DIV65) by incubating 1 well of a 6-well plate (~2 million neurons) with 2ml Accutase. We assessed viability of the obtained cell suspensions by taking a 10 µl aliquot of cell suspension and mixed with 2x Trypan blue. All processed samples contained >90% viable cells. We sequenced 6,000-12,000 neurons per genotype using the V3 10x 3' RNA-Seq kit (10x Genomics, PN-1000075) and sequenced libraries on HiSeq 4000 (Illumina). We used the service at PSOM Next-Generation Sequencing Core, University of Pennsylvania (Philadelphia, PA, USA).

**Neuron sample fixation for Hi-C and ChIP-seq**

To fix neurons for downstream ChIP-seq (CTCF, RNA Polymerase II, H3K27ac, RAD21) and Hi-C assays, we followed previously described protocols[7-15]. Briefly, we fixed cells by incubating in fixation media (1 % (v/v) formaldehyde in DMEM/F-12 (Thermo Fisher Scientific, #11320033)) for 10 min at room temperature. We diluted fixation media from 11% formaldehyde solution (100 mM NaCl (Thermo, AM9760G), 50 mM HEPES-KOH , pH 7.5 (Hampton Research, HR2-729), 1 mM EDTA (Invitogen, 15575-038), 0.5 mM EGTA (BioWorld, 40520008-1), 37% formaldehyde (Sigma, F8775-25mL)). We quenched the fixation process by adding 125 mM glycine (Sigma Aldrich, 50046-250G) and incubating at room temperature for 5 min, followed by a 15 min incubation at 4°C. We washed the cross-linked cells in 1x PBS before flash freezing and storage of pellets at −80 °C.

**ChIP-seq**

We prepared CTCF, RNA Polymerase II (RNAPolII), H3K27ac, and RAD21 ChIP-seq libraries as previously described with minor modifications[7-15].

For CTCF, RNAPolII, and H3K27ac, we crosslinked cell pellets (8-10 million cells) as described above. 12 h prior to lysing pellets, we incubated 20 µg anti-RNAPolII (ActiveMotif 39097), 10 µg anti-CTCF (Millipore 07-729), or 10 µg anti-H3K27Ac (ab4729) with 20 uL Protein A (Thermo Fisher, 20333) and 20 uL Protein G agarose beads (Thermo Fisher, 20398) in 1 mL Phosphate Buffered Saline (PBS) without calcium & magnesium (Corning 21-040-CV) by rotating at 10 rpm at 4 °C until use. We resuspended pellets in pre-chilled cell lysis buffer (10 mM Tris pH 8.0 (Invitrogen, #AM9856), 10 mM NaCl (Invitrogen, #AM9760G), 0.2 % (v/v) NP-40 (Sigma, #I8896), 1 mM phenylmethanesulfonyl fluoride (PMSF; Sigma,

#93482), 0.2 % (v/v) Protease inhibitor cocktail (Sigma, #P8340)) and incubated on ice for 10 min. We homogenized cells using a dounce homogenizer and 30 strokes with pestle A.

To pellet and solubilize nuclei, we centrifuged lysates at 2,500 x g for 5 min at 4°C and resuspended in 500 µl nuclear lysis buffer (10 mM EDTA (Invitrogen, #15575020), 50 mM Tris (pH 8.0), 1 % (w/v) SDS (Invitrogen, #15553027), 1 mM PMSF, 0.2 % (v/v) Protease inhibitor cocktail) on ice for 20 min before mixing with 300 µl IP dilution buffer (150 mM NaCl, 20 mM Tris (pH 8.0), 2 mM EDTA, 1 % (v/v) Triton X-100 (Sigma, #93443), 0.01 % (w/v) SDS, 1 mM PMSF, and 0.2 % Protease inhibitor cocktail). Next, we sheared chromatin to ~200-600 bp DNA fragment size using Qsonica Q800R3 (Qsonica Sonicators, CT) with parameters of 100% amplitude and 30 sec on / 30 sec off pulses. After sonication we centrifuged lysates at 16,000 x g and added pre-clearing solution (50 µg of IgG (Sigma, #I8140), 175 µl of Protein A Agarose suspension (Thermo Fisher Scientific, #15918014), and 175 µl of Protein G Agarose suspension (Thermo Fisher Scientific, #15920010) in 3.7 ml of pre-chilled IP dilution buffer and 0.5 ml of nuclear lysis buffer) to each supernatant. We incubated the samples by rotating at 10 rpm for 2 hrs at 4°C.

We next performed target protein-associated chromatin immunoprecipitation from supernatant using antibody-bound beads by rotating the samples at 10 rpm overnight at 4°C. Approximately 24 hours later, we washed beads once in IP wash buffer 1 (50 mM NaCl, 20 mM Tris (pH 8.0), 2 mM EDTA, 1 % (v/v) TritonX-100, 0.1 % (w/v) SDS), followed by two washing step in high-salt buffer (500 mM NaCl, 20 mM Tris (pH 8.0), 2 mM EDTA, 1 % (v/v) TritonX-100, 0.01 % (w/v) SDS), one washing step in IP wash buffer 2 (250 mM lithium chloride (Sigma, #L9650), 10 mM Tris (pH 8.0), 1 mM EDTA, 1 % (v/v) NP-40, 1 % (w/v) sodium deoxycholate (Sigma, #D6750)), and two washing steps in 1 x TE (Fisher, #BP2473500) at 4 °C. We eluted chromatin from beads in elution buffer (1 % (w/v) SDS, 100 mM sodium bicarbonate (Fisher, #S233) and degraded RNA by adding RNaseA (Roche, #10109169001) at 65°C for 1 hr. To remove residual proteins and reverse crosslink DNA, we added 2.4 U of Proteinase K (NEB, #P8107S) to eluent and incubated at 65°C overnight. We purified ChIP DNA using conventional phenol:chloroform-based extraction and ethanol precipitation methods and stored ChIP DNA at -20 °C until library preparation. We used 5 ng of purified ChIP-seq DNA for downstream library preparation as described below.

For RAD21 ChIP-seq, we performed similar step as described above but with minor modifications to cell membrane lysis. Briefly, we crosslinked cell pellets (8-10 million cells) as described above and stored in -80 °C. 12 h prior to lysing pellets, we incubated 6 µg Rad21 antibody (ab992) with 20 uL Protein A (Thermo Fisher, 20333) and 20 uL Protein G agarose beads (Thermo Fisher, 20398) in 1 mL Phosphate Buffered Saline (PBS) without calcium & magnesium (Corning 21-040-CV) by rotating at 10 rpm at 4 °C until use. We then resuspended pellets in pre-chilled cell lysis buffer (10 mM Tris pH 8.0, 10 mM NaCl, 0.2 % (v/v) NP-40, 1 mM PMSF, 0.2 % (v/v) Protease inhibitor cocktail (Sigma, #P8340)) on ice, rotated at 10 rpm at 4 °C for 20 min, centrifuged at 2,500 x g for 5 min at 4°C, and resuspended the pellet in fresh pre-chilled cell lysis buffer. We homogenized cells using a dounce homogenizer and 630 strokes with pestle A. We performed all following steps as described above for CTCF, H3K27ac, and RNAPolII ChIP, then used 1 ng of purified ChIP-seq DNA for downstream library preparation.

**Hi-C**

We conducted Hi-C on ~2 million cross-linked cells per replicate per condition using a commercial Hi-C kit (Arima Genomics, Inc., #A510008) following the manufacturer's recommended protocol. Following restriction digest of the chromatin with multiple enzymes, we filled in the 5'-overhangs to label the digested ends with a biotinylated nucleotide. We ligated spatially proximal digested ends of DNA and purified the proximity-ligated DNA. To prepare samples for library preparation, we sheared purified proximity-ligated DNA fragments to an average size of 300 to 400 bp using a Covaris S220 sonicator with the following setup: 200 cycles per burst for 55 seconds at 140 W peak incident power and 10% duty factor. We performed size-selection to obtain 200-600 bp DNA fragments using AMPure XP DNA Purification Beads (Beckman Coulter, #A63881). We enriched ligation junctions with incorporated biotin-tag after size-selection using Enrichment Beads provided in Arima-Hi-C kit, and stored streptavidin beads containing enriched DNA fragments up to 3 days before proceeding with library preparation as described below.

**Library preparation (ChIP-seq, Hi-C)**

We conducted genomics library preparation as we have detailed previously with some minor modifications[1]. The experiments are again described in detail here to ensure reproducibility and should be expected to be similar due to reproducibility of the methodological steps[1].

We prepared sequencing libraries using the NEBNext Ultra II Library Prep Kit (NEB, #E7645S) as recommended by the manufacturer, with minor modifications. In brief, for ChIP and Hi-C experiments, we end-repaired and dA-tailed DNA following the manufacturer's protocol. For Hi-C libraries, we washed twice the adaptor-ligated Hi-C libraries on streptavidin beads, first in 150 µl of wash buffer at 55 °C and then next in 100 µl of elution buffer (Arima

Genomics, #A510008), and then eluted ligation products from streptavidin beads by heating samples to 98°C and boiling for 10 min in 15 µl elution buffer. For ChIP-seq libraries, we size-selected adaptor-ligated libraries using AgenCourt Ampure XP beads (Beckman Coulter, #A63881) and then amplified using NEBNext Ultra II DNA Library Prep Kit for Illumina (NEB, #E7645S) as recommended by the manufacturer. For ChIP-seq, DNA fragments <1 kb size were size-selected and amplified using 7-8 PCR cycles. After additional purification using AgenCourt Ampure XP beads (Beckman Coulter, #A63881), we assessed the quality of the individual libraries using Agilent Bioanalyzer High Sensitivity DNA Analysis Kits (Agilent, #5067-4626) and quantified DNA concentration using a Kapa Library Quantification Kit (KAPA Biosytema, #KK4835). We pooled libraries and sequenced on an Illumina NextSeq 500 instrument using 75 bp single-end reads for ChIP-seq and 37 bp pair-end for Hi-C.

**ATAC-seq**

To generate ATAC-seq libraries, we harvested ~50,000 cells following Accutase treatment, cell counting, and centrifugation at 500 g x 5 min at 4°C. We first washed the cells with 1xPBS and resuspended the pellets in 50 µL of lysis buffer (10 mM Tris-HCl (pH7.4), 10 mM NaCl, 3 mM MgCl2, and 0.1% IGEPAL CA-630). We collected cell pellets by centrifugation, discarded the supernatant, and immediately proceeded with the transposition reaction using 50 µL transposition reaction mix. We prepared transposition reaction mix using components of the Nextera DNA Library Prep kit (FC-121-1030). For 50 µl reaction, we mixed 25 µl TD (Tagment DNA Buffer), 2.5 µl TDE1 (Tagment DNA Enzyme, Tn5 Transposase) and 22.5 µl nuclease-free H2O) at 37°C for 30 min. After transposition steps, we performed DNA clean-up, using a QIAGEN MinElute Kit and stored the eluted DNA at −20°C until generating the sequencing library.

To generate sequencing libraries from the transposed and purified DNAs, we thawed the sample to carry out an initial 5-cycle PCR (1 cycle: 72°C for 5 min; 1 cycle: 98°C for 30 s; then 5 cycles: 98°C for 10 s, 63°C for 30 s, 72°C for 1 min) in a 50 µL of pre-mixed reaction including NEBNext High Fidelity 2X Master mix and 5 µL of 25 µM Forward/Reverse ATAC-seq index per sample. To reduce introduction of size bias and GC-content bias in the PCR step, we used a previously described method to monitor the PCR reaction using qPCR and stop amplification before saturation. Briefly, to determine the optimum number of additional cycles of PCR amplification, we used a 5 µL aliquot of the PCR reaction for qPCR quantification. We plot linear Rn (fluorescent signal value) versus cycle number to determine the cycle number that corresponds to 1/3 of the maximum fluorescent intensity. We ran PCR reactions at ~5-7 cycles as determined by qPCR. We purified each PCR reaction with AgenCourt Ampure XP beads (Beckman Coulter, #A63881) and eluted the PCR products in 12 µL elution buffer. We verified the size of the fragments (target 200-2000 bp) with an Agilent High Sensitivity DNA chip. All libraries were pooled for downstream sequencing on an Illumina NextSeq 500 instrument using 75 bp paired-end reads.

**Cell culture for DLX1 neuron overexpression experiments**

We cultured commercially available i3N iPSCs (WTC11.G3-WT, Gladstone) on Matrigel-coated plates (Sigma, #CLS354277-1EA) in mTeSR Plus (STEMCELL Technology, #05825) media. These cells contain a transgene cassette consisting of tetracycline transactivator rtTA3G under the control of the CAG promoter and Neurogenin 2 under the control of the TRE3G promoter stably inserted at the AAVS1 locus, to rapidly differentiate iPSCs to neurons with the addition of doxycycline to the media. We passaged cells using Versene (Thermo Fisher Scientific, 15040066) to disassociate colonies at 37 °C for 5 minutes followed by neutralization in mTesr Plus media. We centrifuged the cells at 250 x g for 3 minutes and split cells in a 1:3 to 1:8 dilution. We expanded cells onto 15cm dishes for the DLX overexpression experiments and when cells reached 40% confluence (DIV-2), we initiated neural induction by including 1ug/mL doxycycline (Sigma, #D3447-500MG) in the mTesR Plus media to activate the Neurogenin 2 transgene. We prepared plates for neural cultures on DIV-2 and DIV-1 by coating with poly-L-ornithine and laminin as described under *'Thawing DIV35 iPSC-derived NPCs for differentiation to DIV65 post-mitotic neurons,'* and pre-warmed plates containing one-half culture volume of iN media at 37 °C prior to re-plating the cells on DIV0. At DIV-1, we replaced media with fresh mTeSR Plus media supplemented with 1µg/mL doxycycline. On DIV0, we washed cells with 1xPBS, dissociated cells with Accutase (Thermo Fisher Scientific, #A1110501) for 5 min, and neutralized Accutase with mTeSR Plus. We centrifuged cells for 3 min at 250 x g and resuspended cells in iN media (Neurobasal media (Thermo Fisher Scientific, #21203-049), GlutaMax (Thermo Fisher Scientific, #35050061), pen/strep (Thermo Fisher Scientific, #15140122), B27 supplement (Thermo Fisher Scientific, #17504-044), 10 ng/mL BDNF (Peprotech, 450-02), 10ng/mL NT-3 (Peprotech, 450-03)) supplemented with 1ug/mL doxycycline, 10uM DAPT (Selleckhem, S2215), 5uM 5-Fluorouracil (Sigma Aldrich, F6627-1G)- Uridine (Sigma Aldrich, U3750-1G) (5FU). We gently triturated cells to a single cell suspension and passed cells through a 70 µm strainer. We then plated cells at a concentration of 78,000/$cm^2$ on the prepared coated plates for the DLX overexpression experiments. We performed a full media change at DIV1 with iN media supplemented with 1ug/mL doxycycline and 1:500 dilution of lentiviral particles containing transgenes encoding DLX1 or mCherry. We aspirated media containing virus after 6 hours and replaced it with fresh iN media supplemented with 1ug/mL doxycycline.

**Lentivirus production for DLX1 neuron overexpression experiments**

We received synthetic full length hDLX1 (ENST00000361725.5) in pUC-GWAmp vectors from Azenta/GENEWIZ including additional KOZAK sequence, HA-tag and 2x NLS and BamHI and BsiWI restriction sites for direct subcloning. After digestion with BamHI (R0136S, NEB) and BsiWI (R3553S, NEB) we used Instant Sticky-end Ligase Master Mix (NEB, M0370S) to subclone the synthetic genes into pAW91.EF1a.mCherry vector consisting of EF-1-alpha promoter and in-frame an mCherry-coding sequence with T2A tag. To produce third generation self-inactivating lentiviruses, we plated HEK293T cells onto 15 cm dishes 24 hrs before transfection in Dulbecco's Modified Eagles Medium (DMEM) (Corning, 10-013-CV) supplemented with 10% fetal bovine serum (FBS) (Atlanta biologicals, S11550) and 1 % (v/v) penicillin-streptomycin (Thermo Fisher Scientific, 15140122). When cells reached 40-60% density, we transfected cells with 2.4 microgram psPAX2 (Addgene: #12260), 4.4 microgram pMD2.G (Addgene: #12259) and 7.5 microgram pAW9.1-hDLX1-FL, or pAW91-mCherry no insert plasmids using FuGENE 6 reagent (Promega, E2693) using 3:1 transfection reagent to DNA ratio. We changed media 12 hours after transfection to 15 mL fresh media. We collected viral supernatant 48 hours after media change from the virus-producing cell lines and centrifuged to remove cells and debris. We mixed the 12 mL media with 4 mL of Lenti-X Concentrator (Cat. Nos. 631231) and incubated for 4 hours at 4°C. We centrifuged the mixture at 1,500 x g for 45 minutes to obtain a high-titer virus-containing pellet which we resuspended in 1xPBS and aliquoted for subsequent transductions. We stored aliquoted lentivirus stocks at -80°C.

**Fluorescence Activated Cell Sorting (FACS) for DLX1 overexpression experiments**

At 48 hours post-transduction, we dissociated cells from the dish by washing with 1xPBS, applying Accutase (Thermo Fisher Scientific, #A1110501) for 6 minutes and gentle scraping with a cell lifter (Corning Incorporated, 3008). We neutralized Accutase with Neurobasal Media (Thermo Fisher Scientific, #21203-049) and centrifuged the cells for 3 minutes at 250xg. We resuspended cells in FACS buffer (1xPBS, 0.5% Bovine Serum Albumin (Sigma, A8806-5G), Protease Inhibitor (Sigma, A8806-5G), and RNase Inhibitor (Thermo, AM2696)) to a concentration of four million cells/mL for sorting. We set gates on the MoFlo Astrios Cell Sorter (Beckman Coulter) using Fluorescence Minus One controls, untreated cells to assess background red fluorescence, and mCherry transduced cells to assess positive transduction. We recovered 61% positive cells in the mCherry condition, and 24% positive cells in the DLX condition. For Hi-C libraries, we sorted one million mCherry, or DLX positive and negative singlets per sample into FACS buffer. Each of the constructs used in the overexpression lead to varying degrees of brightness for mCherry. Therefore, to accurately assay the percentage of cells for each condition that received the virus, images were independently thresholded in ImageJ to the following values: mCherry: minimum displayed value 2000, maximum displayed value 12,000; DLX: minimum displayed value 2600, maximum displayed value 4,000.

**Hi-C for DLX1 overexpression experiments**

After sorting, we fixed the cell suspension at a cell concentration of one million cells/mL for 10 minutes by the addition of 11% formaldehyde to a final concentration of 1% as described under '*Neuron Sample Fixation for HiC and ChIP-seq*. We quenched formaldehyde by adding 2.5 M glycine to a final concentration of 125 mM. We pelleted cells at 1350xg for 5 minutes at 4C, washed with 1 mL 1xPBS, spun down a second time, and flash froze in liquid nitrogen prior to storage at -80C. To perform HiC, we thawed samples on ice for twenty minutes and resuspended in ice cold Lysis Buffer (10mM Tris-HCl pH8.0, 10mM NaCl, 0.2% Igepal CA630 and 1X PIC (Roche)) by pipetting up and down until solution appeared visibly homogenous. We then rotated the sample on a rotator at 4C for twenty minutes, spun down at 600 xg for 5 minutes at 4C, resuspended in 1 mL of lysis buffer, and transferred to a dounce homogenizer. We dounced samples 120 times with Pestle A to fully isolate nuclei and verified successful nuclei isolation by confirming a single-nuclei suspension of intact nuclei using DAPI and fluorescent microscopy.

After isolating nuclei, we pelleted nuclei at 600 x g for 5 minutes at 4C, resuspended in 50 µL of 0.5% SDS, and incubated at 62°C for 10 minutes. We used 145 uL of water and 25 uL of 10% Triton X-100 to quench SDS. To digest the DNA, we added 25 uL of 10X NEBuffer r3.1 (NEB, B6003S), 100 U of DpnII (NEB, R0543S), and 100 U of DdeI (NEB, R0175L) to solution, and digested chromatin overnight at 37C on a rotating heat block set to 850 rpm. The next day, we incubated samples at 65 C for 20 minutes to deactivate restriction enzymes, and then cooled to room temperature. Next, to fill in the fragment overhangs, we added 50 uL of the Biotin Fill-In Master Mix (37.5 uL 0.5 mM biotin-14-dATP (Life Technologies, 19524-016), 1.5 uL 10 mM dCTP, 1.5 uL 10 mM dGTP, 1.5 uL 10 mM dTTP, and 8 uL 5U/uL DNA

Polymerase 1, Large (Klenow) Fragment (NEB, M0210)) to solution. We fully mixed the sample by pipetting with a p200 5 times, and then incubated for 45 minutes with rotation at 850 rpm.

To ligate blunt ends, we added 900 uL of Ligation Master Mix (669 uL water, 120 uL 10X NEB T4 DNA ligase buffer (NEB, B0202), 100 uL 10% Triton X-100, 6 uL 20 mg/mL Bovine Serum Albumin (NEB, B9000S), and 5 uL 400 U/uL T4 DNA Ligase (NEB, M0202) to solution, mixed sample by inverting, and ligated DNA by keeping reactions at room temperature for 4 hours on a nutator with slow rotation. We then reversed crosslinks by pelleting nuclei at 2500 x g for 5 minutes at room temperature and resuspended the pellet in 300 uL 10 mM Tris-HCl, pH8; 0.5M NaCl, 1% SDS solution. 50uL of 20mg/ml proteinase K (NEB, P8107S). We placed samples at 55 C for 5 minutes with rotation at 850 rpm, followed by 55 C for 25 minutes with no rotation, and finally placed the samples at 68 C overnight with no rotation. To degrade RNA, we cooled samples to room temperature, added 5 uL RNase A (Thermo Scientific, EN0531), and incubated at 37 C for 30 minutes.

Finally, we isolated DNA by adding equal volume (350 uL) of phenol chloroform (Thermo Fisher, 15593049), vortexed samples for 30 seconds, and transferred to a phase lock tube (VWR, 2302830). We spun samples down at top speed (21,100 x g) at 4C for 5 minutes and transferred the upper phase to a new DNA low bind tube. To precipitate DNA, we added 35 uL 3 M sodium acetate (Thermo Scientific, R1181), 3 uL glyco blue (Thermo Fisher, AM9515), and 900 uL 100% ethanol, and mixed with 1 mL pipette. We placed samples at -80C for 24 hours, and then spun down at 21,100 x g for 30 minutes at 4C. Upon removal of the supernatant, we washed samples with 800 uL ice cold 70% ethanol in water and spun down again at 21,100 x g for 30 minutes at 4C. We removed the supernatant, air dried samples to allow all ethanol to evaporate, and dissolved in 130 uL Elution Buffer (Qiagen, 19086) at 37C for 10 minutes. We performed DNA shearing, size selection, and biotin enrichment as previously described[16].

**hg38 RefSeq reference transcriptome**

We downloaded our hg38 reference transcriptome on 7th July, 2021 using the UCSC Table Browser with NCBI RefSeq and RefSeq Curated as track and table options respectively (https://genome.ucsc.edu/cgi-bin/hgTables?hgsid=1345872709_tycZ8naeqyTXL51A2BV9FK8CsBk0). From this reference list, we filtered to keep only the canonical chromosomes (chr1-chr22, chrX, and chrY) and removed any duplicate occurrences of a transcript from the chrY if it was also present on chrX. The final reference transcriptome used for all studies had 80,649 transcript annotations.

#### Total RNA-seq analysis for iPSC-derived neurons

We mapped RNA-seq paired-end reads to the hg38 RefSeq reference transcriptome for both cDNA and ncRNA using kallisto (0.46.1) quant with 100 bootstraps of transcript quantification[17]. Following pseudoalignment by kallisto, we converted quantifications of estimated counts into DESeq2 format in R using the library ("tximportData") according to DESeq2 documentation recommendations[18]. We identified differentially expressed transcripts between APP(Swe) versus isogenic control (WT) conditions and PSEN1(M146V) versus isogenic control (WT) conditions at each differentiation stage (iPSC, eNPC, lNPC, neuron) using the DESeq2 (1.22.1) Wald test-statistic and BH adjusted p-value < 0.05. We reported differential genes that passed the p-value threshold and also passed thresholds of abs(log2(fold-change)) 0.32 and median-of-ratios (MOR) normalized count>=300 in at least one of the three genotypes.

We defined "Common FAD DOWN" as transcripts with (1) median-of-ratios (MOR) normalized count>=300 in at least one of the three genotypes in iPSC-derived neurons, (2) a log2[APP(Swe)+0.1/WT+0.1]<-0.32 and log2[PSEN1(M146V)+0.1/WT+0.1] < -0.32, and (3) a significant adjusted p-value of < 0.05 in APP(Swe) vs. WT and PSEN1(M146V) vs. WT pairwise comparisons. We defined "Common FAD UP" as transcripts with (1) median-of-ratios (MOR) normalized count>=300 in at least one of the three genotypes in iPSC-neurons, (2) a log2[APP(Swe)+0.1/WT+0.1]>0.32 and log2[PSEN1(M146V)+0.1/WT+0.1]>0.32, and (3) a significant adjusted p-value of < 0.05 in APP(Swe) vs. WT and PSEN1(M146V) vs. WT pairwise comparisons. We defined transcripts to be "genotype-invariant expressed" in WT and FAD conditions as transcripts with (1) median-of-ratios (MOR) normalized count>=300 in at least one of the three genotypes iPSC-neurons, (2) a -0.32<log2[APP(Swe)+0.1/WT+0.1]<0.32 and -0.32<log2[PSEN1(M146V)+0.1/WT+0.1]<0.32 in iPSC-neurons, and (3) a significant adjusted p-value of >=0.05 in both APP(Swe) vs. WT and PSEN1(M146V) vs WT pairwise comparisons. P-values, estimated counts, MOR normalized counts, and fold-changes are provided for all three genotypes in iPSC-neurons (**Supplementary Table 2)**. Differentially expressed transcripts for each pairwise comparison as well as Common FAD Down and Common FAD Up lists are provided (**Supplementary Table 3)**.

#### Total RNA-seq analysis for NeuN+ nuclei derived from D100 and D200 organoids

We mapped RNA-seq paired-end reads to the hg38 RefSeq reference transcriptome as described above for both cDNA and ncRNA using kallisto (0.46.1) quant with 100 bootstraps of transcript quantification[17]. For D100 organoids, following pseudoalignment by kallisto, we converted quantifications of estimated counts into DESeq2 format in R using the library("tximportData")[18] We identified differentially expressed transcripts between APP(Swe) vs. WT NeuN+ nuclei as: 1) transcripts with median-of-ratios (MOR) normalized counts>=300 in at least one of the genotypes in D100 organoids, (2) a fold-change threshold [APP(Swe)+0.1/WT+0.1]>1.25 for upregulated transcripts and a fold-change threshold of [APP(Swe)+0.1/WT+0.1]<1.25 for downregulated transcripts. For D200 organoids, we normalized estimated counts from kallisto by the median-of-ratios (MOR) method and assessed fold change expression for APP(Swe) vs. WT NeuN+ nuclei [APP(Swe)+0.1/WT+0.1]]. Estimated counts, MOR normalized counts, and fold-changes are provided for WT and APP(Swe) genotypes for NeuN+ neurons derived from both D100 and D200 organoids (**Supplementary Table 4)**. Differentially expressed genes for WT and APP(Swe) genotypes for NeuN+ neurons derived from the D100 condition are provided (**Supplementary Table 4)**.

**ChIP-seq analysis**

We aligned CTCF, RNAPII, RAD21, and H3K27ac ChIP-seq reads for WT, APP(Swe), and PSEN1(M146V) human iPSC-derived neuron conditions to the human (hg38) genome using Bowtie[19] (version 0.12.7). We removed reads with more than two possible alignments using the -m2 flag. We filtered the mapped reads to remove optical and PCR duplicates using samtools[20] (version 1.2) and down-sampled CTCF, RNAPII and H3K27ac ChIP-seq libraries to 24 million reads to achieve equal read numbers across samples. We downsampled RAD21 ChIP-seq libraries to 41 million reads. We identified peaks using MACS2[21] (version 2.1.2) with a p-value cutoff parameter of 1E-4. For CTCF ChIP-seq, we applied default MACS2 parameters. For RAD21, H3K27ac, and RNAPII ChIP-seq, we applied MACS2 broad peak calling approach (--broad --broad-cutoff 1E-4). Peak calls for 3 genotypes and 4 antibodies for iPSC-neurons are provided (**Supplementary Table 14**).

**ATAC-seq analysis**

We aligned ATAC-seq reads for two replicates each in WT, APP(Swe), and PSEN1(M146V) iPSC-neurons to the human (hg38) genome using Bowtie[19] (version 0.12.7) after removing adapters from the fastq files using Cutadapt[22] (version 1.9.1). We removed reads with more than two possible alignments using -m2 flag. We filtered the mapped reads to remove optical,

PCR duplicates and kept only uniquely mapped reads using samtools[20] (version 1.2). To account for the center of Tn5 transposase binding site, we offset all reads aligning to the + strand by +4 bp, and all reads aligning to the – strand were offset by −5 bp. After merging replicates, we downsampled all the libraries to 39 million reads to achieve equal read numbers across samples. We identified peaks for each condition using MACS2[21] (version 2.1.2) with --nolambda --nomodel --keep-dup all --call-summits and a p-value cutoff parameter of 1E-4. Peak calls for 3 genotypes for iPSC-neurons are provided (**Supplementary Table 5**).

**Differential ATAC-seq analysis**

We concatenated ATAC peaks in iPSC-neurons ($P$ value $= 1 \times 10^{-4}$) identified in WT and APP(Swe) conditions and merged any overlapping peaks using BEDTools[23] (v2.29.1) merge. This resulted in a master list of ATAC-seq peaks (N=134158) identified in WT and APP(Swe) iPSC-neurons. We then parsed these peaks into WT-specific or APP(Swe)-specific ATAC-seq classes by (i) calculating the average bigwig signal across the peak interval using the pybigwig package (v0.3.13) in both the WT and APP(Swe) conditions, and (ii) calculating the fold change in ATAC-seq signal as [APP(Swe)/WT]. We assigned a peak as APP(Swe)-specific ATAC-seq peak if it exhibited a fold change ATAC-seq signal as [APP(Swe)/WT]>2.0 and if the APP(Swe) signal at the peak was above 40th percentile signal threshold of all ATAC-seq peaks. We assigned a peak as WT-specific ATAC-seq peak in the same manner with the conditions reversed. The remaining ATAC-seq peaks were classified as Invariant ATAC-seq peaks i.e the peaks with less than two-fold-change in signal between the two conditions and if the signal under the peak was above 40th percentile signal threshold of all peaks in both conditions. We identified a total of 16,385 WT-specific, 10,691 APP(Swe)-specific, and 29,559 genotype-invariant ATAC-seq peaks. (**Supplementary Table 6**). Similarly, we identified differential ATAC-seq peaks between PSEN1(M146V) and WT (**Supplementary Table 6**).

**Classifying TSS±2kb and non-TSS regions of the hg38 genome by ATAC-seq peak signal**

We used BedTools[23] (v2.29.1) subtract to generate a genome-wide hg38 non-TSS regions list that excluded TSS±2kb of all transcripts. Next, we used BedTools[23] (v2.29.1) intersect to identify the number of unique WT-specific, APP(Swe)-specific, and genotype-invariant ATAC-seq peaks that intersected with each of the TSS±2kb and non-TSS regions.

**Hi-C Analysis: Pre-processing**

We aligned 37 bp paired-end Hi-C reads to the hg38 reference genome using bowtie2[24] (version 2.2.5) (global parameters:–very-sensitive –L 30 –score-min L,-0.6,-0.2 –end-to-end–reorder; local parameters:–very-sensitive –L 20 –scoremin L,-0.6,-0.2 –end-to-end–reorder) with the HiC-Pro software (version 2.7.7)[25] (**Supplementary Table 7, tab2**). We filtered non-uniquely mapped reads, PCR duplicates, and unmapped reads and then counted bonafide cis and trans hybrid ligated fragments. After confirming the high quality of Hi-C data with the expected allocation of cis and trans reads **(Supplementary Table 7, tab2)**, by binning paired reads into uniform 10 kb bins, we assembled cis-contact matrices as previously described[11, 14, 15].

After matrix assembly, we applied a mappability filter to remove regions with poor mappability using the hg19 36-mer CRG Alignability track from ENCODE (http://hgdownload.soe.ucsc.edu/goldenPath/hg19/encodeDCC/wgEncodeMapability/wgEncodeCrgMapabilityAlign36mer.bigWig) lifted over to hg38. If the average mappability of a 50 kb window centered on that bin was below 50%, we set the contacts between 10 kb bins to NaN. We filtered rows containing less than 35 non-zero 10kb bins within 750 kb of the diagonal from further HiC analysis. To normalize mappability variation for every row in 10 kb cis contact matrices, we merged replicates and performed Knight-Ruiz matrix balancing for each chromosome individually and for every genotype as previously described[8, 11, 14]. For subsequent loop calling, we retained the final bias factors.

**Hi-C Analysis: Loop calling**

We built algorithms to call loops in Hi-C data genome-wide based on previously published work by our group and others with several modifications (https://bitbucket.org/creminslab/cremins_lab_loop_calling_pipeline_11_6_2021/src/initial/)[8, 11, 12, 14, 16, 26-28]. We have previously described the details of the methods in other manuscripts[1, 11], and they are again described in detail here to ensure reproducibility. Mathematical terms and equations should be expected to be similar due to reproducibility of the methodological steps[1, 11]. We restricted our analysis to $\leq$ 10 Mb of interaction distances between bin-bin pairs for expected modelling and all subsequent loop calling steps. We performed all computations to model the interaction frequency expectation on the merged, Knight-Ruiz balanced contact matrices binned at 10 kb resolution for each of WT, APP(Swe), and PSEN1(M146V) iPSC-derived neurons.

We computed a one-dimensional distance-dependence expected model, $D$, by averaging interaction counts of each of the first 1,000 diagonals **(Equation 1)**[11, 14]:

$$D_d = \underset{(k,l)\in\{(a,b)|b-a=d\}}{\text{gmean}} \left(S_{k,l}\right) \ \forall\, d \text{ such that } 0 \le d \le 1{,}000 \quad \textbf{(1)}$$

Where the expected value is $D_d$ for interaction between two bins separated by $d$ bins, $S$ is the balanced contact matrix, and computed the geometric mean over sets of bin-bin pairs separated by bins $b - a = d$. To avoid the effects of zeros, using a pseudo count of 1, we computed $(X_t)$ as the geometric mean over indices $t$ taken from a set $T$. (**Equation 2**)[11, 14]:

$$\underset{t\in T}{\text{gmean}}(X_t) = \left(\sqrt[|T|]{\prod_{t\in T} X_t + 1}\right) - 1 \quad \textbf{(2)}$$

To estimate the local expected background interaction frequency at each locus we adjusted the one-dimensional distance-dependence $D_d$. We computed a correction factor for bin-bin pairs with interaction distances > 150 kb, using the donut and lower left filters as in Rao et al and our own previous work[8, 11, 12, 14, 16, 26-28]. Mathematically, we represent the donut footprint around bin-bin pair $(i, j)$ as (**Equation 3)**[11, 14]:

$$DF_{i,j} = \{(a,b) \mid (|a-i| \le w) \wedge (|b-j| \le w) \wedge (a \ne i) \wedge (b \ne j) \\ \wedge \left((|a-i| > p) \vee (|b-j| > p)\right)\} \quad \textbf{(3)}$$

where $DF_{i,j}$ represents the set of bin-bin pairs $(a, b)$ that are included in the donut footprint centered on bin-bin pair $(i, j)$, and parameters $p$ and $w$ control the inner and outer radius of the donut shape. For the donut footprint bin-bin pairs were included $(a, b)$ if they lay within a $(2w + 1) \times (2w + 1)$ square centered on $(i, j)$ around bin-bin pair $(i, j)$ unless (i) they fell on the same row or same column as $(i, j)$ or (ii) they lay within a $(2p + 1) \times (2p + 1)$ square centered on $(i, j)$. We used $p = 4$ and $w = 16$ for all the conditions in this analysis. We computed the donut-corrected expected values (**Equation 4**)[11, 14]:

$$E_{i,j}^{DF} = D_{i,j} \times \frac{\sum_{(a,b)\in DF_{i,j}} S_{a,b}}{\sum_{(a,b)\in DF_{i,j}} D_d} \quad \textbf{(4)}$$

where $E^{DF}$ is the matrix of donut footprint corrected expected values. The donut filter summation factor corrects the one-dimensional distance-dependent expected value $D_{i,j}$ at pixel

$i, j$ to account for local enrichment or reduction of balanced values $S_{a,b}$ in the local donut-shaped window relative to the one-dimensional distance-dependent expected values $D_d$. We excluded points that were below the diagonal, had interaction distance greater than 10 Mb or in low-mappability regions.

As proposed by Rao et al.[16], we employed a lower left footprint: (**Equation 5**)[11, 14]:

$$LLF_{i,j} = \{ (a, b) \in DF_{i,j} \mid (a < i) \wedge (b < j) \} \quad \textbf{(5)}$$

where $LLF_{i,j}$ represents the set of bin-bin pairs $(a, b)$ included in the lower left footprint centered on bin-bin pair $(i, j)$. We kept points that are below and to the left of $(i, j)$th pixel in the original donut filter to compute the lower left footprint $LLF_{i,j}$. We next computed lower left corrected expected values as (**Equation 6**)[11, 14]:

$$E_{i,j}^{LLF} = D_{i,j} \times \frac{\sum_{(a,b) \in LLF_{i,j}} S_{a,b}}{\sum_{(a,b) \in LLF_{i,j}} D_d} \quad \textbf{(6)}$$

Finally, we used the maximum of **Equations 4** and **6** as the final corrected expected value for all bin-bin pairs with interaction distances greater than 150 kb.

The sensitivity of loop calling near the diagonal of a contact matrix is lowered due to over-estimation of the donut expected background signal at distances within 150 kb from the diagonal. Therefore, to capture short range interactions, we modeled the on-diagonal (≤ 150 kb) background expected using only the upper-triangle of the donut footprint.We only included in the donut footprint the bin-bin pairs $(a, b)$ that have interaction distances >= the interaction distance of the entry for which the corrected expected value was computed (**Equation 7**)[11, 14]:

$$UTF_{i,j} = \{ (a, b) \in DF_{i,j} \mid b - a \geq j - i \} \quad \textbf{(7)}$$

where the set of bin-bin pairs $(a, b)$ included in the upper triangular footprint centered on bin-bin pair $(i, j)$ is represented by $UTF_{i,j}$.

We then computed the upper triangular corrected expected values using this upper triangular footprint for all bin-bin pairs with interaction distances within 150 kb according to (**Equation 8**)[11, 14]:

$$E_{i,j}^{UTF} = D_{i,j} \times \frac{\sum_{(a,b)\in UTF_{i,j}} S_{a,b}}{\sum_{(a,b)\in UTF_{i,j}} D_d} \quad \textbf{(8)}$$

In summary, we computed our final expected values $E_{i,j}$ for bin-bin pairs within 150 kb interaction distances using the upper triangular corrected expected values and for bin-bin pairs with interaction distances beyond 150 kb using the maximum of the donut and lower left footprints corrected expected values (**Equation 9**)[11, 14]:

$$E_{i,j} = \begin{cases} E_{i,j}^{UTF}, & \text{for } b - a \leq 15 \\ \max\left(E_{i,j}^{DF}, E_{i,j}^{LLF}\right), & \text{for } 15 < b - a \leq 1000 \end{cases} \quad \textbf{(9)}$$

**Loop Calling: Computing P-values, Multiple Testing Correction, and Clustering**

We have previously described the details of the methods for computing P-values, multiple testing correction, and clustering in other manuscripts[1, 11], and they are again described in detail here to ensure reproducibility. Mathematical terms and equations should be expected to be similar due to reproducibility of the methodological steps[1, 11]. To compute a biased expected value, we used the final expected value $E_{i,j}$ and the Knight-Ruiz balanced bias vector $c$ for comparison to the raw read counts $X_{i,j}$ (**Equation 10**)[11, 14]:

$$E_{i,j}^{\text{biased}} = E_{i,j} \times c_i \times c_j \quad \textbf{(10)}$$

We computed P-values against the null hypothesis that the raw read count $X_{i,j}$ was less than or equal to the biased expected value $E_{i,j}^{biased}$ to assemble a matrix of P-values $P_{i,j}$. We computed the probability that the raw read count $X_{i,j}$ was less than or equal to a Poisson-distributed random variable $X'_{i,j}$ with mean $E_{i,j}^{biased}$ (**Equation 11**)[11, 14]:

$$P_{i,j} = P\left(X_{i,j} \leq X'_{i,j}\right); \quad X'_{i,j} \sim \text{Poisson}\left(E_{i,j}^{\text{biased}}\right) \quad \textbf{(11)}$$

To perform multiple testing correction, we applied lambda-chunking strategy[16]. Briefly, with a $2^{1/3}$ bin spacing using logarithmically spaced bins, we first stratified bin-bin pairs $(i, j)$ according to their biased expected values $E_{i,j}^{biased}$. We next applied Benjamini-Hochberg false discovery rate control to obtain q-value matrices $Q_{i,j}$ for the P-values $P_{i,j}$ for each chunk separately. $Q_{i,j}$ represents the maximum false discovery rate (FDR) at which an interaction would be called significant[11, 14].

From the matrix of q-values $Q_{i,j}$ computed above, we identified clusters of nearby significant bin-bin pairs. For a bin-bin pair $(i, j)$ to be called as significant, we required it to pass 3 thresholds: (i) a q-value threshold $Q_{i,j} \leq 0.1$ (corresponding to an FDR of 10%); (ii) a balanced contact value $S_{i,j} \geq 8.0$; and (iii) a fold-change between the balanced and final expected value $\frac{S_{i,j}}{E_{i,j}} \geq 1.5$. To reduce false-positives, we removed clusters with lower than three adjacent significant bin-bin pairs[11, 14].

We further applied a progressively stringent q-value threshold and re-clustered bin-bin pairs that passed the more stringent q-value until at least a three adjacent bin-bin pairs remained in the cluster. The q-value thresholds were applied in order-of-magnitude steps from 0.1 to 1E-20 FDR. To further reduce the possibility of false positive interactions called near the diagonal, we removed all refined clusters whose interaction distance was within 20 kb from the diagonal[11, 14]. The final loops identified in each of the three genotypes is shown in (**Supplementary Table 9**).

**3DeFDR for genotype-specific loop calling**

We applied our recently published 3DeFDR-HiC analysis package[26] to identify genotype-specific looping interactions in APP(Swe) versus WT (two replicates each). To identify which of the identified loops were APP(Swe)-specific or WT-specific, we input: i) raw 10 kb assembled Hi-C matrices per replicate, ii) bias factors obtained from the KR balancing of 10kb matrices per replicate, iii) set of loop calls in APP(Swe) and WT (as described in previous section).

3DeFDR-HiC[13] describes the Hi-C read counts using the negative binomial distribution parameterized in terms of its mean and dispersion. During the initial steps of preparing data in 3DeFDR-HiC, we excluded points that are: i) beyond a distance threshold (>10 Mb) from the diagonal, ii) have zero in all replicates, and iii) rows that failed balancing. We used simple scaling method to determine the size factors for the Hi-C matrices. We then applied the quantile-adjusted conditional maximum likelihood (qCML) method in 3DeFDR-HiC to estimate dispersion at each distance scale and a weighted LOWESS curve to fit the dispersion across all distance scales. Subsequently, for each pixel, we computed a likelihood ratio test (LRT). This LRT compares a null model of no differential interactions (where the true mean parameter is shared across all replicates of WT and APP(Swe)) to an alternative model in which the two genotypes have a different true mean. To obtain q-values, we then applied Benjamini-Hochberg FDR correction on the resulting p-values called via the LRT by considering only those pixels that are involved in the looping interactions. For a looping pixel to be called significantly differential across the two genotypes, we applied a q-value threshold ≤ 0.3 (corresponding to an FDR of 30%). Remaining pixels that did not pass the q-value threshold were classified to be Invariant across the two genotypes i.e are not significantly differential looping pixels.

To assign the looping clusters as either APP(Swe)-specific, WT-specific, or genotype-invariant, we checked the frequency of the differential or invariant pixels within each of the loop clusters. Clusters were classified as follows: 1) if a loop cluster had higher number of WT-specific pixels than the number of APP(Swe)-specific and Invariant pixels, then it was classified to be a WT-specific looping cluster; 2) if the cluster had higher number of APP(Swe)-specific pixels than the number of WT-specific and Invariant pixels, then it was classified to be APP(Swe)-specific looping cluster; and 3) if the number of Invariant pixels were higher than number of APP(Swe)- and WT-specific pixels, or had equal numbers of APP(Swe)- and WT-specific pixels, the cluster was said to be Invariant across the two genotypes. We retained only those clusters that had at least three looping pixels within it. We identified 3,346 WT-specific loops, 1,680 APP(Swe)-specific loops and 29,527 Invariant loops. (**Supplementary Table 10**).

**3DNetMod for TAD/subTAD detection**

We identified TADs and subTADs genome-wide using 3DNetMod [11, 14, 29] as extensively documented and validated and our publicly available code https://bitbucket.org/creminslab/cremins_lab_tadsubtad_calling_pipeline_11_6_2021. We have previously described the details of the methods for detecting TADs/subTADs[1, 11], and they are again described in detail here to ensure reproducibility. Mathematical terms and equations should be expected to be similar due to reproducibility of the methodological steps[1, 11].

We used parameters that we optimized on 10 kb binned, scaled, Knight-Ruiz balanced Hi-C matrices for iPSC-derived neurons in WT, APP(Swe), and PSEN1(M146V) (hg38). Briefly, we log transformed each merged replicate's genome-wide counts data and chunked matrices into 6 Mb regions with 4 Mb overlap. We removed chunked regions from further analysis if they exhibited non-zero counts for ≥ 2/3 of all pixels on the diagonal or exhibited consecutive zero counts on the diagonal for ≥ 500kb. We identified plateaus of consecutive gammas showing the same number of domains (mean per 20 partitions), in 0.01 gamma steps, and set a minimum consecutive gamma plateau size of 20 for 6 Mb chunked regions. A sweep of gamma values was computed as the mean gamma at every plateau. We ran 3DNetMod 20 times (i.e. 20 partitions) and identified the optimal genomic location of domains at each gamma and computed the consensus genomic location of domains from the 20 partitions via the adjusted rand index[11, 14, 29].

We then filtered domains smaller than 130 kb from the genome-wide list and removed domains within 20 bins from the edges of chunked regions and merged to create a concatenated list of all remaining genome-wide domains after filtering steps from both the 6 Mb chunked regions. We merged domains that co-localized +/- 70 kb with both boundaries into a single TADs/subTAD unit with start and end coordinates separated by the largest genomic distance.

We next established a set of unique boundary locations by adjusting boundaries so that TADs/subTADs with similar genomic coordinates shared a single consistent boundary[11, 14, 29]. To achieve this, we iterated through all domains assessing left and right boundaries separately for each domain. If two or more domains come into close contact such that the gap between them is smaller than 7.5% of the domain size for all adjacent domains or within 70 kb, then the adjacent boundaries were adjusted to an average boundary.

Finally, we categorized domains into three layers: TADs, subTADs and ultra-nested subTADs. We defined ultra-nested subTADs as the innermost layer of domains[1, 11]. We defined TADs as the outermost layer[1, 11]. subTADs were classified as domains above the innermost and nested below the outermost layers[11,29] (**Supplementary Table 8**).

**Insulation score (IS) calculation**

We applied a 12 kb square window (12 x 12 bins on 10 kb binned matrices) with one bin offset from the diagonal on Knight-Ruiz balanced, scaled cis Hi-C maps on merged replicates of each genotype. To compute IS genome-wide, we summed counts in the 12 x 12 bin IS window, normalized by mean per chromosome, and log transformed the counts[10, 13]. IS values at the beginning of each chromosome corresponding to insufficient counts were discarded as NaN. We computed mean insulation score pileups at boundaries (+/- 400 kb around the center of each boundary) for each of the TADs, subTAD, and ultra-nested subTAD boundaries in WT, APP(Swe), and PSEN1(M146V) iPSC-neurons.

**Scalar normalization of ChIP-seq and ATAC-seq bigwig signals for each genotype in iPSC-derived neurons**

To normalize bigwig signal between WT and APP(Swe) for each H3K27ac, CTCF, RAD21 and RNA Polymerase II ChIP-seq and ATAC-seq libraries, we used pyBigwig (version 0.3.13) to get signal under each pileup interval in the bigwig file and we divided this signal by scalar size factors. We determined each size factor by finding the maximum global signal for each chromosome, computing the mean of those maximums, and then dividing that mean by minimum of WT and APP(Swe) pileup signal.

**Aggregate peak analysis (APA)**

We have previously described the details of running APA analysis in our previous works[1, 11], and they are again described in detail here to ensure reproducibility. Briefly, we extracted 130 kb x 130 kb squares from the scaled, 10 kb KR balanced contact matrices for each loop in WT, APP(Swe), and PSEN1(M146V) genotypes. We excluded loops smaller than 150 kb due to the high distance-dependent contact frequency at short genomic ranges. We computed the average interaction frequency in pixels within each looping class separately for each genotype to obtain the APA plots[10, 13].

**Subclassification of loops into Promoter-to-Promoter, Promoter-to-NonPromoter, and NonPromoter-to-NonPromoter**

We defined loop anchors as the region between start and end coordinates of the two sides of the rectangle enclosing all the bin-bin pairs in the looping cluster. We defined promoter regions as TSS±2kb of all hg38 RefSeq transcripts.

We classified WT-specific, APP(Swe)-specific, and genotype-invariant loops into:

(a) *Promoter-to-Promoter loops:* loops connecting anchors co-localized on both sides with Promoters of gene isoforms (promoters both sides)

(b) *Promoter-to-NonPromoter loops:* loops connecting anchors co-localized on one side with Promoters of gene isoforms and devoid of Promoters on the other anchor (promoter one side)

(c) *NonPromoter-to-NonPromoter loops:* loops connecting anchors devoid of any Promoters on both sides (no promoters on any side)

**Quantifying ATAC-seq signal at Promoter-to-NonPromoter and Promoter-to-Promoter loop anchors**

To create ATAC-seq MA plots for the Promoter loop anchors, we curated a list of genomic coordinates for TSS±2kb regions for unique gene isoforms anchoring WT-specific or APP-specific loops. We identified all unique, non-redundant ATAC-seq peaks in WT and APP(Swe) iPSC-derived neuron conditions co-localized with these genomic coordinates. For each intersecting ATAC-seq peak, we plotted mean of the scaled ATAC-seq bigwig signal in WT and APP(Swe) on the x-axis and log2 (APP(Swe)/WT) ATAC-seq signal on the y-axis.

To create ATAC-seq MA plots for the NonPromoter loop anchors, we curated a list of genomic coordinates for the full NonPromoter loop anchor. We identified all unique, non-redundant ATAC-seq peaks in WT and APP(Swe) iPSC-derived neuron conditions co-localized with these genomic coordinates. For each intersecting ATAC-seq peak, we plotted mean of the scaled ATAC-seq bigwig signal in WT and APP(Swe) on the x-axis and log2 (APP(Swe)/WT) ATAC-seq signal on the y-axis.

To create ATAC-seq MA plots for Promoters that do not loop, we curated a list of genomic coordinates for TSS±2kb regions for unique gene isoforms that do not engage in loops. We identified all unique, non-redundant ATAC-seq peaks in WT and APP(Swe) iPSC-derived neuron conditions co-localized with these genomic coordinates. For each intersecting ATAC-seq peak, we plotted mean of the scaled ATAC-seq bigwig signal in WT and APP(Swe) on the x-axis and log2 (APP(Swe)/WT) ATAC-seq signal on the y-axis.

**Stratifying unique gene isoforms into those anchoring WT-specific loops, APP(Swe)-specific loops, genotype-invariant loops, or not looping**

We made a genome-wide list of the unique, non-redundant TSS±2kb genomic coordinates representing all hg38 Refseq gene isoforms. We intersected all promoters with the anchors of WT-specific, APP(Swe)-specific, and genotype-invariant loops using the genomic coordinates of -10 kb of the start to + 10 kb after the end of both anchor 1 and anchor 2. Using BEDTools[23] (v2.29.1) intersect we identified all the gene isoforms whose TSS±2kb intersected the WT-specific, APP(Swe)-specific, and genotype-invariant anchors as follows:

**Class1**: Gene isoforms whose TSS±2kb overlaps at least one WT-specific loop anchor but not any APP(Swe)-specific anchor and can intersect any number of Invariant loop anchors.

**Class2**: Gene isoforms whose TSS±2kb region overlaps at least one APP(Swe)-specific loop anchor but not WT-specific loop anchor and can intersect any number of Invariant loop anchors.

**Class3**: Gene isoforms whose TSS±2kb overlap with only Invariant loop anchors.

**Class4**: All the remaining hg38 gene isoforms whose TSS±2kb does not overlap with loop anchors in our human iPSC-derived neuron Hi-C maps. We further removed gene isoforms from Class4 whose TSS±2kb fell in deadzone or low mappabiliy regions to ensure they were not due to overlap with unmapped Hi-C regions.

We plotted expression of only isoforms with DESeq2 MOR-normalized TPM counts>300 in at least one genotype in iPSC-derived neurons based on bulk-RNASeq (**Supplementary Table 12**).

**Defining H3K27ac+ enhancer regions in WT and APP(Swe) iPSC-neurons**

We identified putative cis regulatory elements genome-wide in WT and APP(Swe) using H3K27ac ChIP-seq peaks from each genotype in human iPSC-derived neurons. For WT neurons: Using BEDTools[23] (v2.29.1), we excluded any WT H3K27ac peaks that overlapped any TSS±2kb, Refseq exons, 5'UTRs, or 3'UTR regions. Thus, we identified only those WT H3K27ac peaks that were at introns and intergenic regions of the hg38 genome. For APP(Swe) neurons: we used APP(Swe) H3K27ac ChIP-seq peaks and excluded APP(Swe) H3K27ac peaks that overlapped any TSS±2kb, Refseq exons, 5'UTRs, or 3'UTR regions. We identified a total of 34,423 H3K27ac putative enhancer regions in WT and 33,673 H3K27ac putative enhancer regions in APP(Swe) human iPSC-derived neurons (**Supplementary Table 15**).

**Defining non-looping negative controls, including non-looping putative enhancers and non-looping promoters of gene isoforms with or without CTCF peaks**

For non-looping putative enhancers: we used BEDTools[23] (v2.29.1) subtract to remove all H3K27ac+ enhancers regions that overlapped either WT-specific, APP(Swe)-specific, or genotype-invariant loops in iPSC-derived neurons. We further used BEDTools[23] (v2.29.1) intersect to classify non-looping non-coding H3K27ac+ regions into two groups:

i) Non-looping putative enhancers co-localized with CTCF peaks and
ii) Non-looping putative enhancers devoid of CTCF occupancy

We used similar logic to classify non-looping gene isoforms into two groups:

i) Non-looping genes with a TSS±2kb co-localized with CTCF peaks and
ii) Non-looping genes with a TSS±2kb devoid of CTCF occupancy

**Subclassification of Promoter-to-Promoter and Promoter-to-NonPromoter loops by their co-localization with CTCF, non-coding H3K27ac, and TSS±2kb promoters**

We further subclassified Promoter-to-Promoter and Promoter-to-NonPromoter WT-specific, genotype-invariant, and APP(Swe)-specific loops into those that also co-localized with CTCF or non-coding H3K27ac+ peaks in WT or APP(Swe) iPSC-derived neurons.

We subclassified Promoter-to-NonPromoter loops as follows:

a. **<u>Promoter-to-NoncodingH3K27ac-positive(+) loops</u>**: loops connecting one anchor co-localized with Promoters of gene isoforms to a second anchor devoid of Promoters and containing non-coding or intronic H3K27ac peak (promoter one side and distal non-coding putative enhancer on the second side). We also further sub-classified each loop based on the presence or absence of CTCF occupancy at the loop anchors:
   i. **Promoter CTCF-to-Distal CTCF**
   ii. **Promoter CTCF-to-Distal NONE**
   iii. **Promoter NONE-to-Distal CTCF**
   iv. **Promoter NONE-to-Distal NONE**

b. **<u>Promoter-to-NoncodingH3K27ac-negative(-) loops</u>**: loops connecting one anchor co-localized with Promoters of gene isoforms to a second anchor devoid of Promoters or exons and also devoid of H3K27ac signal (promoter one side and no evidence for a regulatory element or coding region on the second side). We also further sub-classified each loop based on the presence or absence of CTCF occupancy at the loop anchors:
   i. **Promoter CTCF-to-Distal CTCF**
   ii. **Promoter CTCF-to-Distal NONE**
   iii. **Promoter NONE -to-Distal CTCF**
   iv. **Promoter NONE -to-Distal NONE**

Loops for each class formed by Class1 and Class2 transcript classes are included in (**Supplementary Table 11**).

**Computing burst size and burst frequency from single-cell RNA-seq data**

We analyzed the single-cell RNAseq libraries generated from 10x Genomics v3 (3′ scRNaseq) method. We used 10x Genomics Cell Ranger[30] (v6.1.2) ‘cellranger mkfastq’ and ‘cellranger count’ to align fastq reads to the GRCh38 human genome with default settings. After alignment, the Cell Ranger filtered matrix files contained 7070 cells in WT and 16279 cells in APP(Swe) from iPSC-derived neurons. We then downsampled WT cells using downsampleMatrix with prop=0.8 parameter to match the sequencing read depth of APP(Swe). Next, we merged WT and APP(Swe) matrices and loaded them into Seurat[31] (v4.1.0) using ‘CreateSeuratObject’ function with min.cells = 1100, min.features = 800. These parameters ensured that only cells with at least 800 detected genes and genes that were detected in at least 1100 cells across WT and APP(Swe) genotypes were retained. We further applied nFeature_RNA < 3000 and mitochondrial percentage < 10 for doublet removal and quality-control filtering. These filters resulted in 5544 cells in WT, 13366 cells APP and 7160 total genes detected. We downsampled APP(Swe) to 5544 cells to match WT condition and applied Relative count normalization method in NormalizeData by specifying normalization.method ='RC' and scale.factor = 10000. Final dimensions of the normalized datasets were 5544 cells and 7160 genes for each of the WT and APP(Swe) genotypes.

We computed burst size and burst frequency consistent with the approach previously published[32]. Briefly, from the normalized single-cell RNA-seq data across 5544 cells and 7160 genes in WT, we computed the mean (μ) and variance (σ2) for each gene by sampling 50 cells from each distribution 1000 times. We next computed burst size as $bs = \sigma 2/\mu$ and burst frequency as $bf = \mu/(bs-1)$. Similarly, we repeated the analysis on the normalized counts from APP(Swe) (**Supplementary table 13**).

**Motif analysis at the distal non-coding H3K27ac+ ATAC+ and promoter regions of WT-specific (Class 1) and APP(Swe)-specific (Class 2) loops**

We used MEME AME suite (version 5.1.1) and the JASPAR2022_CORE_vertebrates_non-redundant.meme database to perform motif enrichment analysis. We used the following parameters in ame: --scoring avg --method fisher --evalue-report-threshold 10.0.

<u>For WT-specific (Class 1) Promoter-to-NoncodingH3K27ac-positive(+) loops:</u>

<u>> At distal noncoding anchors:</u>

For each of the 4 CTCF loop classes, we intersected each distal non-coding H3K27ac+ loop anchor with ATAC-seq peaks that also co-localized with the WT H3K27ac+ enhancer peak. If the distal anchor intersected with only one ATAC-seq peak, that peak was used. If the distal anchor intersected multiple ATAC-seq peaks at the H3K27ac+ enhancer peak, then the widest ATAC-seq peak was used. Thus, each distal non-coding H3K27ac+ anchor was represented by one ATAC-seq peak for downstream motif analysis.

<u>For APP(Swe)-specific (class 2) Promoter-to-NoncodingH3K27ac-positive(+) loops:</u>

<u>> At distal noncoding anchors:</u>

For each of the 4 CTCF loop classes, we intersected each distal non-coding H3K27ac+ loop anchor with ATAC-seq peaks that also co-localized with the APP(Swe) H3K27ac+ enhancer peak. If the distal anchor intersected with only one ATAC-seq peak, that peak was used. If the distal anchor intersected multiple ATAC-seq peaks at the H3K27ac+ enhancer peak, then the widest ATAC-seq peak was used. Thus, each distal non-coding H3K27ac+ anchor was represented by one) ATAC-seq peak for downstream motif analysis.

For H3K27ac, RNA Polymerase II, CTCF or RAD21 ChIP-seq and ATAC-seq heatmaps at promoter and distal loop anchors, we plotted the heatmaps using deeptools package (version 3.5.3). For promoter anchors, heatmaps were plotted around the +/-5kb region upstream and downstream of each TSS and TES respectively. For the distal non-coding anchors, we plotted heatmaps that were +/- 100kb centered around the H3K27ac, CTCF, RNA Polymerase II, RAD21 or ATAC-seq peaks that intersected the distal non-coding loop anchors.

<u>As controls for the motif analyses, we created 4 background control regions:</u>
For distal non-coding regions:

1. Non-looping putative enhancers co-localized with CTCF peaks
2. Non-looping putative enhancers devoid of CTCF occupancy

For non-looping gene isoform promoters:

3. Non-looping genes with a TSS-2kb co-localized with CTCF peaks and
4. Non-looping genes with a TSS-2kb devoid of CTCF occupancy

We performed motifs analysis by downsampling controls to match numbers of the test peaks.
> For motif analysis at the distal non-coding H3K27ac+ ATAC+ peaks derived from either Promoter CTCF-to-Distal CTCF or Promoter NONE-to-Distal CTCF loop anchors: we used non-looping putative enhancers co-localized with CTCF peaks as negative controls.
> For motif analysis at the distal non-coding H3K27ac+ ATAC+ peaks derived from either Promoter CTCF-to-Distal None or Promoter None-to-Distal None loop anchors: we used non-looping putative enhancers devoid of CTCF occupancy as negative controls.
> For motif analysis at the -2kb TSS region derived from either Promoter CTCF-to-Distal CTCF or Promoter CTCF-to-Distal None loop anchors: we used non-looping -2kb TSS regions co-localized with CTCF peaks as negative controls.
> For motif analysis at the -2kb TSS region derived from either Promoter None-to-Distal CTCF or Promoter None-to-Distal None loop anchors: we used non-looping -2kb TSS regions devoid of CTCF occupancy as negative controls.

**Re-analysis of published DLX1, DLX5 ChIPseq from mouse brain tissue**

We downloaded published DLX1, DLX5 ChIP-seq fastq files from Lindtner et al. 2019[33] in mouse E16.5 Basal ganglia from GEO (GSM3559636, GSM3559637, GSM3559645). We mapped the ChIP-seq reads to mm10 reference genome using Bowtie[19] (version 0.12.7). We removed reads with more than two possible alignments using the -m2 flag. We filtered the mapped reads to remove optical and PCR duplicates using samtools[20] (version 1.2) and down-sampled the ChIP-seq libraries to 19 million reads to achieve equal read numbers across samples. We identified peaks using MACS2[21] (version 2.1.2) with a p-value cutoff parameter of 1E-4 using default MACS2 parameters. We converted bigwigs and peak calls from mm10 to hg38 using UCSC tools *liftOver.* We used the ChIP-seq signal from the hg38 converted bigwig files for mouse DLX1, DLX5 E16.5 brain tissue samples for the analysis/figures.

We also obtained DLX1, DLX5 motifs from JASPAR (https://ccg.epfl.ch//pwmtools/pwmscan.php; JASPAR Core 2020 vertebrate library, motif: DLX1 MA0879.1; DLX5 MA1476.1; p-value 1e-3;).

**Hi-C and loop calling after transduction of D3 i3N-neurons with lentivirus encoding DLX1 or mCherry transgenes**

We created 20kb cis-interaction Hi-C heatmaps for i3Ns engineered with lentivirus to overexpress mCherry, or DLX1. To identify chromatin loops genome-wide, we applied the same methods as described above with minor modifications. After assembling raw 20kb binned Hi-C cis matrices, we implemented a 1D distance dependence median-of-ratios normalization to correct for sequencing depth differences across all 3 conditions. We balanced matrices with the Knight-Ruiz methodology as described above.

We computed a donut expected model using the parameters p=4 and w=10 and identified loop clusters at q-value 0.2. We identified 2,318 loops in mCherry+ i3Ns, and 984 loops in DLX1+ i3Ns. For these difficult experiments, we did not have extensive replicates needed to model loop interaction frequency variance. Therefore, we pursued a simple approach to identify condition-specific loops between mCherry and DLX1+ overexpression. We first created a merged non-redundant list of loop anchors from both the conditions. Using BEDTools[23] (v2.29.1), we classified loops as mCherry+-specific if either one of upstream or downstream loop anchors did not overlap with the location of loop anchors in the DLX1+ condition. Similarly, loops we classified loops as DLX1+-specific loops if at least one of the loop anchors did not overlap with any loop anchors in the mCherry+ condition. We found 1,590 mCherry+-specific loops, 277 DLX1+-specific loops and 727 invariant loops between mCherry+ and DLX1+ conditions.

**Classification of mCherry+-specific, DLX1+-specific and invariant loops into Promoter-Promoter, Promoter-distal loops (Related to Figure 5k)**

We created a concatenated the list of unique, non-redundant mDLX1, mDLX5 peaks after *liftOver* to hg38 genome. We identified only those mDLX1/5 peaks at hg38 introns or intergenic regions (noncoding DLX1/5 peaks). Briefly, using BEDTools[23] (v2.29.1), we excluded any mDLX1, mDLX5 peaks that overlapped any TSS±2kb, Refseq exons, 5'UTRs, or 3'UTR regions. We intersected the noncoding DLX1/5 peaks with DLX1 motifs and counted the number of motifs overlapping each peak. As defined previously, for promoter regions, we used the genome-wide list of unique, non-redundant TSS±2kb genomic coordinates representing all hg38 Refseq gene isoforms. We created 3 lists of noncoding DLX1/5 peaks with 0, 1, >=2 DLX1 motifs. We classified each mCherry+-specific, DLX1+-specific and invariant loops "*Promoter-to-distal noncoding loops amenable to DLX binding*" if: one anchor of the loop overlaps a defined promoter region and the other non-promoter anchor overlapped a noncoding DLX1/5 peak with either >=2 or 1 or 0 DLX1 (TAATTA) motifs. We plotted the number of mCherry-specific, DLX1+-specific and invariant loops that were thus classified as *Promoter-to-distal noncoding loops amenable to DLX binding* whose distal non-promoter anchor with noncoding DLX1/5 overlapped >=2 TAATTA motifs, or 1 TAATTA motif or 0 motifs.

**Defining H3K27ac+ enhancer regions in D3 i3Ns after overexpression (Figure 5m)**

We utilized published data from Sanchez-Priego et al. 2022[34] D35 i3Ns that was closely similar to our D3 i3N experimental conditions. We downloaded the provided H3K27ac CUT&RUN bigwig and peak call files for 3 replicates at T0 timepoint in glutamatergic neurons mapped to hg38 genome from GEO (GSM5862749, GSM5862753, GSM5862756). We created a concatenated list of unique, non-redundant peaks from the 3 replicates. Using BEDTools[23] (v2.29.1), we excluded any H3K27ac peaks that overlapped any TSS±2kb, Refseq exons, 5'UTRs, or 3'UTR regions and thus identified only those H3K27ac peaks that overlapped introns and intergenic regions. We merged the 3 H3K27ac bigwig replicates and utilized this merged bigwig for plotting.

**Clustering analysis of single-cell RNA-seq data in WT and APP(Swe) iPSC-derived neurons**

We analyzed the single-cell RNAseq libraries generated from 10x Genomics v3 (3′ scRNaseq) method. We used 10x Genomics Cell Ranger[30] (v6.1.2) 'cellranger mkfastq' and 'cellranger count' to align fastq reads to the GRCh38 human genome with default settings. After alignment, the Cell Ranger filtered matrix files contained 7070 cells in WT and 16279 cells in

APP(Swe) from iPSC-derived neurons. We then downsampled WT cells using downsampleMatrix with prop=0.8 parameter to match the sequencing read depth of APP(Swe). Next, we merged WT and APP(Swe) matrices and loaded them into Seurat[31] (v4.1.0) using 'CreateSeuratObject' function with min.cells = 1100, min.features = 800. These parameters ensured that only cells with at least 800 detected genes and genes that were detected in at least 1100 cells across WT and APP(Swe) genotypes were retained. We further applied nFeature_RNA < 3000 and mitochondrial percentage < 10 for doublet removal and quality-control filtering. We downsampled APP(Swe) to match WT condition and applied Log Normalize method in NormalizeData by specifying normalization.method = LogNormalize and scale.factor = 10000. We next merged the resulting normalized WT and APP(Swe) Seurat objects using `merge` function. We next applied `FindVariableFeatures` with selection.method='vst' to identify highly variable features. We applied a linear transformation to scale the data for all the genes using `ScaleData` function. Next, we performed PCA on the scaled data on using `RunPCA` using all features. After applying `FindNeighbors()` with dims=1:5 , we finally clustered the merged WT and APP(Swe) Seurat object using `FindClusters` with resolution = 0.35 and this resulted in a total of 9 clusters. We applied non-linear dimensional reduction UMAP for visualization using `RunUMAP`. We identified markers that define each clusters using `FindMarkers` with parameters only.pos = TRUE, min.pct = 0.25, logfc.threshold = 0.25 and annotated the clusters according to the markers identified. Finally, we used `DimPlot`, `DoHeatmap` and `FeaturePlot` functions in Seurat for different visualizations of 5,539 cells and n=7,153 features in WT and APP(Swe) on the figures.
